# Structural basis of PETISCO complex assembly during piRNA biogenesis in *C. elegans*

**DOI:** 10.1101/2021.05.20.444926

**Authors:** Cecilia Perez-Borrajero, Nadezda Podvalnaya, Kay Holleis, Raffael Lichtenberger, Emil Karaulanov, Bernd Simon, Jérôme Basquin, Janosch Hennig, René F. Ketting, Sebastian Falk

**Affiliations:** Structural and Computational Biology Unit, EMBL Heidelberg, Meyerhofstraße 1, 69117, Heidelberg, Germany; Biology of Non-coding RNA Group, Institute of Molecular Biology, Mainz, Germany; International PhD Programme on Gene Regulation, Epigenetics & Genome Stability, Mainz, Germany; Department of Structural and Computational Biology, Max Perutz Labs, University of Vienna, 1030 Vienna, Austria; Bioinformatics Core Facility, Institute of Molecular Biology, Mainz, Germany; Max-Planck-Institute of Biochemistry, Department of Structural Cell Biology, Am Klopferspitz 18, 82152 Martinsried, Germany; Chair of Biochemistry IV, Biophysical Chemistry, University of Bayreuth, Universitätsstrasse 30, 95447 Bayreuth, Germany; Institute of Developmental Biology and Neurobiology, Johannes Gutenberg University, 55099 Mainz, Germany

**Keywords:** piRNA, 21U RNA, Piwi, C. elegans, PETISCO, RRM domain, ERH-2, IFE-3, TOFU-6, PID-3, TOST-1, PID-1

## Abstract

Piwi-interacting RNAs (piRNAs) constitute a class of small RNAs that bind PIWI proteins and are essential to repress transposable elements in the animal germline, thereby promoting genome stability and maintaining fertility. *C. elegans* piRNAs (21U RNAs) are transcribed individually from minigenes as precursors that require 5’ and 3’ processing. This process depends on the PETISCO complex, consisting of four proteins: IFE-3, TOFU-6, PID-3, and ERH-2. We employ biochemical and structural biology approaches to characterize the PETISCO architecture and its interaction with RNA, together with its effector proteins TOST-1 and PID-1. These two proteins define different PETISCO functions: PID-1 governs 21U processing whereas TOST-1 links PETISCO to an unknown process essential for early embryogenesis.

Here, we show that PETISCO forms an octameric assembly with each subunit present in two copies. Determination of structures of the TOFU-6/PID-3 and PID-3/ERH-2 subcomplexes, supported by *in vivo* studies of subunit interaction mutants, allows us to propose a model for the formation of the TOFU-6/PID-3/ERH-2 core complex, and its functionality in germ cells and early embryos. Using NMR spectroscopy, we demonstrate that TOST-1 and PID-1 bind to a common surface on ERH-2, located opposite its PID-3 binding site, explaining how PETISCO can mediate different cellular roles.

## Introduction

RNA molecules typically require processing after transcription before becoming fully functional. In eukaryotes, messenger RNAs (mRNAs) and many non-coding RNAs are spliced, capped, and poly-adenylated by processing factors (Hocine et al. 2010). Transcripts can also be chemically modified, trimmed or cleaved by ribonucleases, or extended by the addition of non-templated nucleotides (Roundtree et al. 2017; Yu and Kim 2020). Such processing steps are crucial for activation, (de)stabilization, localization, and many other aspects relevant to RNA function.

Piwi-interacting RNAs (piRNAs) constitute one of the largest classes of non-coding RNA transcripts whose processing is only starting to be understood (Weick and Miska 2014; Ozata et al. 2019). piRNAs play a crucial role in the germline, where they act as specificity factors in genome defense pathways with transposable elements as major targets (Luteijn and Ketting 2013; Czech and Hannon 2016). The proteins guided by piRNAs, Piwi proteins, are an animal-specific subgroup of the Argonaute family. Piwi proteins are guided by piRNAs that bind to their target site, leading to either transcript cleavage or modification of chromatin, depending on the subclass of Piwi proteins involved (Luteijn and Ketting 2013). In either case, the sequence of the piRNAs bound by the Piwi proteins dictates the target specificity of the silencing process and is therefore crucial for function. Hence, the mechanism(s) that act in piRNA precursor selection and processing determine the specificity of Piwi proteins.

The precursors of piRNAs are single-stranded RNA transcripts. In *Caenorhabditis elegans* (*C. elegans*), these are produced from a multitude of miniature genes, each producing a small transcript of 27-30 nucleotides (Ruby et al. 2006; Gu et al. 2012). Mature piRNAs are bound by the Piwi protein PRG-1 and are typically 21 nucleotides long with an uracil base at the 5′ end. For these reasons, the piRNAs in *C. elegans* are often named 21U RNAs. To form mature 21U RNAs, the precursor transcripts are shortened at both ends, including the removal of the 5’ cap and trimming of the 3’ end (Ruby et al. 2006; Wang and Reinke 2008; Batista et al. 2008; Das et al. 2008; Gu et al. 2012; Weick and Miska 2014; Tang et al. 2016). The vast majority of 21U RNAs stem from dedicated loci characterized by a specific sequence motif in their promoter termed the Ruby motif (Ruby et al. 2006; Cecere et al. 2012; Weick et al. 2014). Within the *C. elegans* genome, these loci are strongly clustered, suggesting that they may act in concert (Ruby et al. 2006). The transcription of these loci, including the termination of their transcription, bears hallmarks of small nuclear RNA (snRNA) biogenesis, suggesting that the *C. elegans* piRNA system has evolutionary connections to these non-coding snRNAs that play essential roles in splicing (Kasper et al. 2014; Beltran et al. 2019; Weng et al. 2019; Beltran et al. 2020; Berkyurek et al. 2021).

Following the genetic identification of several 21U RNA processing genes (Goh et al. 2014; Albuquerque et al. 2014), we and others identified a four-member protein complex required for 21U RNA biogenesis, which we named PETISCO (PID-3, ERH-2, TOFU-6, IFE-3 small RNA complex) (Cordeiro Rodrigues et al. 2019; Zeng et al. 2019). PETISCO interacts with and stabilizes 21U RNA precursors. Interestingly, PETISCO was shown to be additionally required for early embryogenesis, a function independent of 21U RNA biogenesis (Cordeiro Rodrigues et al. 2019; Zeng et al. 2019). In this case, loss of PETISCO function leads to a so-called ‘maternal effect lethal’ (Mel) phenotype, in which first-generation homozygous mutant animals develop normally, but their offspring arrest in embryogenesis. At the molecular level, low levels of the splice-leader transcript SL1, a small nuclear RNA (snRNA) involved in trans-splicing, were found to be bound by PETISCO. SL1-derived 21U RNAs have also been described, albeit at very low levels (Gu et al. 2012). Whether these findings relate to the Mel phenotype of PETISCO mutants is unclear, but they do strengthen the link between 21U RNAs and snRNAs in *C. elegans*. The two described functions of PETISCO are specified by two different effector proteins, PID-1 and TOST-1. PID-1:PETISCO mediates 21U RNA biogenesis (Albuquerque et al. 2014), while TOST-1:PETISCO is required for early embryogenesis (Cordeiro Rodrigues et al. 2019; Zeng et al. 2019).

The PETISCO subunits contain domains often present in RNA-binding proteins (Cordeiro Rodrigues et al. 2019). PID-3 and TOFU-6 are restricted to the nematode phylum and contain two domains. PID-3 has a predicted RNA-recognition (RRM) domain and an Argonaute-related middle (MID) domain. TOFU-6 contains an RRM domain, an extended Tudor (eTudor) domain, and a C-terminal eIF4E interaction motif. IFE-3 is one of the five highly conserved *C. elegans*’ eIF4E homologs (Keiper et al. 2000), and binds to the C-terminus of TOFU-6 (Cordeiro Rodrigues et al. 2019). Finally, ERH-2 is one of the two *C. elegans* paralogs of ‘enhancer of rudimentary’ (Erh), a factor that is conserved throughout eukaryotes (Weng and Luo 2013). Erh was shown to participate in the RNA exosome-mediated degradation of meiotic RNAs in *Schizosaccharomyces pombe* (*S. pombe*) (Sugiyama et al. 2016) and to facilitate miRNA processing in human cells (Fang and Bartel 2020; Hutter et al. 2020; Kwon et al. 2020).

An approximate architecture of PETISCO was previously derived from yeast two-hybrid (Y2H) studies (Fig. 1A) (Cordeiro Rodrigues et al. 2019). However, the structural basis of PETISCO assembly and its interaction with the effector proteins PID-1/TOST1 and RNA substrates remain poorly understood, limiting our understanding of PETISCO function. Here, we study PETISCO assembly using a bottom-up approach with purified proteins, interaction studies, and structural analyses. We find that PETISCO forms a dimer of tetramers, in which dimerization is mediated both by PID-3 and ERH-2. Crystal structures of the PID-3/TOFU-6 and ERH-2/PID-3 subcomplexes reveal insights into PETISCO assembly, function, and subcellular localization. Using NMR spectroscopy, we also characterize the mutually exclusive interplay of ERH-2 with the two effector proteins TOST-1 and PID-1. These results represent the first structural characterization of a piRNA biogenesis complex, and we start to reveal how PETISCO may execute its dual role *in vivo*.

**Figure 1.**
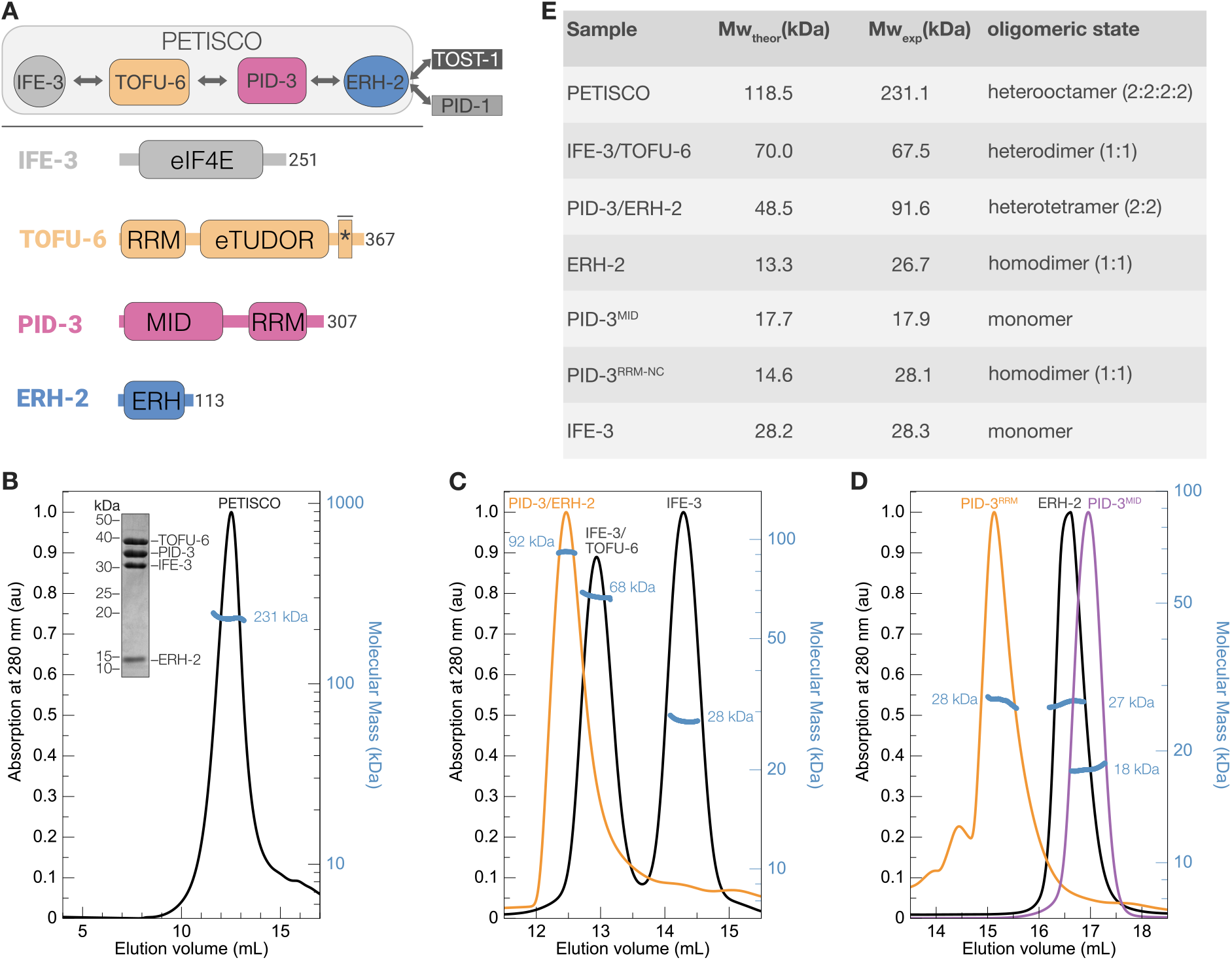
PETISCO assembles into a hetero-octamer with 2:2:2:2 stoichiometry. (A) Top: schematic representation of the PETISCO topology and subunit interactions and binding of the effector proteins TOST-1 and PID-1. Bottom: domain organization of IFE-3, TOFU-6, PID-3 and ERH-2 from *C. elegan*s. Rounded rectangles indicate predicted domains. The asterisk marks the position of the IF4E interaction motif that mediates binding of TOFU-6 to IFE-3. (B-D) SEC-MALS chromatograms showing UV absorption at 280 nm and the calculated molecular mass in kilodaltons (kDa). The UV absorption signal was normalized to the highest peak. (B) SEC-MALS profile of PETISCO. The inset shows a Coomassie-stained SDS polyacrylamide gel of PETISCO after SEC. (C) SEC-MALS profiles of the IFE-3/TOFU-6 and IFE-3 (black line), and PID-3/ERH-2 (orange line). (D) SEC-MALS profiles of ERH-2 (black line), PID-3^MID^ (purple line) and PID-3^RRM^ (orange line) domains. (E) Summary of molecular masses and stoichiometries of PETISCO and its subunits.

## Results

### PETISCO forms an octameric assembly

PETISCO consists of the proteins IFE-3, TOFU-6, PID-3, and ERH-2. Previous Y2H experiments (Cordeiro Rodrigues et al. 2019) revealed a linear topology in which IFE-3 binds to TOFU-6, which in turn binds PID-3, which associates with ERH-2 (Fig. 1A, Supplemental Fig. 1A). To analyze the oligomeric state and stoichiometry of PETISCO, we recombinantly expressed and purified PETISCO components from bacterial cells and subjected the complex to size-exclusion chromatography (SEC) coupled to multiangle light scattering (MALS) (Fig. 1B). The complete PETISCO complex eluted as a single peak and showed an average molecular mass of 231 kDa as determined by SEC-MALS (Fig. 1B), twice the sum of the individual components (assuming a 1:1:1:1 complex, 116 kDa), suggesting that PETISCO forms a hetero-octameric assembly, a dimer of tetramers, with 2:2:2:2 stoichiometry (Fig. 1B). Next, we set out to determine which protein or domains mediate oligomerization. The linear topology of PETISCO suggested that the complex could be divided into IFE-3/TOFU-6 and PID-3/ERH-2 subcomplexes (Fig. 1A). We individually purified these and analyzed their molecular mass by SEC-MALS (Fig. 1C). In the case of IFE-3/TOFU-6, we measured an average mass of 69 kDa, in line with the calculated mass of a heterodimer. The PID-3/ERH-2 subcomplex, however, had an average mass of 92 kDa, consistent with that of a hetero-tetramer, suggesting that the PID-3/ERH-2 module mediates oligomerization. Since the human and fission yeast ERH orthologs have been shown to form homodimers (Wan et al. 2005; Xie et al. 2019; Hazra et al. 2020), we hypothesized that ERH-2 is responsible for dimerization. Therefore, we determined the oligomeric state of both ERH-2 and PID-3 separately. However, PID-3 full-length protein could not be prepared in a quality suitable for SEC-MALS, and therefore we analyzed the PID-3 MID and RRM domains individually, denoted here as PID-3^MID^ and PID-3^RRM^, respectively. We determined average masses of 18, 28, and 27 kDa for PID-3^MID^, PID-3^RRM-NC^, and full-length ERH-2^FL^, respectively (Fig. 1D,E and Supplemental Figure S1C). This suggests that PID-3^MID^ is monomeric, whereas both PID-3^RRM^ and ERH-2 are homodimers (Fig. 1E). We thus concluded that both PID-3^RRM^ and ERH-2 contribute to PID-3/ERH-2 subcomplex dimerization, and through binding of two IFE-3/TOFU-6 subcomplexes, this results in the formation of the octameric PETISCO.

### Crystal structure of the TOFU-6–PID-3 RRM core complex

The topological arrangement of PETISCO places TOFU-6 and PID-3 at the core, and we thus proceeded to narrow down their interacting regions. Our previous experiments indicated that the interaction between TOFU-6 and PID-3 is mediated by the RRM domains (denoted as TOFU-6^RRM^ and PID-3^RRM^) (Fig. 1A, Supplemental Fig. S1A) (Cordeiro Rodrigues et al. 2019). To better map which domains mediate the interaction between TOFU-6 and PID-3, we used a combination of pull-down experiments and SEC. We recombinantly co-expressed maltose binding protein (MBP)-tagged TOFU-6 constructs with glutathione-S-transferase (GST)- tagged PID-3 constructs. Both MBP and GST pull-downs revealed that the RRM domains of PID-3 and TOFU-6 mediate the interaction between the two proteins and that neither the eTudor domain of TOFU-6 (TOFU-6^eTUDOR^) nor the PID-3^MID^ are required (Supplemental Fig. S2A). This is supported by SEC using purified TOFU-6^RRM^ and PID-3^RRM^ proteins (Supplemental Fig. S2B).

To gain structural insights into the TOFU-6^RRM^/PID-3^RRM^ complex, we determined the structures of PID-3^RRM^ and the TOFU-6^RRM^/PID-3^RRM^ complex, at 1.8 Å and 1.7 Å resolutions, respectively (Fig. 2A and Supplemental Fig. S2C, Supplemental Table S1). Both structures are very similar. The PID-3^RRM^ forms a homodimer in both cases, with slight differences in the relative orientation of the RRM domains. (Supplemental Fig. S2C). Here, we will focus on the analysis of the TOFU-6^RRM^/PID-3^RRM^ complex structure, containing one tetrameric TOFU-6^RRM^/PID-3^RRM^ complex in the asymmetric unit (Fig. 2A), consistent with the SEC-MALS analysis (Supplemental Fig. S2D). The PID-3^RRM^ adopts a canonical RRM fold with an antiparallel four-stranded β-sheet packing opposite two α-helices (α1 and α2) (Supplemental Fig. S2E). The TOFU-6^RRM^ has a similar architecture but contains an additional fifth β-strand β4* located between the α2 and β4 elements (Supplemental Fig. S2F). The PID-3^RRM^ dimerizes via the α1 helix by a combination of hydrophobic and polar interactions (Fig. 2A and 2B) reminiscent, for instance, of the dimer interface in HuR-RRM3 (Pabis et al. 2019; Ripin et al. 2019) and RBPMS (Teplova et al. 2016). Phe217 from one PID-3^RRM^ protomer packs in a hydrophobic pocket created by Phe217, Ala220, Val228, and Ile231 from the other protomer. Moreover, Gln221 forms a hydrogen bond with the polypeptide backbone of the neighboring protomer (Fig. 2B).

**Figure 2.**
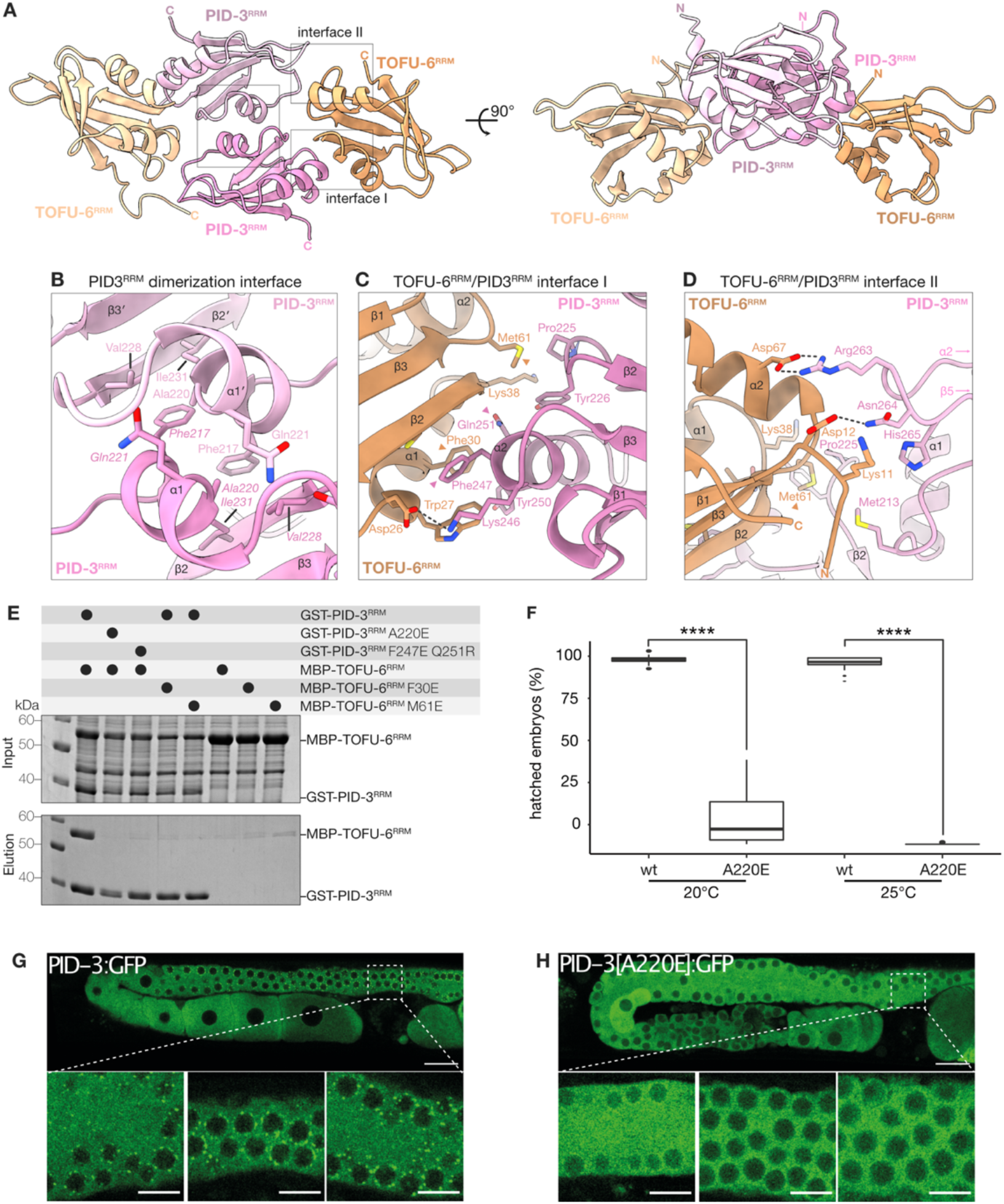
The TOFU-6 and PID-3 RRM domains form the PETISCO core. (A) Crystal structure of the PID-3^RRM^/TOFU-6^RRM^ complex shown in cartoon representation in two orientations related by a 90° rotation about the horizontal axis. The two PID-3^RRM^ protomers are shown in different shades of pink, while the two TOFU-6^RRM^ protomers in different shades of orange. The N- and C-terminal residues are highlighted. (B) Zoomed-in view of the homodimerization interface of PID-3^RRM^. Interacting residues are shown in stick representation and labeled. (C, D) Zoomed-in view of a representative set of residues at the PID-3/TOFU-6 interaction interface I (C) and interface II (D). Interacting residues are shown in stick representation as indicated. (E) Analysis of the effect of structure-guided mutations on the PID-3/TOFU-6 interaction by GST pull-down assays. Wild-type (WT) and mutant versions of MBP-tagged TOFU-6 constructed were co-expressed with WT and mutant versions of GST-tagged PID-3 constructs. Input and elution fractions were analyzed by SDS-PAGE followed by Coomassie staining. (F) Box-plot showing the percentage of hatched embryos of *pid-3::gfp(wt)* and *pid-3[a220e]::gfp* animals grown at 20 and 25°C. Progeny of 30 different mothers were analyzed for each condition, and development of at least 2600 eggs was scored. A two-sided T-test was used to assess significance. P-values are indicated in the graph. (G,H) Single plane confocal micrographs of PID-3::GFP(WT) and PID-3[A220E]::GFP at 25°C. The boxes indicate the regions (above the spermatheca) from which three zoomed-in examples are shown below. Scale bars: 20 µm in overview, 8 µm in zoom-in.

Two TOFU-6 RRM domains bind on either side of the interfaces created through the PID-3^RRM^ dimerization and contact both PID-3 protomers (Fig. 2A). The TOFU-6 α1 helix, β2 strand, and β3-α2 loop form a surface that interacts with the α1-β2 loop and α2 helix from the PID-3^RRM^ (Interface I/protomer 1) by a combination of hydrophobic and polar interactions (Fig. 2C). The TOFU-6^RRM^ interaction with the second PID-3^RRM^ protomer involves electrostatic interactions between the N-terminus and the α2 helix of TOFU-6^RRM^ with the PID-3^RRM^ α2-β5 loop (Interface II/protomer 2), featuring a salt bridge between TOFU-6 Asp67 and PID-3 Arg263, as well as a hydrogen bond between TOFU-6 Asp12 and PID-3 Asn264 (Fig. 2D).

Our structural analysis suggests that dimerization of the PID-3^RRM^ is a prerequisite for TOFU-6^RRM^ binding and that two interfaces at the two protomers contribute to the interaction. To test this prediction, we engineered a series of substitutions both at the PID-3 dimerization interface (PID-3^RRM^ A220E) and at interface I (TOFU-6^RRM^ F30E and M61E; PID-3^RRM^ F247E/Q251R) and analyzed their impact on the interaction by bacterial co-expression pull-down experiments. Both PID-3 mutant proteins (A220E and F247E/Q251R) are folded (Supplemental Figure S2G), however, the A220E mutation rendered PID-3^RRM^ monomeric, while the F247E/Q251R mutant remained homodimeric as expected from the structural analysis (Supplemental Fig. S2H-I). Mutations at the PID-3^RRM^ dimerization surface and interface I disrupted TOFU-6 binding, confirming that these sites contribute to a stable association with TOFU-6^RRM^ (Fig. 2E).

Next, we tested the effect of the monomer-inducing A220E mutation in PID-3 *in vivo*. Using CRISPR-Cas9 mediated gene editing, we first created a strain expressing C-terminally GFP-tagged PID-3, such that *in vivo* expression could be monitored by fluorescence microscopy. We then introduced the A220E mutation and scored its effect on subcellular localization and embryonic viability. *pid-3(a220e)* animals showed a strong Mel phenotype, which was fully penetrant at 25°C (Fig. 2F), consistent with loss of PETISCO function. In addition, PID-3(A220E) did not form peri-nuclear foci (Fig. 2G,H), indicating that PID-3 dimerization is required for the previously described P granule localization of PETISCO (Cordeiro Rodrigues et al. 2019; Zeng et al. 2019).

### ERH-2 binds to a region upstream of the PID-3 RRM domain

Next, we investigated the interaction between PID-3 and ERH-2. ERH-2 consists of an enhancer of rudimentary (ERH) domain followed by a C-terminal region of ∼ 15 amino acids (Fig. 1A). Previous experiments (Zeng et al. 2019; Cordeiro Rodrigues et al. 2019) suggested a direct interaction between ERH-2 and PID-3, in particular between PID-3^RRM^ and ERH-2 (Cordeiro Rodrigues et al. 2019). However, the PID-3 constructs used in the latter study contained additional N-terminal and C-terminal regions flanking the PID-3^RRM^ (Supplemental Fig. 1A). To better define the ERH-2 binding site of PID-3, we used PID-3 constructs covering the RRM domain (PID-3^RRM^) or the RRM domain with an N-terminal extension (PID-3^RRM-N^) and performed bacterial co-expression pull-down experiments. While PID-3^RRM-N^ pulled down ERH-2, PID-3^RRM^ failed to bind ERH-2 (Fig. 3A). To rule out that the C-terminal extension downstream of the PID-3 RRM domain contributed to binding, we performed SEC experiments. We incubated PID-3^RRM-N^ and PID-3^RRM-C^ with ERH-2 and analyzed the mixtures by SEC to assess binding. This confirmed that PID-3^RRM-N^ did, but PID-3^RRM-C^ did not interact with ERH-2 (Supplemental Fig. S3A,B). We then asked if the N-terminal extension of the PID-3 RRM alone was sufficient for ERH-2 binding. We purified a GST-tagged PID-3 peptide (PID-3^pep^, residues 171-203) corresponding to this region and found that ERH-2 could indeed bind to GST-PID-3^pep^ (Fig. 3A).

**Figure 3.**
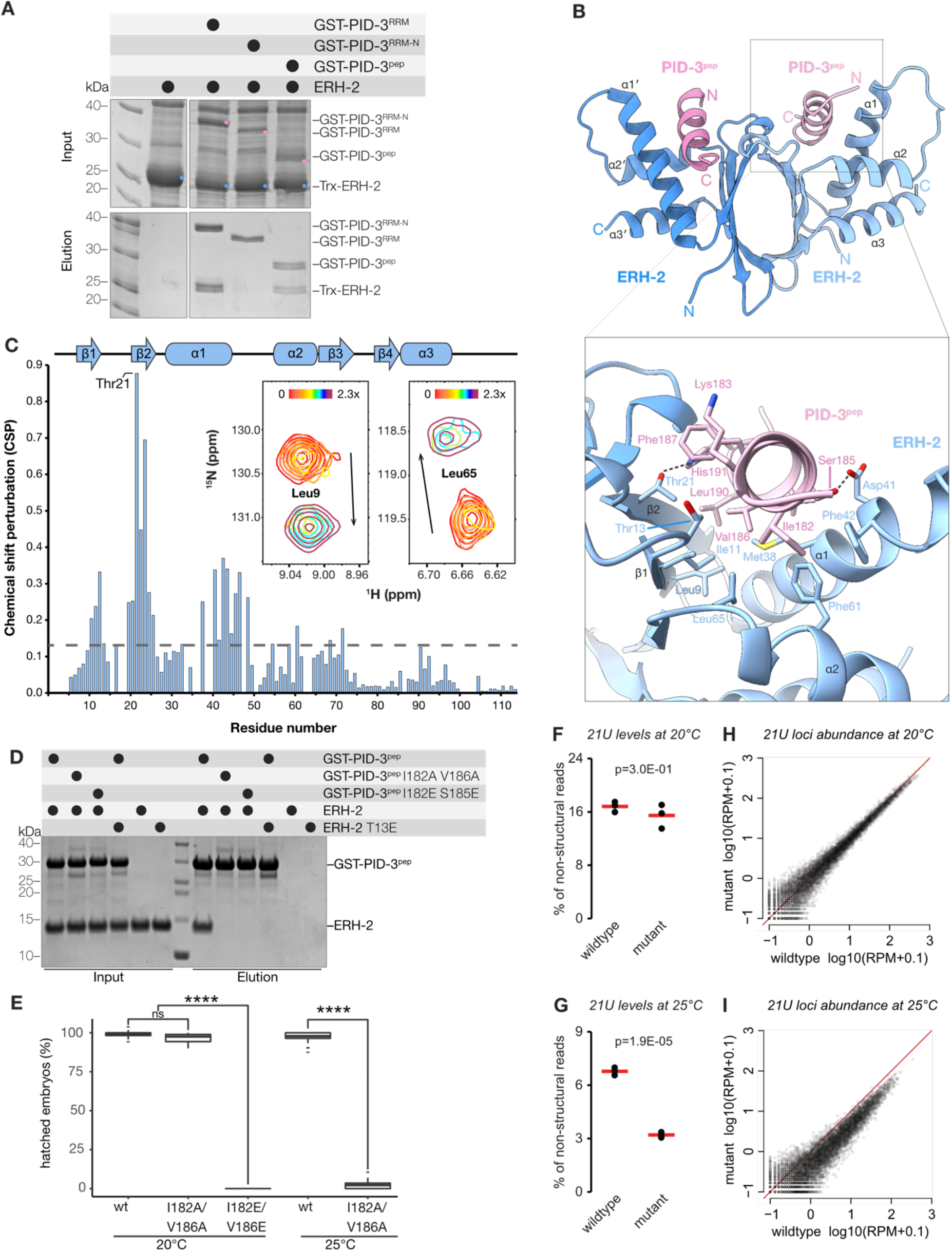
Structural and functional analysis of the PID-3/ERH-2 interaction. (A) GST pulldown assays. GST-tagged PID-3 constructs were co-expressed with Trx-tagged ERH-2 in bacterial cells and then subjected to co-precipitation using glutathione-coupled beads. Input and elution fraction were analyzed on SDS-PAGE gels with Coomassie brilliant blue staining. (B) Top: The crystal structure of the ERH-2ΔC/PID-3^pep^ complex is shown in cartoon representation. The two ERH-2 protomers are shown in different shades of blue, while the two PID-3 peptides are in different shades of pink. Bottom: Zoomed-in view of the interaction between ERH-2 and PID-3^pep^. Interacting residues are shown in stick representation. (C) Binding of PID-3^pep^ to ERH-2 monitored by NMR spectroscopy. Inset: upon incremental addition of PID-3^pep^, the ^1^H-^15^N amide signals of ^15^N-labeled ERH-2 move in slow exchange from their free (shown in red), to their bound state (in maroon) with a 2.3-fold molar excess of PID-3^pep^. Two representative zoomed-in regions of the overlaid spectra are shown (full view in Supplementary Fig. S5B). The arrows indicate the direction of the change in CSP from free to bound states. The CSP values determined for a saturated 1:3 ERH-2:PID-3^pep^ complex are plotted below as a function of residue number. The most highly perturbed amino acid, Thr21, is indicated. The grey dashed line corresponds to the average CSP of all residues. The secondary structure elements as found in the crystal structure are shown above. (D) Mutational analysis of the PID-3/ERH-2 interface. GST-tagged PID-3 wild-type and mutants were co-expressed with wild-type or mutant Trx-tagged wild-type or mutant ERH-2 in bacterial cells as described in (A). Pink circles highlight the bands corresponding to the bait and blue circles the prey in the input. (E) Box-plot showing the percentage of hatched embryos of *pid-3::gfp(wt)* and *pid-3[i182a;v186a]::gfp* and *pid-3[i182e;v186e]::gfp* animals grown at 20 and 25°C. Progeny of 30 different mothers were analyzed for each condition, and development of at least 2600 eggs was scored. A two-sided T-test was used to assess significance. P-values are indicated in the graph. (F-G) Total 21U levels in wildtype and *pid-3[i182a/v186a]*-mutant embryos grown at 20°C (F) and 25°C (G) (n=3). Group means are depicted by red lines and p-values are calculated using two-tailed unpaired t-test. (H-I) Scatter-plots depicting the relative abundance of individual 21U loci in *pid-3[i182a/v186a]*-mutant vs. wildtype embryos grown at 20°C (H) and 25°C (I). RPM, Reads Per Million non-structural sRNA reads.

To obtain quantitative insights into PID-3^pep^/ERH-2 interaction, we used isothermal titration calorimetry (ITC). We determined a dissociation constant (*K*d) of 0.65 µM and a stoichiometry N ∼ 1 (0.97) (Supplemental Fig. S3C, Table 1), consistent with the 2:2 stoichiometry observed in the case of the full-length PID-3/ERH-2 complex by SEC-MALS (Fig. 1C). Finally, we tested if the PID-3^pep^ binding specificity may explain our previous observation that PETISCO specifically incorporates ERH-2, and not its close paralogue ERH-1 (36% sequence identity, 60% similarity; Supplemental Fig. S3D, (Cordeiro Rodrigues et al. 2019)). ERH-1 did not bind PID-3^pep^ in GST pull-down, SEC or, ITC experiments (Supplemental Fig. S3E-G), indicating that PID-3^pep^ can discriminate between the two ERH paralogs. We concluded that the N-terminal extension of the PID-3^RRM^ domain binds to ERH-2, and is likely responsible for the ERH-2 specificity of PETISCO.

**Table 1:** Dissociation constants determined using ITC.

| Protein/Complex | Ligand | $K_d$ (μM) | N | $\Delta H$ (kcal/mol) | -T$\Delta S$ (kcal/mol) |
| --- | --- | --- | --- | --- | --- |
| ERH-2 | PID-3<sup>pep</sup> | $0.65 \pm 0.08$ | 0.97 | $-12.3 \pm 0.1$ | $3.9 \pm 0.2$ |
| ERH-1 | PID-3<sup>pep</sup> | X | X | X | X |
| ERH-2 | TOST-1<sup>pep</sup> | $13 \pm 2^a$ | 1<sup>b</sup> | $-22 \pm 1^a$ | $15 \pm 2^a$ |
| ERH-2 | PID-1<sup>pep</sup> | $24 \pm 3^a$ | 1<sup>b</sup> | $-25 \pm 2^a$ | $19 \pm 2^a$ |
| ERH-2 | TOST-1<sup>FL</sup> | $13 \pm 1^a$ | 1<sup>b</sup> | $-12 \pm 1^a$ | $5.4 \pm 0.5^a$ |
| ERH-2: PID-3<sup>pep</sup> 1:2 | TOST-1<sup>pep</sup> | $4 \pm 2^a$ | 1<sup>b</sup> | $-13 \pm 5^a$ | $6 \pm 6^a$ |
| ERH-2: PID-3<sup>pep</sup> 1:2 | TOST-1<sup>FL</sup> | $15 \pm 2^a$ | 1<sup>b</sup> | $-16 \pm 1^a$ | $9.3 \pm 0.6^a$ |
| ERH-1 | TOST-1<sup>pep</sup> | $10 \pm 2^a$ | 1<sup>b</sup> | $-20 \pm 3^a$ | $14 \pm 4^a$ |
| ERH-1 | PID-1<sup>pep</sup> | $73 \pm 11^a$ | 1<sup>b</sup> | $-32 \pm 7^a$ | $26 \pm 7^a$ |

<sup>a</sup>The values reported correspond to the average value and standard deviation of the mean of two or three measurements at 25 °C. <sup>b</sup>The N value was fixed at 1 based on the known stoichiometries, due to difficulties accurately determining the concentrations of the TOST-1<sup>pep</sup> and PID-1<sup>pep</sup> lacking aromatic amino acids. X denotes no binding. One representative binding isotherm for each is shown in Supplemental Fig. S9.

### Structural insights into the formation of the ERH-2/PID-3 complex

We next determined the crystal structures of a C-terminally truncated ERH-2 (residues 1-99, ERH-2^ΔC^) in the free and PID-3^pep^ bound states, at resolutions of 1.50 Å and 2.17 Å, respectively (Fig. 3B, Supplemental Table S1). The overall structure of ERH-2^ΔC^ is similar to human ERH (Wan et al. 2005) and *S. pombe* Erh (Xie et al. 2019) (Supplemental Fig. S4A). ERH-2^ΔC^ adopts a mixed α-β-fold, with a four-stranded antiparallel β-sheet that packs against three amphipathic α-helices on the backside (Fig. 3B). The front side of the β-sheet mediates dimerization which results in the formation of a pseudo-β-barrel structure. The structure of free ERH-2^ΔC^ is very similar to PID-3^pep^-bound ERH-2^ΔC^, with the exception that the loop connecting helices α1 and α2 is not visible in the electron density, suggesting it becomes ordered upon PID-3^pep^ binding. (Supplemental Fig. S4B).

PID-3^pep^ is unfolded in solution (Supplemental Fig. S4C) but forms an amphipathic α-helix when bound to ERH-2^ΔC^ (Fig. 3B, Supplemental Fig. S4D). PID-3^pep^ shows well-ordered electron density for residues 177-193 (Chain C) and 179-193 (Chain D) and occupies a similar surface on ERH as DGCR8 in the human ERH/DGCR8^peptide^ complex (Kwon et al. 2020), and Mmi1 in the fission yeast Erh1/Mmi1^peptide^ complex (Xie et al. 2019) (Supplemental Fig. S4A). However, while both Mmi1 and DGCR8 bind as extended peptides lacking defined secondary structure, PID-3^pep^ binds as an α-helix (Supplemental Fig. S4A). The hydrophobic interface of the amphipathic PID-3^pep^ helix is formed by Ile182, Val186, Phe187, Val189, and Leu190 and points towards a hydrophobic groove in ERH-2 formed by strands β1, β2, as well as the α1 helix (Fig. 3B and Supplemental Fig. S4E). In addition to the hydrophobic interactions, hydrogen bonds formed between PID-3^pep^ Ser185 and ERH-2 Asp41, as well as between PID-3^pep^ His191 and ERH-2 Thr21 further contribute to the affinity and specificity of the interaction.

In a complementary approach, we employed NMR spectroscopy and assigned the backbone chemical shifts (^1^H-^15^N, C^α^, and C^β^) of free full-length ERH-2. We used these chemical shifts to determine the secondary structural elements of ERH-2, which were fully consistent with the crystal structure, and also showed a disordered C-terminus (residues 104-113, Supplemental Fig. S5A). In addition, the loop region between α1 and α2 encompassing residues 46-55 was found to contain two sets of amide peaks. This indicates the presence of two well-defined alternative conformations that exchange in the millisecond timescale, and explains the lack of order in the crystal structure of free ERH-2^ΔC^ (Supplemental Fig. 4B).

We then monitored the interaction between PID-3^pep^ and ERH-2. Unlabeled PID-3^pep^ was titrated into ^15^N-labeled ERH-2, and the positions of the amide chemical shifts were measured at each point by ^1^H-^15^N HSQC experiments (Fig. 3C and Supplemental Fig. S5B). We observed binding in the slow exchange regime, whereby peaks disappear from one location and reappear at a new position corresponding to the bound complex. This indicates a strong interaction with a dissociation constant in the low micromolar to nanomolar range, in agreement with the ITC data (Supplemental Fig. S3C). We next assigned the chemical shifts of the ERH-2/PID-3^pep^ complex and calculated the chemical shift perturbations (CSPs). The ERH-2 residues most affected by the interaction with PID-3^pep^ were located in β2, with Thr21, Trp22, and Gly23 exhibiting the largest CSPs (Fig. 3C). Mapping of the amide CSPs onto the crystal structure of ERH-2^ΔC^ was consistent with the binding interface seen in the crystal structure (Supplemental Fig. S5C). Taken together, the crystal structure and NMR data provide a clear view for the basis of the ERH-2/PID-3 interaction.

### Mutational analysis of the ERH-2/PID-3 interface

To test the relevance of residues found at the complex interface, we used GST-pulldown experiments with purified proteins. Mutation of PID-3 Ile182 and Ser185 to glutamate residues (I182E; S185E), completely abrogated binding of PID-3^pep^ to ERH-2 (Fig. 3D), and even milder substitutions of PID-3 Ile182 and Val186 to alanine (I182A; V186A) were sufficient to abolish binding (Fig. 3D). Mutation of Thr13 in ERH-2, which lines the hydrophobic pocket (Fig. 4B), to glutamate (T13E), also abrogated binding to PID-3^pep^ (Fig. 4D). *In vivo*, PID-3 (I182E; V186E) worms displayed a full Mel phenotype at 20°C (Fig. 4E), while the milder PID-3 (I182A; V186A) mutations showed a Mel phenotype only at 25°C. Finally, at the subcellular level, loss of ERH-2 binding did not affect the localization of PID-3 (Supplemental Fig. S5D).

**Figure 4.**
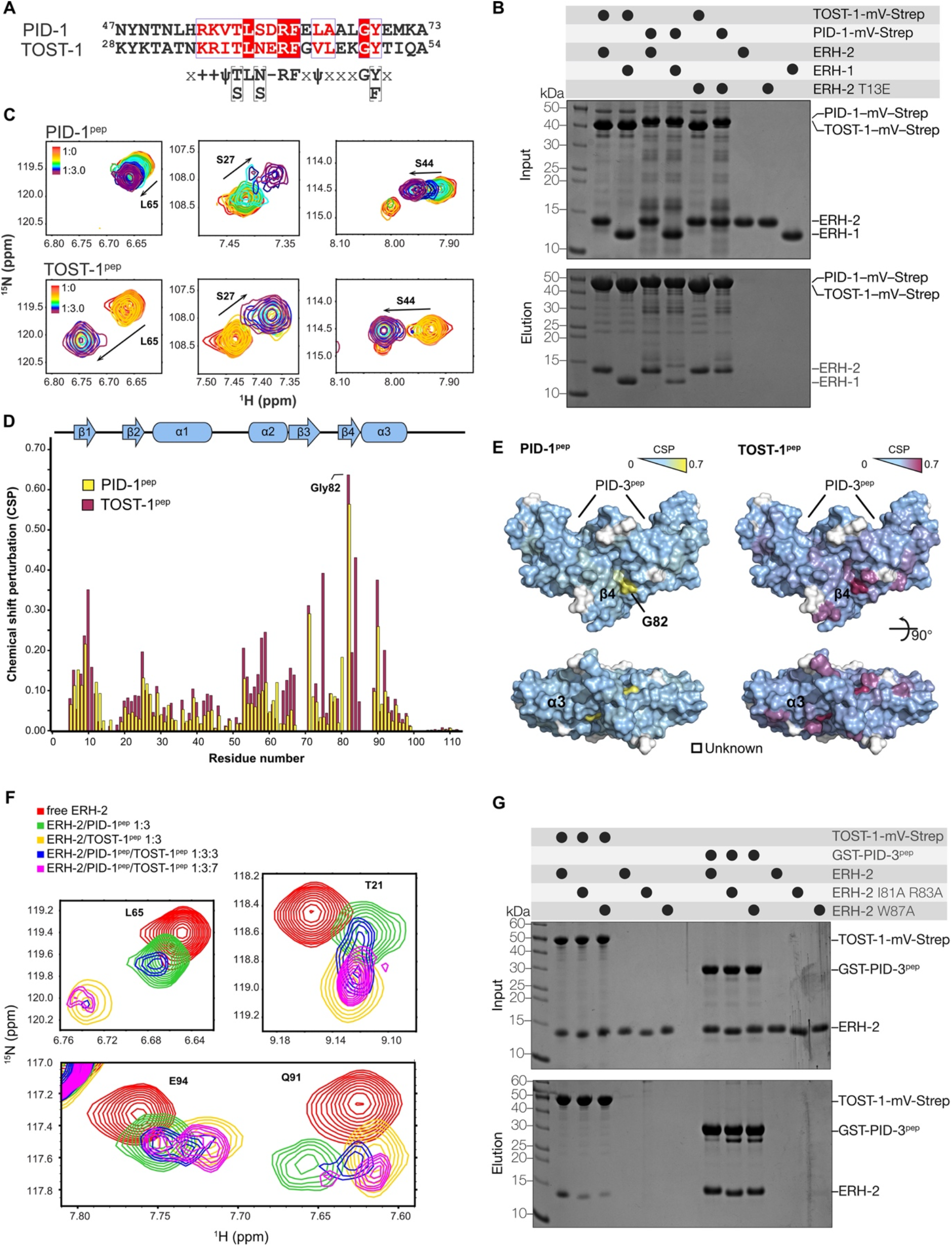
PID-1^pep^ and TOST-1^pep^ bind to ERH-2 through a common interface. (A) Sequence alignment of *C. elegans* PID-1 and TOST-1 corresponding to the conserved motif previously determined (Cordeiro Rodrigues et al. 2019). Analysis of the interaction between TOST-1/PID-1 with ERH-1/ERH-2 by Streptavidin pull-down assays. Purified, recombinant TOST-1 or PID-1 mVenus-Strep fusion proteins were used as baits and ERH-1 and ERH-2 as prey. Input and elution fractions were analyzed by SDS-PAGE followed by Coomassie staining. (C) PID-1^pep^ and TOST-1^pep^ bind to ERH-2 in a similar fashion. NMR-monitored titrations of ^15^N-labeled ERH-2 upon addition of unlabeled TOST-1 and PID-1 peptides, as indicated (full view in Supplemental Fig. S6). (D) The bound ERH-2/peptide complexes (1:3 molar ratios) were assigned and the extent of the CSPs quantified. The peptides produced changes in similar directions (indicated by the arrows), and generally included the same set of perturbed amino acids. (E) Mapping of the CSP values onto the dimeric structure of ERH-2 (PDB: 7O6N) shows that the most affected residues lie in strand β4 and adjacent helix α3, opposite the interface bound by PID-3^pep^. (F) Interplay of TOST-1^pep^ and PID-1^pep^ binding to ERH-2. Comparison of ^15^N-labeled ERH-2 moieties in its free form (red), bound to PID-1^pep^ (green, 1:3 molar ratio), TOST-1^pep^ (yellow, 1:3), in the presence of equimolar amounts of both peptides (blue, 1:3:3), or in the presence of excess TOST-1^pep^ (magenta, 1:3:7). (G) Pull-down assays with purified recombinant wild-type and mutant versions of ERH-2 as prey and TOST-1-mVenus-Strep as bait (streptavidin pull-down) or GST-PID-3^pep^ as bait (GST pull-down). Input and elution fractions were analyzed by SDS-PAGE followed by Coomassie staining.

To analyze the effect of the PID-3 (I182A; V186A) mutation on 21U RNA biogenesis, we sequenced mature piRNAs at permissive (20°C) and restrictive (25°C) temperature in wild-type and mutant animals. This revealed a temperature-sensitive effect on overall 21U RNA levels (Fig. 3I-F). This was found to affect all 21U RNA species (Fig. 3G-H). We note that elevated temperatures also reduced 21U RNA levels in wild-type animals, but to a lesser extent. The reason behind this effect is currently unclear.

These results strongly support the *in vivo* relevance of the structures that we determined, but also suggest that additional interactions between ERH-2 and PID-3 may exist within the full PETISCO complex.

### ERH-2 interacts with the effector proteins TOST-1 and PID-1 through a common interface

PID-1 and TOST-1 share a common sequence motif with which they interact with ERH-2, as previously assessed through Y2H and MS analysis (Fig. 4A) (Cordeiro Rodrigues et al. 2019). To understand the interplay of ERH-2 with these proteins, we purified full-length TOST-1 and PID-1, C-terminally tagged with monomeric Venus StrepII tag (mV-Strep), and probed their binding to ERH-2 and ERH-1. Both PID-1 and TOST-1 bound ERH-2 (Fig. 4B), but unlike PID-3, they also interacted with ERH-1 (Fig. 4B). We noted as well that TOST-1 was a more efficient bait than PID-1 (Fig. 4B), suggesting that it binds more strongly to ERH-2 and ERH-1 than PID-1.

To define the binding interface at the residue level, we analyzed the interaction of PID-1 and TOST-1 with ERH-2 using NMR spectroscopy. We titrated peptides corresponding to the conserved region of TOST-1 (residues 28-53, TOST-1^pep^) and PID-1 (residues 47-74, PID-1^pep^) into ^15^N-labeled full-length ERH-2 to monitor amide CSPs (Fig. 4C and Supplemental Fig, S6A, B). The changes occurred in slow-intermediate exchange for TOST-1^pep^, and in intermediate exchange for PID-1^pep^ (Fig. 4C). This indicates binding constants in the micromolar range for both peptides, with PID-1^pep^ binding being weaker than TOST-1^pep^.

To determine where PID-1^pep^ and TOST-1^pep^ bind ERH-2, we assigned the backbone chemical shifts of the respective complexes and calculated their amide CSPs (Fig. 4D). The changes observed occurred in similar residues and mostly in the same direction, indicating that both peptides bound in analogous fashion and to the same surface of ERH-2 (Fig. 4C). Both peptides caused the largest perturbations at the interface formed by strand β4, including the highly perturbed amino acids Ile81, Gly82, and Arg83, as well as the adjacent *α*3 helix, including Trp87 (Fig. 4D,E). Mapping of the largest CSPs on the surface of ERH-2 showed that the interface bound by TOST-1/PID-1 laid opposite the interface used for PETISCO binding through PID-3^pep^ (Fig. 4E). Of note, we observed only one set of peaks for both the ERH-2 homodimer (save for the flexible loop), as well as the protein-peptide complexes. This suggests a 2:2 stoichiometry for protein to effector peptide, as any other scenario would break the symmetry required to observe only one species in solution.

We then explored the PID-1 and TOST-1 interplay on ERH-2, and compared spectra of free ERH-2 and in the presence of PID-1^pep^, TOST-1^pep^, as well as mixtures of both. When the peptides were present at the same concentrations, we observed chemical shifts at intermediate locations between that of the ERH-2/TOST-1^pep^ and ERH-2/PID-1^pep^ complexes (Fig. 5F).

**Figure 5.**
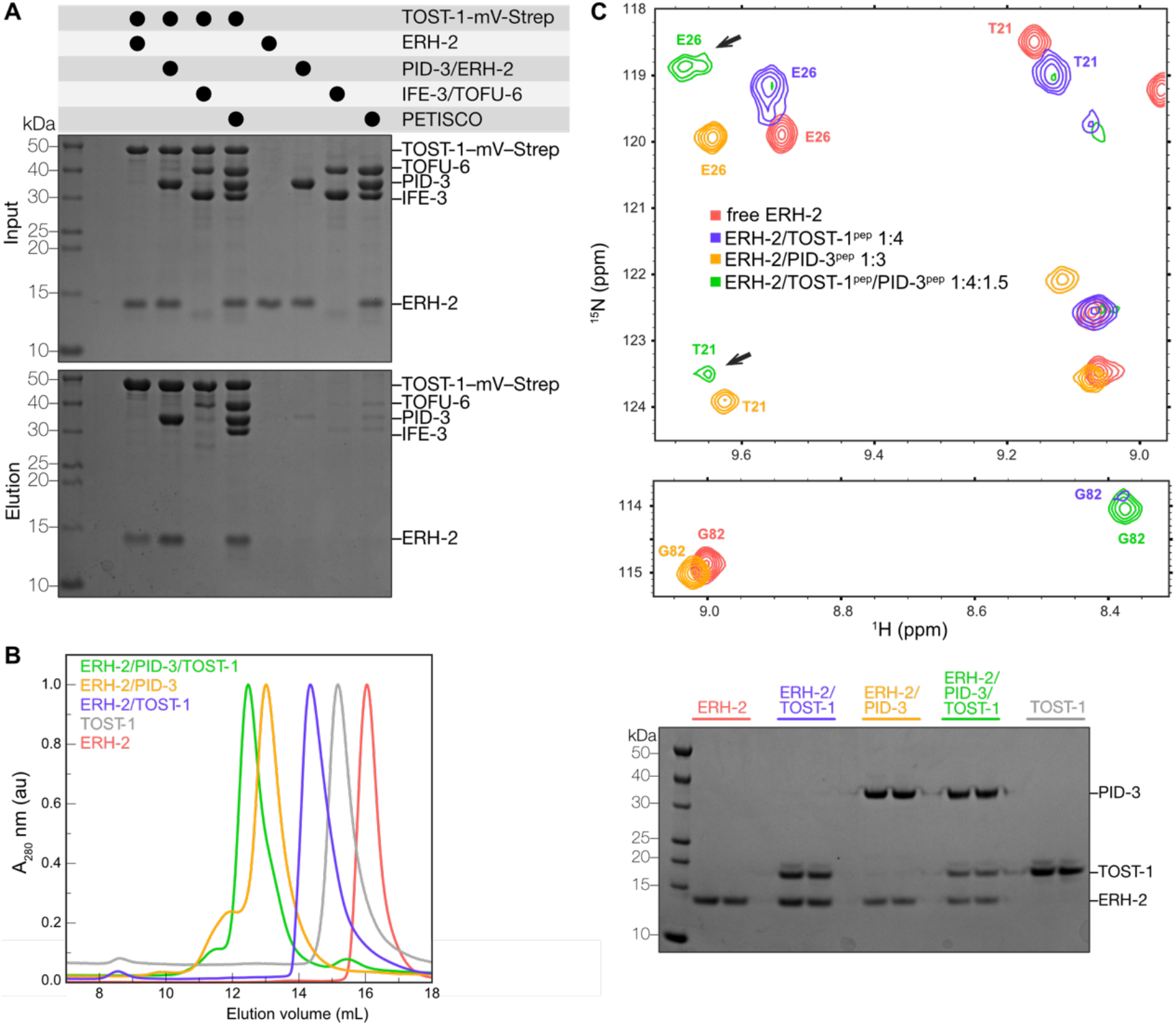
PID-3 and TOST-1 associate with ERH-2 simultaneously. (A) Analysis of the interaction between TOST-1 with PETISCO components. Pull-down experiments with purified, recombinant TOST-1-mVenus-Strep fusion as bait with the indicated preys. Input and elution fractions were analyzed by SDS-PAGE followed by Coomassie staining. (B) Interactions between ERH-2, TOST-1, and PID-3 were assessed by SEC. *Left panel*: Purified proteins were incubated alone or in the indicated mixtures in 1:1 ratio and subjected to SEC*. Right panel*: Analysis of the peak fraction from SEC by SDS-PAGE and Coomassie staining. (C) The formation of a ternary ERH-2/TOST-1^pep^/PID-3^pep^ complex shown by NMR spectroscopy. Zoomed-in views of overlaid ^1^H-^15^N HSQCs of ERH-2 in its free form (red), in complex with TOST-1^pep^ (blue, 1:4 molar ratio), in complex with PID-3^pep^ (orange, 1:3 molar ratio), or in the presence of both (green, 1:4:1.5 ERH-2:TOST-1^pep^:PID-3^pep^). Residues Thr21 (T21), Glu26 (E26), and Gly82 (G82) are indicated, with their positions color coded according to the same scheme. The black arrows highlight amide peaks present in the ternary complex that do not correspond to either the ERH-2/TOST-1^pep^ or ERH-2/PID-3^pep^ subcomplexes.

However, we cannot establish at the microscopic level whether this mixed state arises from rapid exchange between the two complexes in the NMR timescale, or from association of both TOST-1 and PID-1 peptides, one at each available ERH-2 site. Nevertheless, once TOST-1^pep^ was present in excess, we observed chemical shifts consistent with that of an ERH-2/TOST-1^pep^ complex only (Fig. 4F, Supplemental Fig. S7). Since the peptides associate at the same interface and with comparable dissociation constants, we concluded that PID-1 and TOST-1 compete for ERH-2 binding, and their relative concentrations will determine which of the two complexes will be favored. Despite the larger size of the ERH-2/TOST-1^FL^ complex (60 kDa), we detected very similar amide CSPs upon addition of full-length TOST-1 (TOST-1^FL^) to ^15^N-labeled ERH-2, indicating that the shorter peptide recapitulates full-length TOST-1 binding (Supplemental Fig. S8).

To validate the PID-1/TOST-1 binding interface observed by NMR, we designed two ERH-2 mutants (I81A/R83A and W87A), and performed pull-down experiments with purified proteins. Both ERH-2 mutants, in particular W87A, showed a weaker interaction with TOST-(Fig. 4G). These mutations in ERH-2 did not affect PID-3^pep^ binding, and conversely, the mutation in ERH-2 (T13E) that disrupted association with PID-3 did not significantly affect TOST-1 binding (Fig. 4B), consistent with their binding at opposite sides.

### Analysis of the interplay between TOST-1 and PID-3 upon binding to ERH-2

Our analysis suggested that ERH-2 could simultaneously bind TOST-1 and PID-3. To test this, we performed pull-down assays with TOST-1-mVenus-Strep as bait and ERH-2 alone, as part of an ERH-2/PID-3 subcomplex, or with full PETISCO as prey. We found that TOST-1 pulled down ERH-2 in all conditions (Fig. 5A). In addition, we observed that more ERH-2 is pulled down when ERH-2 is in complex with PID-3 or as part of PETISCO, indicating that the presence of PID-3 might facilitate the binding to TOST-1. The formation of a trimeric complex was supported by SEC experiments with purified full-length proteins, as a sample containing ERH-2, TOST-1, and PID-3 elutes at smaller elution volumes than the dimeric subcomplexes (Fig. 5B).

To obtain quantitative insights, we performed ITC measurements. Both TOST-1^FL^ and TOST-1^pep^ bound to ERH-2 with a dissociation constant of 13 µM, and ERH-2 had a higher affinity to TOST-1^pep^ than to PID-1^pep^, both consistent with the NMR experiments (Table 1, Supplemental Fig. S8, Supplemental Fig. S9). The difference in binding affinities for TOST-1^pep^ and PID-1^pep^ was more pronounced for ERH-1, with the dissociation constant differing by more than a factor of 5. However, we note that *in vivo* ERH-1 has not been detected in complex with TOST-1 or PID-1.

Similar binding affinities between ERH-2 and TOST-1^pep^ were obtained when the experiments were performed in the presence of PID-3^pep^ (Table 1 and Supplemental Fig. S9).

We also used NMR spectroscopy to monitor the interplay of TOST-1^pep^ and PID-3^pep^ on ERH- Upon addition of PID-3^pep^ to a saturated ERH-2/TOST-1^pep^ complex, we observed changes in the amide positions of ^15^N-labeled ERH-2 to locations consistent with the formation of an ERH-2/PID-3^pep^ complex. However, signature chemical shifts corresponding to TOST-1-bound ERH-2 remained, suggesting the formation of a ternary complex (Fig. 5C, full view in Supplemental Fig. S10). Consistent with this idea, we observed the appearance of new peaks not present in either the ERH-2/TOST-1^pep^ or ERH-2/PID-3^pep^ subcomplexes (Fig. 5C).

Interestingly, some amide peaks diagnostic of the ERH-2/TOST-1^pep^ interaction became stronger in the presence of PID-3^pep^ (Fig. 5C), suggesting that PID-3^pep^ stabilizes the interaction of TOST-1^pep^ and ERH-2. Given that the two binding sites are connected through a central β-sheet, this synergy in binding could be explained by an allosteric mechanism. Taken together, the pull-down, SEC, ITC, and NMR results clearly show that both PID-3^pep^ and TOST-1^pep^ can simultaneously associate with ERH-2 to form a trimeric complex and that the association of ERH-2 with PID-3^pep^ facilitates its interaction with TOST-1^pep^.

### A composite structural model of the PETISCO core

In the ERH-2^ΔC^/PID-3^pep^ subcomplex PID-3^pep^ contains residues 177-193 and in the PID-3^RRM^/TOFU-6^RRM^ subcomplex the PID-3^RRM^ domain contains residues 196-274. Hence, only two connecting residues between PID-3^pep^ and PID-3^RRM^ are unaccounted for. This allowed us combine both structures into a composite model. Using structure calculation protocols with slow simulated annealing, we generated a structure ensemble of 20 structures. We fixed atoms present in the crystal structures except the linker region connecting PID-3^pep^ and PID-3^RRM^, allowing the two subcomplexes to sample different relative orientations. In the resulting ensemble, ERH-2 is positioned above the PID-3^RRM^ domain through the interaction with PID-3^pep^ (average r.m.s.d. of the Cα atoms of 6 ± 1 Å) (Figure 6A). Because ERH-2 is bound by two PID-3^pep^ moieties, this restricts the flexibility of ERH-2 with respect to the RRM domain core. In addition, we modeled RNA onto the PID-3^RRM^/TOFU-6^RRM^ complex guided by known RRM-RNA structures (Upadhyay and Mackereth 2020; Auweter et al. 2006; Teplova et al. 2016) and we noticed that while the RNA-binding surface of the TOFU-6^RRM^ would be accessible, the RNA-binding surface of the PID-3^RRM^ would be occupied by the ERH-2^ΔC^/PID-3^pep^ module (Figure 6B), suggesting that TOFU-6 is more likely to bind RNA than PID-3.

**Figure 6.**
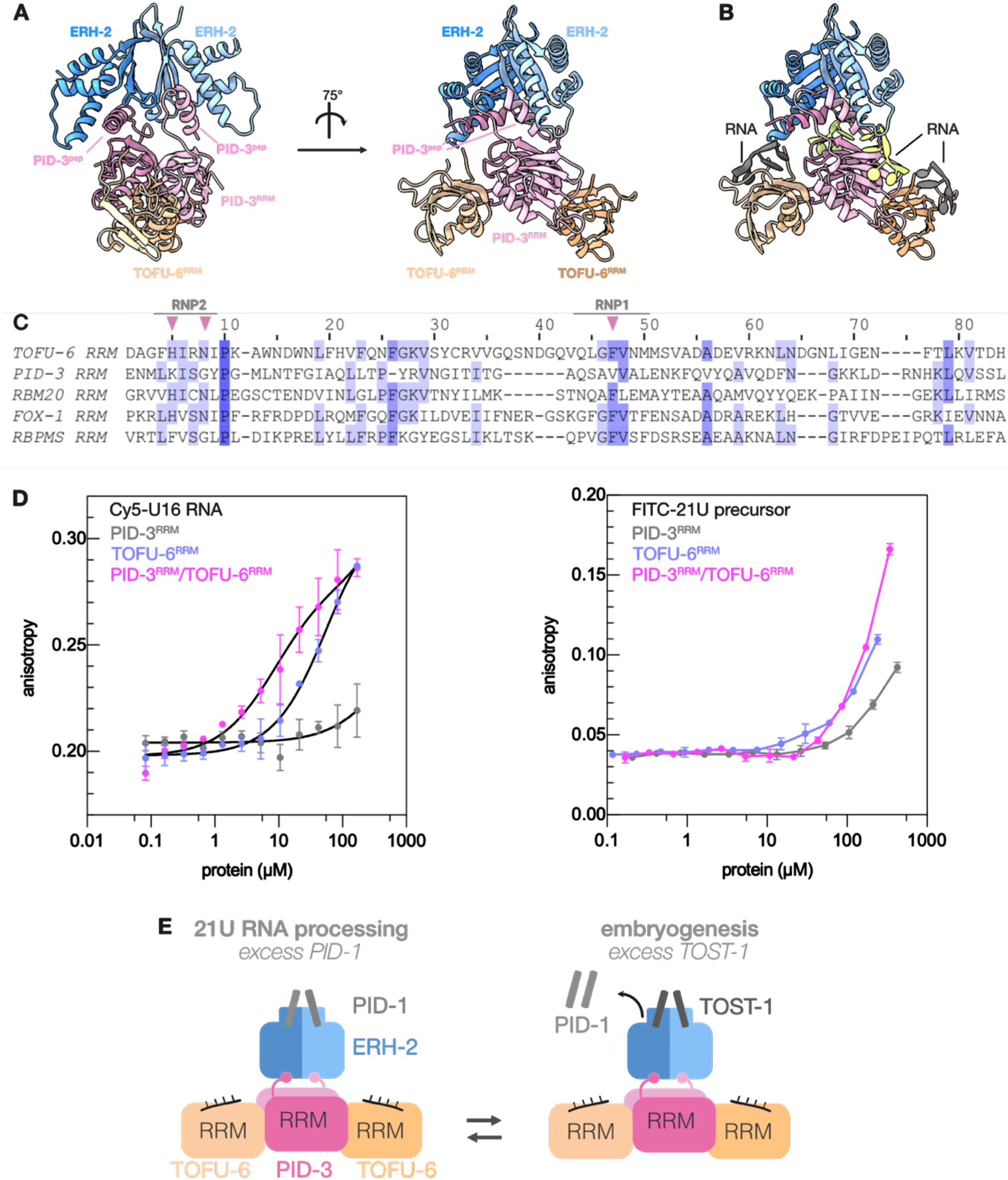
Modelling of the PETISCO core complex. (A) Modelling of the PETISCO core complex by combining the PID-3^RRM^/TOFU-6^RRM^ and ERH-2^ΔC^/PID-3^pep^ subcomplexes. The core complex is shown in two different orientations. The PID-3^RRM^/TOFU-6^RRM^ complex is colored as in Fig. 2, ERH-2^ΔC^/PID-3^pep^ as in Fig. 3 (B) Modelling of RNAs on the four RRMs of the PID-3^RRM^/TOFU-6^RRM^ complex. The position of the four modelled RNA molecules are shown in ribbon presentation. RNAs on the TOFU-6^RRM^ and PID-3^RRM^ are shown in grey and green, respectively. (C) Structure-based multiple sequence alignment of PID-3^RRM^ and TOFU-6^RRM^ in comparison to well-characterized RNA-binding RRMs of RBM20, FOX-1 and RBPMS. The RNP1 and RNP2 motifs contributing to RNA binding are indicated and differences are highlighted with pink triangles. (D) Fluorescence anisotropy binding assay using a Cy5-U16-mer RNA (*left panel*) and a FITC-28-mer RNA (*right panel*). Values are presented as average and error bars correspond to the standard deviation of three (n=3) technical replicates. The solid lines for TOFU-6^RRM^ and PID-3^RRM^/TOFU-6^RRM^ in the left panel represent the fit of a single-binding site model to the data, whereas for PID-3^RRM^ and all three curves in the right panel the solid lines only serve as a guide for the eye. (E) Schematic model of the PETISCO core complex (TOFU-6^RRM^/PID-3^RRM^/ERH-2) and its interaction with the effector proteins PID-1 and TOST-1. TOFU-6^RRM^/PID-3^RRM^ form a tetrameric core and ERH-2 binds to a region upstream of the PID-3^RRM^. TOST-1 and PID-1 bind to a common surface on ERH-2, located opposite to its PID-3 binding site, and thereby specify PETISCO function.

To investigate the potential RNA-binding properties of the TOFU-6^RRM^/PID-3^RRM^ complex in more detail, we compared the RRM domains of both TOFU-6 and PID-3 to well-characterized RRMs using structure-based sequence alignments (Pei et al. 2008). RRM domains bind single-stranded RNA molecules via the outer β-sheet surface with the contribution of two conserved motifs called RNP1 and RNP2, located in the β3 and β1 strands, respectively (Maris et al. 2005). The residues critical for RNA binding, located in RNP1 and RNP2, are retained in TOFU-6^RRM^, whereas the PID-3^RRM^ shows several differences, including the absence of aromatic residues in RNP1 and RNP2 (Fig. 6C).

We next tested RNA binding of PID-3^RRM^, TOFU-6^RRM^, and the PID-3^RRM^/TOFU-6^RRM^ complex experimentally using fluorescence anisotropy assays using two different probes: i) a Cy5-labeled 16-mer oligo(U) RNA and ii) a FITC-labeled 28-mer RNA based on the 21U-3372 precursor (AUUGCAUCUAAAGUUGAUUGAAGAGUUA) containing all four nucleotides and with no predicted secondary structure (Cecere et al. 2012). Consistent with the modelling and sequence analysis, the PID-3^RRM^ showed the lowest binding affinity for both RNAs (Fig. 6D. We were able to fit the data to a single binding site model for the poly(U) probe and derive binding constants for the TOFU-6^RRM^ (*K*d of ∼60 µM) and the tetrameric PID-3^RRM^/TOFU-6^RRM^ complex (*K*d ∼ 10 µM). The higher RNA binding affinity of the PID-3^RRM^/TOFU-6^RRM^ complex could result from additional contribution from the PID-3^RRM^, or from other, currently unknown effects. We also note that the 21U precursor-related probe was bound weaker than the poly(U) RNA. Presently we have no explanation for this observation, but we note that the PETISCO complex also has additional RNA binding domains, for example IFE-3 that can bind 5′-cap structures (Peter et al. 2015), that may well affect overall RNA binding by PETISCO.

## Discussion

### PETISCO - Oligomeric state

We show that PETISCO is an octameric protein complex, consisting of two copies of each IFE-3, TOFU-6, PID-3, and ERH-2 with a mass of ∼236 kDa. We find that the PID-3^RRM^/TOFU-6^RRM^ complex binds a model RNA substrate with higher affinity than the isolated TOFU-6^RRM^, suggesting that oligomerization enhances RNA binding properties, but the functional consequence of PETISCO dimerization for piRNA processing remains to be determined. However, there are interesting parallels with other RNA processing complexes. First, it was recently demonstrated that the yeast and human THO (TREX) complexes involved in the transcription and export of RNA also associate into higher-order oligomers (Schuller et al. 2020; Pühringer et al. 2020). Whereas the yeast THO complex forms a dimer, the human THO complex forms a tetramer (Schuller et al. 2020; Pühringer et al. 2020). Although the functional consequence of oligomerization remains to be shown experimentally, it was hypothesized that dimerization of the yeast THO complex plays a crucial role in preventing R-loop formation during transcription of mRNA by RNA polymerase II (Schuller et al. 2020). In addition, dimerization is required for the function of the *D. melanogaster* SFiNX complex, which facilitates co-transcriptional gene silencing downstream of the piRNA-PIWI complex. Dimerization of SFiNX is promoted through the interaction with the dynein light chain protein ‘Cut up’/LC8, which forms a homodimer and binds a short linear motif present in one of the SFiNX subunits. On the functional level, dimerization of SFiNX is required for the nucleic acid-stimulated formation of biomolecular condensates *in vitro* and heterochromatin formation at piRNA target loci *in vivo* (Schnabl et al. 2021). In the case of SFiNX, dimerization thus plays a direct role in the formation of condensates. Since the proteins that participate in the processing of piRNAs also localize in biomolecular condensates, the P granules, dimerization might also play a role in modulating potential phase separation properties of PETISCO.

Finally, the human ERH homolog has recently been shown to support the microprocessor complex during microRNA (miRNA) processing (Fang and Bartel 2020; Hutter et al. 2020; Kwon et al. 2020). Microprocessor is a trimeric complex consisting of one copy of DROSHA and two copies of DGCR8, and dimerization of DGCR8 is necessary for miRNA processing (Faller et al. 2007). The dimeric ERH protein binds to a short linear motif located in the N-terminal region of DGCR8 and is thought to additionally contribute to DGCR8 dimerization (Kwon et al. 2020). At the functional level, oligomerization facilitates processing of suboptimal miRNA hairpins located in clusters (Fang and Bartel 2020; Hutter et al. 2020). In PETISCO, we find a similar architecture: PID-3 dimerizes through its RRM domain and is supported by the binding of ERH-2 to a motif upstream of the RRM (Figure 6E). Within PETISCO, ERH-2 not only reinforces dimerization, but also binds the effector proteins TOST-1 and PID-1 at a surface opposite of PID-3. Similarly, SAFB has been shown to bind human ERH (Drakouli et al. 2017) and help process suboptimal miRNAs (Hutter et al. 2020; Fang and Bartel 2020). It thus appears that ERH homologs may act as signal integrators to control RNA processing. Because human ERH stimulates processing of suboptimal miRNA precursors, we hypothesize that PETISCO dimerization may likewise stimulate 21U RNA processing. However, dissecting whether these effects stem from dimerization or PETISCO integrity will be difficult, given the inter-dependence of PID-3 dimerization and TOFU-6 binding.

### Subcellular localization of PETISCO

We find that loss of ERH-2 binding by PID-3 does not affect its subcellular distribution over the cytoplasm and P granules. However, we show that dimerization of PID-3 is a prerequisite for TOFU-6 binding and for localization of PID-3 to P granules. Previous work from Zeng et al. (2019) showed that an allele of *tofu-6(ust95)* that results in partial deletion of the eTudor domain specifically affected TOFU-6 P granule localization and 21U RNA production, but not embryonic viability. Together, these results strongly imply that TOFU-6 plays an important role in P granule localization. Interestingly, the depletion of PID-3 by RNAi leads to loss of TOFU-6 from P granules (Zeng et al. 2019), suggesting that PID-3 can somehow enhance TOFU-6’s ability to mediate this function. Whether this relates to the dimerization we describe, or to other effects, cannot currently be resolved.

### Interplay between TOST-1 and PID-1

We find that TOST-1 can outcompete PID-1 for PETISCO binding when present in excess. Is this relevant for PETISCO functionality *in vivo*? A conclusive answer to this question will require an experiment in which the onset of expression of PID-1 and TOST-1 in the various stages of germ cell development can be assessed. Due to the stability of the fluorescent tags, and their large size with respect to the proteins of interest, standard localization studies under steady-state conditions (Zeng et al. 2019) are not suitable to resolve this. However, we do know that PID-1 does not have a role in the embryo, and that 21U RNAs are already expressed at early stages, as judged from PRG-1 expression patterns (Batista et al. 2008; Wang and Reinke 2008). At the same time, germ cells do not show any defects in *tost-1* mutants, while it is present in, and required for the development of embryos (Cordeiro Rodrigues et al. 2019; Zeng et al. 2019). Interestingly, in *tost-1* mutants, some increase in 21U RNA levels have been reported (Cordeiro Rodrigues et al. 2019; Zeng et al. 2019), indicating a competition between the two PETISCO functions. Everything considered, it seems likely that PID-1-bound PETISCO assemblies are present in germ cells when TOST-1 starts to be upregulated, and that the PID-1-TOST-1 exchange that we describe is relevant to establish a good balance between PETISCO functions *in vivo*.

### Evolutionary aspects

Evolutionary analysis of PETISCO components revealed that PID-3, TOFU-6, TOST-1, and PID-1 are restricted to nematodes, while ERH (ERH-2) proteins and eIF4E proteins (IFE-3) are present throughout eukaryotes (Cordeiro Rodrigues et al. 2019). *C. elegans* contains two ERH paralogs, ERH-1 and ERH-2. Strikingly, PID-3 can only interact with ERH-2, suggesting that ERH-2 has adopted a specialized function in PETISCO. Homology modelling of ERH-1 and superposition with our ERH-2 structure revealed that PID-3 binding residues Ile11, Thr13 and Gly20 in ERH-2 are not conserved in ERH-1 (Leu8, Pro10, and Ser20, Supplemental Fig. S3D, S12A). Especially ERH-1 Ser20 sterically interferes with PID-3 binding and we demonstrated that mutation of Thr13 in ERH-2 abrogates PID-3 binding. Interestingly, TOST-1 and PID-1 can, at least *in vitro*, interact with both ERH-1 and ERH-2. Furthermore, TOST-1 is present in nematode species that only contain one ERH paralog (Cordeiro Rodrigues et al. 2019), suggesting that in these species the single ERH protein may execute both PETISCO-related (bound by TOST-1) and PETISCO-unrelated activities.. Interestingly, there are also several nematode species that contain PID-3 but no ERH-2 paralog. Even though in *C. elegans* PID-3 does not bind ERH-1, it would be interesting to see whether PID-3 can interact with ERH-1 in those species, possibly representing an ancestor complex to what we now see in *C. elegans*.

Analysis of sequence conservation revealed that the most conserved feature of the PID-3^RRM^ domain is the surface that mediates homo-dimerization. In contrast, the residues involved in TOFU-6 binding are less well conserved (Supplemental Fig. S12B). On the other hand, most residues of the TOFU-6^RRM^ involved in PID-3^RRM^ binding are well conserved (Supplementary Figure S12C). This is consistent with our previous evolutionary analysis, which revealed that PID-3 is more widespread than TOFU-6 in the nematode phylum (Cordeiro Rodrigues et al. 2019). It might be that the TOFU-6 binding interface’s evolutionary pressure is reduced compared to the PID-3 dimerization interface, or that TOFU-6 is replaced with another protein, possibly also an RRM domain protein, in species that have PID-3 but no TOFU-6.

These aspects clearly show that ERH is a nexus around which many RNA processing reactions concentrate, and which is amenable to significant variation in interacting proteins. The many parallels between ERH function in miRNA processing and PETISCO show that small RNA biogenesis pathways represent one domain of RNA processing that exploits ERH as an interaction platform. However, given the conservation of ERH also in species lacking small RNAs, such as *S. cerevisiae*, and the role of ERH in RNA decay in *S. pombe*, it is clear that ERH proteins fill a niche in RNA processing that is much more general (Weng and Luo 2013).

## Material and Methods

### Protein production

The genes coding for PETISCO subunits (IFE-3, TOFU-6, PID-3, ERH-2), TOST-1, PID-1 and ERH-1 were cloned into modified pET vectors using ligation independent cloning. All proteins were produced as an N-terminal His-Tagged fusion protein with varying fusion partners. IFE-3, ERH-1, ERH-2, PID-1 and TOST-1 contained a His_6_-Trx-3C tag, and TOFU-6 a His_10_-MBP-3C tag and PID-3 a His_6_-GST-3C tag. Addition of 3C protease allowed to cleave this His-fusion protein tag from the protein of interest. Proteins were produced in the *E. coli* BL21(DE3) derivates strain in terrific broth medium with the respective antibiotics. Briefly, cells were grown at 37°C, and when the culture reached an optical density (OD) at 600 nm of 2-3, the temperature was reduced to 18°C. After 2 h at 18°C, 0.2 mM IPTG was added to induce protein production for 12-16 h overnight.

To reconstitute PETISCO, single PETISCO subunits were co-expressed from individual plasmids with different antibiotic resistance markers. IFE-3 (His_6_-Trx-3C tag; ampicillin resistance) was co-expressed with TOFU-6 (His_10_-MBP-3C tag; kanamycin resistance) and PID-3 (His_6_-GST-3C tag; streptomycin resistance) with ERH-2 (His_6_-Trx-3C tag; ampicillin resistance). Cell pellets expressing the IFE-3/TOFU-6 and PID-3/ERH-2 subcomplexes were mixed in a 1:1 ratio and resuspended in lysis buffer (25 mM Tris/HCl, 50 mM NaPO_4_, 250 mM NaCl, 10% (v/v) glycerol, 5 mM 2-mercaptoethanol, pH 7.5) and lysed by sonication. PETISCO was purified by immobilized metal affinity chromatography (IMAC) using Ni^2+-^chelating beads (HisTrap FF; GE Healthcare). The HisTrap FF column was washed with 20 column volumes of lysis buffer and proteins were eluted with 20 mM Tris/HCl pH 7.5, 150 mM NaCl, 500 mM imidazole, 10% (v/v) glycerol, 5 mM 2-mercptoethanol. PETISCO was dialyzed against 20 mM Tris/HCl pH 7.5, 150 mM NaCl, 10% (v/v) glycerol, 5 mM 2-mercaptoethanol and the His-fusion tags were cleaved by the addition of 3C protease during dialysis. The His-fusion tag and His-tagged 3C protease were removed by a second IMAC step. PETISCO was subsequently purified using Heparin affinity chromatography (HiTrap Heparin HP, GE Healthcare)) and size-exclusion chromatography using an S200 increase 10/300 column (GE Healthcare) in a buffer containing 20 mM Tris/HCl pH 7.5, 150 mM NaCl, 10% (v/v) glycerol, 2 mM DTT. All steps were performed on ice or at 4°C.

### Size-exclusion chromatography (SEC) assay

Purified proteins were incubated alone or in different combinations as indicated, in concentrations between 20-40 µM (total volume of 200 µl) in SEC buffer (20 mM Tris/HCl pH 7.5, 150 mM NaCl, 2 mM DTT). Samples were incubated for 1 h on ice to allow complex formation. Complex formation was assayed by comparing the elution volumes in SEC on either a Superdex 200 Increase 10/300 or Superdex 200 Increase 3.2/300 (GE Healthcare) column. The SEC peak fractions were analyzed by SDS–PAGE and visualized by Coomassie brilliant blue staining.

### Size-exclusion chromatography coupled to multi-angle light scattering (SEC-MALS)

The molecular mass and the oligomeric state of PETISCO, its subunits and other proteins in solution were determined by size exclusion chromatography (SEC) coupled to multi-angle light scattering (MALS). Individual proteins or protein complexes were analyzed at concentrations between 2 and 5 mg/mL in a buffer consisting of 20 mM Tris/HCl pH 7.5, 150 mM NaCl, 2 mM DTT. A Superdex 200 Increase 10/300 GL column (GE Healthcare Life Sciences) was connected to a 1260 Infinity HPLC system (Agilent Technologies) coupled to a MiniDawn Treos detector (Wyatt Technologies) with a laser emitting at 690 nm. An RI-101 detector (Shodex) was used for refractive index measurement. Data analysis was performed using Astra 7 software package (Wyatt Technologies).

### ERH-2^ΔC^ and PID-3^pep^ production and purification

ERH-2^ΔC^ containing residues 1-99 of *C. elegans* ERH-2 was cloned into a modified pET-vector with a 6xHis-Thioredoxin (Trx) tag followed by a 3C protease cleavage site. PID-3^pep^ containing residues 171-203 of *C. elegans* PID-3 was cloned into a modified pET-vector with a 6xHis-GST tag followed by a 3C protease cleavage site. Cell growth and protein production was performed as described above. ERH-2^ΔC^ and PID-3^pep^ were purified by immobilized metal affinity chromatography using a Ni^2+^ matrix. The His-Trx and His-GST tags were cleaved with His-tagged 3C protease, and both the tags and 3C protease were removed in a reverse IMAC step. ERH-2^ΔC^ and PID-3pep were present in the flow-through and were concentrated using ultrafiltration and subjected to size-exclusion chromatography on a Superdex 75 (16/600) column equilibrated with 20 mM Tris/HCl pH 7.5, 150 mM NaCl, and 5 mM 2-mercaptoethanol. After size-exclusion chromatography, ERH-2^ΔC^ was concentrated to 15 mg ml^-1^and PID-3^pep^ to 5 mg ml^-1^ and stored at -80°C until further use.

### PID-3^RRM^ and TOFU-6^RRM^ production and purification

TOFU-6^RRM^ (residues 1-99) and PID-3^RRM^ (residues 196-274) were cloned into a modified pET-vector containing His_10_-MBP-3C and His_6_-GST-3C tags, respectively. TOFU-6^RRM^ and PID-3^RRM^ were expressed individually as described for ERH-2^ΔC^ and PID-3^pep^. To purify the PID-3^RRM^/TOFU-6^RRM^ complex, the cultures expressing individual PID-3^RRM^ and TOFU-6^RRM^ proteins were mixed before cell harvest. The PID-3^RRM^/TOFU-6^RRM^ complex was purified by immobilized metal affinity chromatography using a Ni^2+^ matrix, followed by cation-exchange chromatography and size-exclusion chromatography on a Superdex 200 (16/600) column (GE Healthcare) equilibrated in 20 mM Tris/HCl pH 7.5, 150 mM NaCl, and 5 mM 2-mercaptoethanol.

### Crystallization

All crystallization trials were performed using a vapor diffusion set-up by mixing the protein complex and crystallization solution in a 1:1 and 2:1 ratio. For ERH-2^ΔC^, the best diffracting crystals grew in condition A2 from the Morpheus screen (Molecular Dimensions). For crystallization of the ERH-2^ΔC^/PID-3^pep^ complex, these were mixed in a 1:1 ratio. The best diffracting crystals grew in condition A5 from the JCSG+ (Molecular Dimensions). The best crystals of PID-3^RRM^ were obtained in condition C5 from the Morpheus screen (Molecular Dimensions). Crystals were harvested and directly frozen in liquid nitrogen before data collection at 100 K. Crystals from ERH-2^ΔC^, ERH-2^ΔC^/PID-3^pep,^ and the PID-3^RRM^ grew in conditions that did not require further cryoprotection and were therefore directly frozen in liquid nitrogen before data collection at 100 K.

Initial small crystals for the PID-3^RRM^/TOFU-6^RRM^ complex were obtained in several commercial screens in conditions containing sodium acetate buffer at pH 4-5 and various precipitants and additives. Through several rounds of micro-seeding, we obtained larger crystals. The best crystals grew in 0.1 M Sodium acetate pH 5.0, 200 mM NaCl and 17% (v/v) PEG3350. Crystals were soaked with a mother liquor containing 300 mM NaI instead of 200 mM NaCl, and with 25% (v/v) glycerol for cryoprotection, and then frozen in liquid nitrogen.

### Data processing, phase determination, refinement, and modelling

All data were processed with Xia2/Dials (Winter et al. 2013) within CCP4i2 (Potterton et al. 2018). For ERH-2^ΔC^, the phases were determined by molecular replacement using the human ERH structure (PDB 2nml) as model, while in the case of the ERH-2^ΔC^/PID-3^pep^ complex, we used the *C. elegans* ERH-2^ΔC^ structure (PDB 7O6L). The phases of the PID-3^RRM^/TOFU-6^RRM^ complex were solved by single isomorphous replacement with anomalous scattering (SIRAS) phasing from iodide with Autosol from Phenix (Terwilliger et al. 2009). The mean figure of merit over all resolution shells had a value of 0.30 and an estimated map correlation coefficient value of 40 ± 11. The phases for the PID-3^RRM^ were determined by molecular replacement using the *C. elegans* the PID-3^RRM^ domain from the PID-3^RRM^/TOFU-6^RRM^ complex structure (PDB 7OCZ) as model. The models were automatically built using Buccaneer (Cowtan 2006) or Autobuild (Terwilliger et al. 2008), manually completed with COOT (Emsley et al. 2010), and refined with phenix.refine (Afonine et al. 2012) and refmac5 (Kovalevskiy et al. 2018). PDB redo (Joosten et al. 2012) and molprobity (Williams et al. 2018) were used for validation. Data collection, phasing, and refinement statistics are listed in Supplemental Table S1. Molecular graphics of the structures were created using UCSF ChimeraX (Goddard et al. 2018).

### Pull-down assays with purified proteins

For interaction studies with purified proteins, appropriate protein mixtures (bait 10-20 µM, prey in 1.2-fold molar excess) were incubated in 20 mM Tris/HCl (pH 7.5), 150 mM NaCl, 10% (v/v) glycerol, 0.05% (v/v) NP40, 1 mM DTT for 30 min at 4°C. The protein mixtures were then incubated with the indicated beads: Glutathione sepharose beads (Cube Biotech), Amylose sepharose beads (New England Biolabs)), and Strep-Tactin XT beads (IBA) for 2 h. Post incubation, the beads were washed three times with 0.2 mL incubation buffer, and the retained material was eluted with 0.05 mL incubation buffer supplemented with 20 mM of reduced glutathione, 20 mM maltose, or 50 mM biotin. Input material and eluates were analyzed by SDS–PAGE and Coomassie brilliant blue staining.

### Co-expression pull-down assays

For interaction studies by the co-expression co-purification strategy, two plasmids containing the gene of interest and different antibiotic resistance markers were co-transformed into BL21(DE3) derivative strains to allow co-expression. Cells were grown in TB medium at 37°C, and when the culture reached an OD at 600 nm of 2-3, the temperature was reduced to 18°C. After 2 h at 18°C, 0.2 mM IPTG was added to induce protein production for 12-16 h overnight. Cell pellets were resuspended in 2 ml of lysis buffer (50 mM Sodium phosphate, 20 mM Tris/HCl, 250 mM NaCl, 10 mM Imidazole, 10% (v/v) glycerol, 0.05% (v/v) NP-40, 5 mM beta-mercaptoethanol pH 8.0) per gram of wet cell mass. Cells were lysed by ultrasonic disintegration, and insoluble material was removed by centrifugation at 21,000xg for 10 min at 4°C. For GST pull-downs, 500 µL of supernatant was applied to 20 µL glutathione-coupled resin (Cube Biotech); for MBP pull-downs, 500 µL supernatant was applied to 20 µL amylose resin (New England Biolabs) and incubated for two hours at 4°C. Subsequently, the resin was washed three times with 500 µL of lysis buffer. The proteins were eluted in 50 µL of lysis buffer supplemented with 20 mM reduced glutathione or 20 mM maltose in the case of glutathione or amylose beads, respectively. Input material and eluates were analyzed by SDS–PAGE and Coomassie brilliant blue staining.

### Circular dichroism spectroscopy

Far-UV CD spectra were measured using a Chirascan plus spectrometer (Applied Photophysics) and a 0.5-mm path-length cuvette at 22°C. Samples were prepared at 15 µM in 5 mM sodium phosphate, 80 mM NaCl pH 8.0. At least three spectra were collected for each sample at 180-260 nm with 1-nm steps and averaged.

### Multiple sequence alignments

Multiple sequence alignments were generated using MUSCLE (Edgar 2004), structure-based multiple sequence alignments using PROMALS3D (Pei et al. 2008). For the visualization of multiple sequence alignments Jalview 2.0 was used (Waterhouse et al. 2009).

### Isothermal titration calorimetry

All ITC experiments were carried out at 25°C using the PEAQ-ITC Isothermal titration calorimeter (Malvern). The data were processed and curves were fitted using the PEAQ-ITC software. Before the measurements, all samples were dialyzed simultaneously against 1 L of sample buffer (50 mM Tris, 250 mM NaCl, 1 mM TCEP, pH 7.50) to ensure complete buffer matching. The proteins and peptides were concentrated to the values listed below:

| Concentration in the cell ($\mu\text{M}$) | Concentration in the syringe ($\mu\text{M}$) |
| --- | --- |
| 27 $\mu\text{M}$ ERH-1 | 271 $\mu\text{M}$ PID-3<sup>pep</sup> |
| 23 $\mu\text{M}$ ERH-2 | 250 $\mu\text{M}$ PID-3<sup>pep</sup> |
| 20 $\mu\text{M}$ TOST-1<sup>pep</sup> | 600 $\mu\text{M}$ ERH-1 |
| 20 $\mu\text{M}$ TOST-1<sup>pep</sup> | 600 $\mu\text{M}$ ERH-2 |
| 20 $\mu\text{M}$ PID-1<sup>pep</sup> | 600 $\mu\text{M}$ ERH-1 |
| 20 $\mu\text{M}$ PID-1<sup>pep</sup> | 600 $\mu\text{M}$ ERH-2 |
| 20 $\mu\text{M}$ ERH-2 | 431 $\mu\text{M}$ TOST-1<sup>FL</sup> |
| 20 $\mu\text{M}$ ERH-2: 40 $\mu\text{M}$ PID-3<sup>pep</sup> | 431 $\mu\text{M}$ TOST-1<sup>FL</sup> |
| 20 $\mu\text{M}$ ERH-2: 40 $\mu\text{M}$ PID-3<sup>pep</sup> | 285 $\mu\text{M}$ TOST-1<sup>pep</sup> |

In each case, 2 µL of the titrant per injection were added to 200 µL of reactant cell solution. Due to difficulties in accurately measuring the TOST-1^pep^ concentration (low extinction coefficient at 280 nm), this value was floated and the N value fixed at the known stoichiometry of 1. The heats of dilution were obtained by repeating the experiments with sample buffer in the calorimeter cell, while keeping all other experimental parameters identical. These dilution heats were fit to a straight line by linear regression and subtracted from the heats of binding. The baseline-corrected binding isotherms were then fit to a 1:1 model to give the reported thermodynamic parameters. For the purposes of comparing binding of TOST-1 in the presence and absence of PID-3^pep^, the heats of association were assumed to derive solely from TOST-1 binding, as we did not detect binding between TOST-1 and PID-3 peptides (not shown). For multiple measurements, the averages and population standard deviations of the fit results (Kd, ΔH°bind, and -TΔS°bind) are reported.

### Fluorescence anisotropy experiments

The RNAs used for anisotropy experiments were purchased from Ella Biotech (Martinsried, Germany). The pre-piRNA is based on the 21U-3372 precursor sequence.

*5’-Cy5-U16:* Cy5-UUUUUUUUUUUUUUUU

*5’-FITC-pre-piRNA:* FITC-AUUGCAUCUAAAGUUGAUUGAAGAGUUA

The fluorescently labeled RNAs were used at a concentration of 100 nM and incubated with increasing concentrations of protein in a volume of 20 µL in a buffer containing 10 mM Tris/HCl pH 7.5, 50 mM KCl, and 0.5 mM EDTA at 15°C for at least 2 h. The anisotropy was recorded on a FS5 Spectrofluorometer (Edinburgh Instruments) with the following settings: For Cy5: Excitation 649 nm, bandwidth 3 nm; Emission 675 nm, bandwidth 3 nm.

For FITC: Excitation 495 nm, bandwidth 3 nm; Emission 519 nm, bandwidth 3 nm. The anisotropy was measured 5 times in 30 s, and the average anisotropy value was calculated. The data were fitted with a single binding site model using Graphpad Prism.

### Modelling

A homology model of *C. elegans* ERH-1 was generated using the I-TASSER server (Yang et al. 2015) with default settings and the *C. elegans* ERH-2 structure (PDB: 7OCZ) as a template. We generated a model of the PETISCO core complex containing EHR2(1-113) PID3(179-274) TOFU6(RRM) by combining the coordinates of the crystal structures of ERH2(3-99)/PID3(177-193/179-193) and PID3(198-274/196-274)/TOFU6(7-99/7-91). A template structure of the complex was generated in cns-1.2 (Brunger 2007) by adding the missing linker residues (194-197 and 194-195) followed by a short energy minimization. To explore the structural flexibility of this core complex, we generated an ensemble of 20 structures from the energy minimize template by randomly choosing 10 different C_*α*_ pairs of the two subcomplexes and ran a simulated annealing structure calculation protocol (Linge et al. 2003) with one distance restraint for each of the chosen pairs adding or subtracting 7 Å to the distance in the template structure respectively. In the structure calculation, all residues observed in the crystal structures where fixed (Simon et al. 2010) except for eight linker residues (PID3 192-199) connecting the two subcomplexes. The resulting ensemble of structures exhibits an average root mean square deviation of the C_*α*_ atoms of 6 ± 1 Å to the template structure.

### Production of fractionally deuterated ^13^C/^15^N-labeled ERH-2

The gene encoding full-length *C. elegans* ERH-2 (NCBI gene ID: 185323) was cloned into pET28a (Addgene: 69864-3), containing an N-terminal His_6_ tag followed by a TEV protease cleavage site (MGSSHHHHHHSSGENLYFQGHMAS) using NheI and BamHI restrictions sites. Due to the size of the dimeric ERH-2 protein (∼ 27 kDa), fractional deuteration in combination with stable isotope labeling was used for recombinant protein expression in *Escherichia coli* BL21 (λDE3) to allow NMR triple resonance experiments for backbone assignments (^13^C^α^, ^13^C^β^, ^15^NH). A starter 50-mL culture was grown at 37°C for 16 hours with shaking at 200 rpm, using M9 minimal medium (dissolved in H_2_O) supplemented with 2g/L of ^12^C-glucose and 0.5 g/L ^14^NH_4_Cl. The cells were collected by centrifugation and resuspended in 500 mL of M9 minimal medium dissolved in ∼ 98 % D_2_O supplemented with 2g/L ^13^C-glucose and 0.5 g/L^15^NH_4_Cl, followed by incubation for 16 hours at 37°C. The resulting culture was used to inoculate additional 2.5 L of minimal M9/D_2_O/^13^C/^15^N medium, to a starting OD_600_ of ∼ 0.3. After growth at 37°C, protein expression was induced at an OD_600_ of ∼ 1 with 2 mM IPTG, and the culture was allowed to shake for 16 hours at 20 °C. Subsequently, the cells were collected by centrifugation, resuspended in lysis buffer (20 mM Na_2_HPO_4_, 20 mM imidazole, 1 M NaCl, 1 M urea, 10 % v/v glycerol, pH 7.4), and frozen at -20°C until use. The cell pellet was then thawed, and the lysis buffer supplemented with lysozyme (Genaxxon Bioscience) and protease inhibitor cocktail (Roche) according to instructions. After 15-minute incubation on ice, the mixture was homogenized (Microfluidics Corp. Microfludizer M-110L Fluid Processor) by passaging four times. Unless otherwise noted, all purification steps were performed at room temperature. The lysate was cleared by centrifugation, and applied to tandem fast-flow Ni^+2^-NTA HisTrap (GE Healthcare) columns. These were washed with 20 column volumes of lysis buffer. His-tagged ERH-2 was eluted in one step with ∼35 mL of 20 mM Na_2_HPO_4_, 500 mM imidazole, and 0.5 M NaCl, at pH 7.4, and dialyzed against 2 L of TEV cleavage buffer (50 mM Tris, 250 mM NaCl, 1 mM DTT, pH 7.5) for 30 minutes at room temperature. At this point, ∼ 1 mg of TEV protease was added, and ERH-2 was allowed to continue dialyzing in its presence at 4°C for 16 hours. The resulting cleaved product contained five non-native residues at the N-terminus and was further purified using a second HisTrap, followed by size exclusion chromatography column (Superdex 75 16/600, GE Healthcare) equilibrated with NMR buffer (50 mM Tris, 250 mM NaCl, 1 mM TCEP, pH 7.50). The appropriate pure fractions were pooled, and the protein was concentrated using Amicon 3 kDa MWCO centrifugal filters (Merck Millipore). The protein concentration was calculated using the measured absorbance at 280 nm under native conditions and assuming an extinction coefficient of 19940 M^-1^ cm^-1^.

### NMR spectroscopy

NMR experiments were performed at 25°C using a Bruker Avance III 800 MHz spectrometer equipped with a cryoprobe. The ERH-2 sample was concentrated to 200-600 micromolar and contained 0.02% sodium azide, 10 % D_2_O (as a locking agent), and 90 % NMR buffer. For backbone assignments and titrations, 2D and 3D transverse relaxation optimized spectroscopy (TROSY) pulse sequences with ^2^H-decoupling and apodization weighted sampling were utilized (Pervushin et al. 1997; Salzmann et al. 1998; Simon and Köstler 2019). The resulting spectra were processed and analyzed using NMRPipe (Delaglio et al. 1995) and NMR-FAM Sparky (Lee et al. 2015), respectively. 90% of all amides were assigned. The resulting chemical shifts (^1^H^N^, ^15^N, ^13^C^α^, and ^13^C^β^) were deposited under BMRB: 50914 and used as input for secondary structure prediction using the program MICS (Shen and Bax 2012) (Supplementary Figure S5A). To assign the peptide-bound ERH-2 complexes, NMR-monitored titrations were used in combination with TROSY-HNCA experiments to track the new positions of the amide peaks (Salzmann et al. 1998).

### Peptides and NMR-monitored titrations

The lyophilized TOST-1 and PID-1 peptides were purchased from Proteogenix at 95 % purity. Approximately 10 mg of each peptide was resuspended in ∼ 0.8 mL of NMR buffer, and further dialyzed using Pur-A-Lyzer dialysis containers (Sigma) with a MWCO of 1 kDa, against an additional 2L of NMR buffer to ensure complete buffer match. The peptide concentration was typically ∼ 2 mM, calculated using the measured absorbance at 280 nm under native conditions and assuming extinction coefficients of 2980 M^-1^ cm^-1^ for both TOST-1 and PID-1 peptides. The peptides were added in a stepwise manner to labeled ERH-2 at a concentration of ∼ 200 µM, and the changes in chemical shifts were monitored with TROSY-HSQC experiments at each titration point. The extent of amide ^1^H-^15^N chemical shift perturbation (CSP) in free versus saturated peptide-bound ERH-2 were calculated according to Williamson et al. (Williamson 2013) to compensate for frequency bandwidths differences between ^15^N and ^1^H dimensions. The resulting values are plotted as a function of residue number.

### List of peptides used in this study

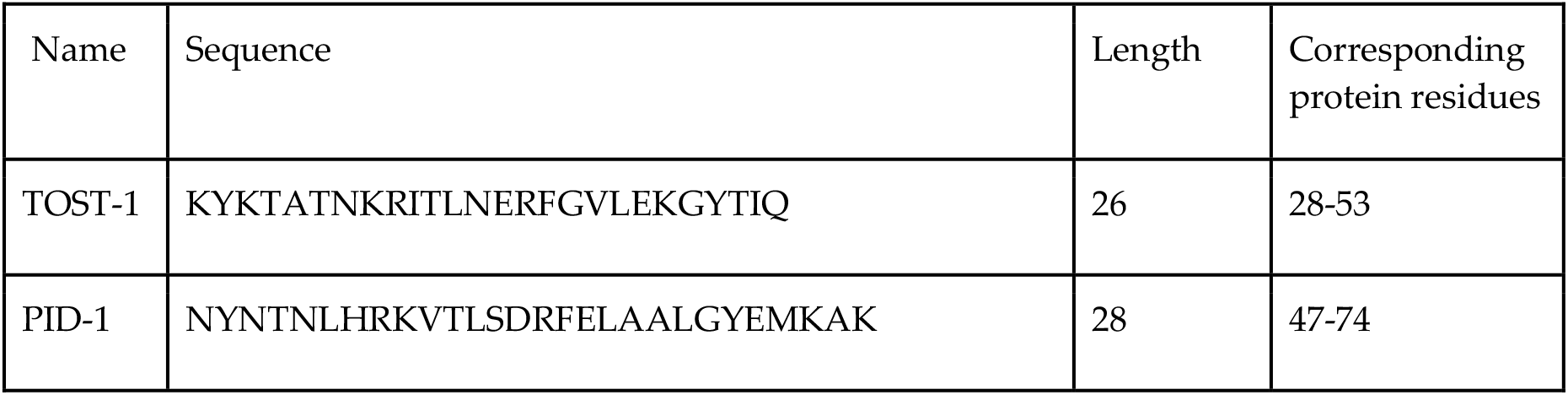
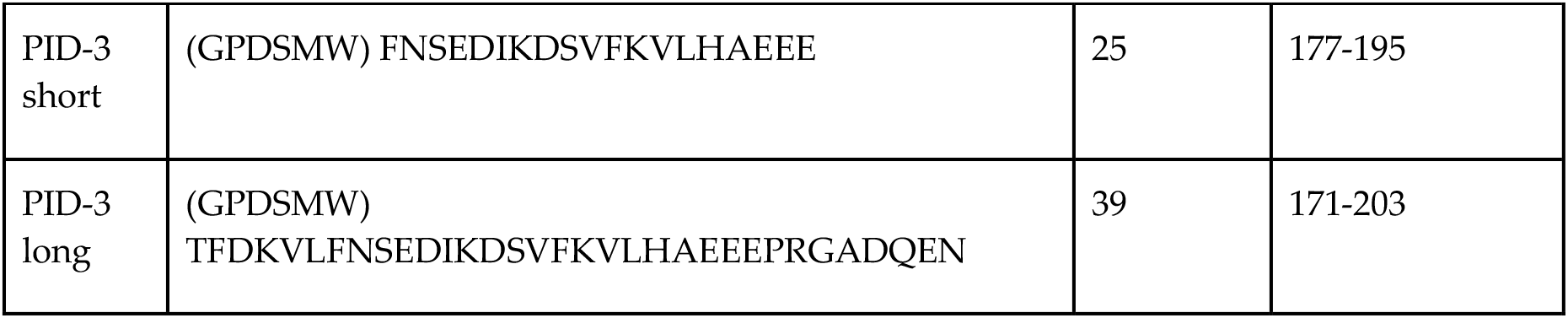

### Worm methods

Worms were cultivated at 20°C on the OP50 plates. For experiments at 25°C, animals were grown at 25°C for 48 hours.

### List of strains

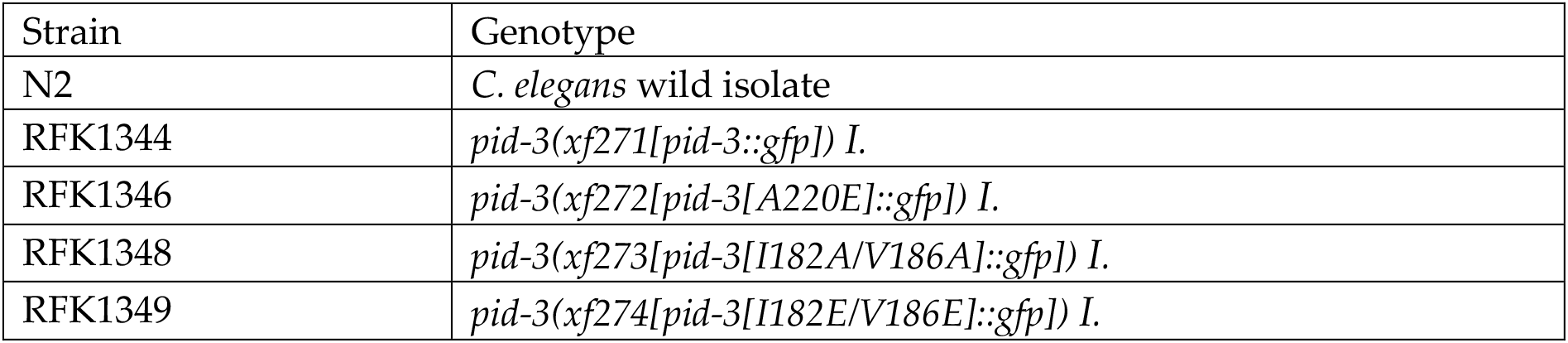

### CRISPR/Cas9 genome editing

Protospacer sequences were chosen using CRISPOR (http://crispor.tefor.net) cloned in pRK2412 by site-directed, ligase-independent mutagenesis (SLIM).

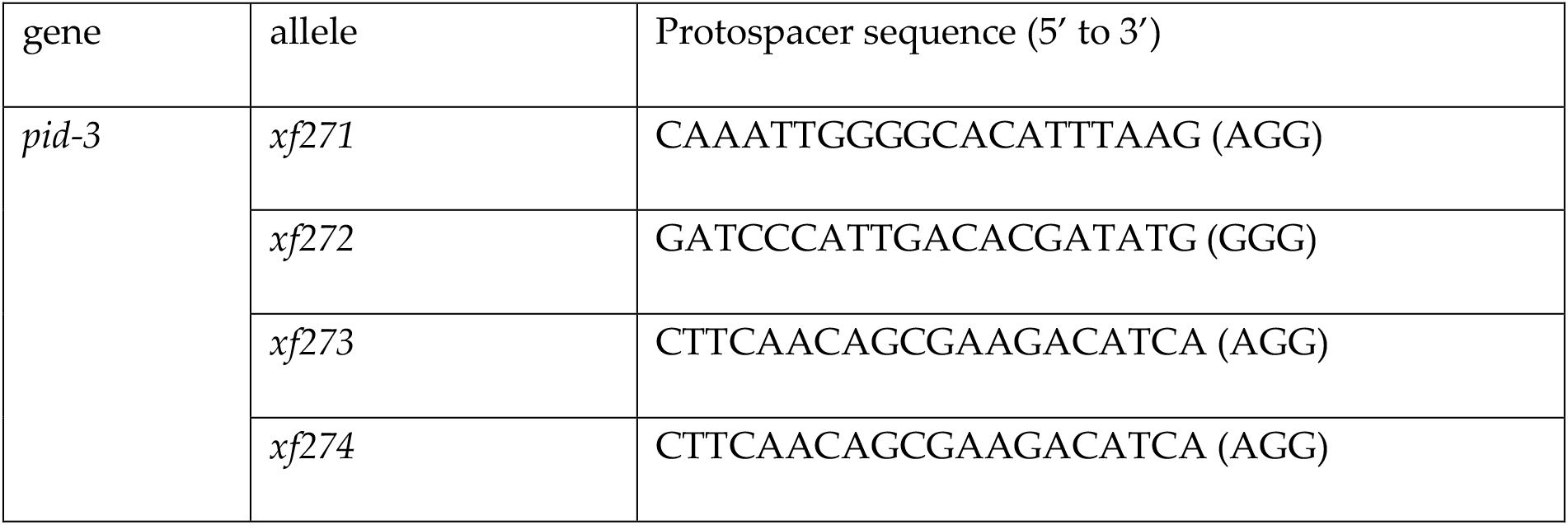

A PCR product from plasmid pDD282 (Addgene plasmid # 66823) was used as a donor template for insertion of *gfp*. For creation of point mutations, the following Ultramer® DNA oligodeoxynucleotides from Integrated DNA Technologies™ were used as a donor template:

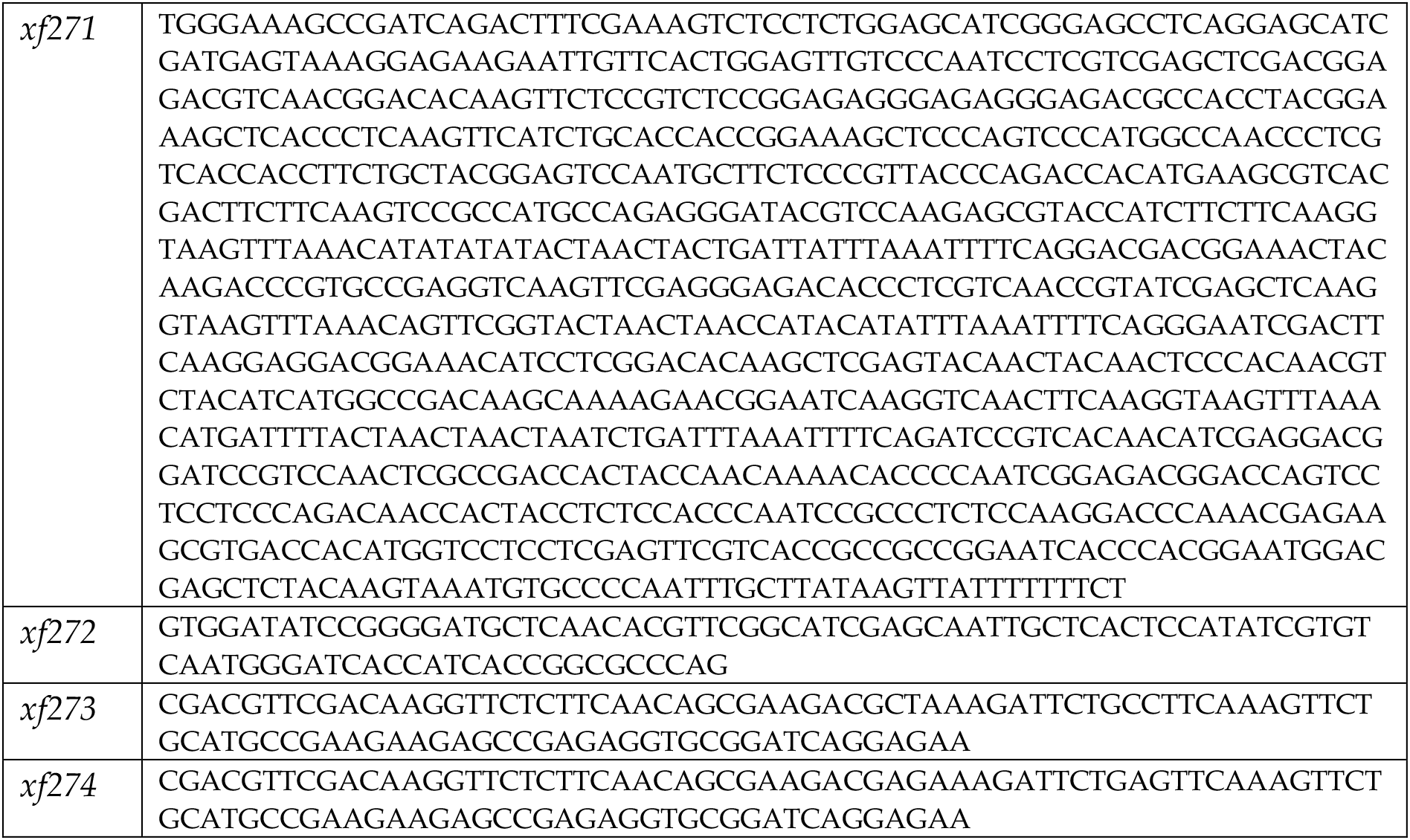

In all cases dpy-10 co-conversion was used. All alleles were outcrossed twice before experiments.

### Mel Phenotype

For RFK1344, 1346 and 1348 30 young adult worms were singled and left for 24 hours (20°C) or overnight (25°C). Afterwards, mothers were removed and the eggs were counted, and after two days, developed animals were scored. RFK1349 is Mel (maternal embryonic lethal) at 20°C, therefore progeny of heterozygous mothers was singled and genotyped. At least 2200 eggs from at least 23 worms were counted for each condition.

### Microscopy

Young adult worms or young gravid adults grown at 25°C were washed in M9 buffer and paralyzed with 60 mM sodium azide in M9. As soon as worms stopped moving, they were imaged on TCS SP5 Leica confocal microscope at 25°C. Images were processed using Fiji and Adobe illustrator.

### 21U RNA Sequencing

NGS library preparation was performed with NEXTflex Small RNA-Seq Kit V3 following Step A to Step G of Bioo Scientific’s standard protocol (V19.01) with a starting amount of 30 ng, amplified in 21 PCR cycles, purified by running on an 8% TBE gel and size-selected in the range of 144-163 nt. The libraries were profiled using a High Sensitivity DNA kit on a 2100 Bioanalyzer (Agilent Technologies) and quantified using the dsDNA HS Assay Kit in a Qubit 2.0 Fluorometer (Life Technologies). All samples were pooled in equimolar ratio and sequenced on a NextSeq 500/550 Flowcell, SR for 75 cycles plus 7 cycles for the index read. The raw sequence reads in FastQ format were cleaned from adapter sequences and size-selected for 18-40 base-long inserts (plus 8 random adapter bases) using cutadapt v.2.4 (http://cutadapt.readthedocs.org) with parameters “-a AGATCGGAAGAGCACACGTCT -m 26 -M 48”. Read alignment to the *C. elegans* genome (Ensembl WBcel235/ce11 assembly) with concomitant trimming of the 8 random bases was performed using Bowtie v.1.2.2 (http://bowtie-bio.sourceforge.net) with parameters “-v 1 -M 1 -y --best --strata --trim5 4 -- trim3 4 -S” and the SAM alignment files were converted into sorted BAM files using Samtools v.1.9 (http://www.htslib.org). *C. elegans* WBcel235/ce11 gene annotation in GTF format was downloaded from Ensembl release 96 (ftp://ftp.ensembl.org/pub/). Aligned reads were assigned to small RNA loci and classes using Samtools, GNU Awk and Subread featureCounts v.1.6.2 (http://bioinf.wehi.edu.au/featureCounts/). Structural reads aligning sense to rRNA, tRNA, snRNA and snoRNA loci were excluded from further analysis. 21U-RNAs were strictly defined as reads 21 base long starting with T and fully overlapping with annotated piRNA (21ur) loci on the same DNA strand. For maximal specificity, excluded from analysis were a small number (3.6%) of ambiguous 21ur loci co-localising on the same strand with miRNAs, snoRNAs or any other RNA exons. The abundance of 21U-RNAs was normalized per million mapped non-structural reads. Sequencing data are available at the NCBI Sequence Read Archive (SRA) under BioProject PRJNA743298 (pre-publication reviewer link: https://dataview.ncbi.nlm.nih.gov/object/PRJNA743298?reviewer=g2qqsvjnla2nn2h0r05bpjf4r3).

### Accession numbers

The coordinates and the structure factors have been deposited in the Protein Data Bank with accession codes PDB ID 7O6L, 7O6N, 7OCX, and 7OCZ. NMR backbone chemical shifts of free *C. elegans* ERH-2 were deposited at the Biological Magnetic Resonance Bank under accession number 50914. Sequencing data are available at the NCBI Sequence Read Archive (SRA) under BioProject PRJNA743298.

## Acknowledgments

We thank all of the members of the Falk, Ketting, and Hennig laboratories for great help and discussion. Svenja Hellmann is thanked for support in genome editing, and the IMB Media Laboratory and the Microscopy Core Facilities for equipment and support. We thank the IMB Genomics Core Facilities for excellent support. We also thank the beamline scientists from the ESRF beamlines ID-23-1 and ID23-2 (Grenoble, France) and Swiss light source PXII (Villigen, Switzerland) for excellent support with data collection, as well as the EMBL Biophysics Facility. The Hennig lab gratefully acknowledges support via an Emmy-Noether Fellowship of the Deutsche Forschungsgemeinschaft (DFG) (HE 7291). This study was supported by funding from the DFG via the priority program SPP1935 to J.H. (EP37/3-1 and EP37/3-2) and R.F.K. (KE1888/7-1) and lab start-up funding from the University of Vienna to S.F. C.P.B. thanks the EMBL Interdisciplinary Postdoc (EI3POD) Program fellowship under Marie Sklodowska-Curie Actions COFUND (grant number 664726) for support.

## Author Contributions

C.P.B., K.H., N.P., R.L., J.B., J.H., and S.F. planned and executed experiments. K.H., R.L. and S.F. purified proteins and performed protein interaction analysis. K.H. and S.F. generated crystals. S.F. and J.B. collected crystallographic data, S.F. solved and modeled all crystal structures. N.P. generated and analyzed *C. elegans* strains and performed protein purifications. C.B.P. and J.H. performed and analyzed ITC and NMR experiments. S.F., R.F.K. and J.H. supervised the projects and provided funds. C.P.B., R.F.K., and S.F. wrote the initial draft article. All authors edited and approved the manuscript.

## References

Afonine PV, Grosse-Kunstleve RW, Echols N, Headd JJ, Moriarty NW, Mustyakimov M, Terwilliger TC, Urzhumtsev A, Zwart PH, Adams PD. 2012. Towards automated crystallographic structure refinement with phenix.refine. Acta Crystallogr D Biol Crystallogr 68: 352–367.

Albuquerque BFM de, de Albuquerque BFM, Luteijn MJ, Cordeiro Rodrigues RJ, van Bergeijk P, Waaijers S, Kaaij LJT, Klein H, Boxem M, Ketting RF. 2014. PID-1 is a novel factor that operates during 21U-RNA biogenesis in Caenorhabditis elegans. Genes & Development 28: 683–688.

Auweter SD, Fasan R, Reymond L, Underwood JG, Black DL, Pitsch S, Allain FH-T. 2006. Molecular basis of RNA recognition by the human alternative splicing factor Fox-1. EMBO J 25: 163–173.

Batista PJ, Ruby JG, Claycomb JM, Chiang R, Fahlgren N, Kasschau KD, Chaves DA, Gu W, Vasale JJ, Duan S, et al. 2008. PRG-1 and 21U-RNAs interact to form the piRNA complex required for fertility in C. elegans. Mol Cell 31: 67–78.

Beltran T, Barroso C, Birkle TY, Stevens L, Schwartz HT, Sternberg PW, Fradin H, Gunsalus K, Piano F, Sharma G, et al. 2019. Comparative Epigenomics Reveals that RNA Polymerase II Pausing and Chromatin Domain Organization Control Nematode piRNA Biogenesis. Dev Cell 48: 793–810.e6.

Beltran T, Pahita E, Ghosh S, Lenhard B, Sarkies P. 2020. Integrator is recruited to promoter-proximally paused RNA Pol II to generate Caenorhabditis elegans piRNA precursors. EMBO J e105564.

Berkyurek AC, Furlan G, Lampersberger L, Beltran T, Weick E-M, Nischwitz E, Cunha Navarro I, Braukmann F, Akay A, Price J, et al. 2021. The RNA polymerase II subunit RPB-9 recruits the integrator complex to terminate Caenorhabditis elegans piRNA transcription. EMBO J 40: e105565.

Brunger AT. 2007. Version 1.2 of the Crystallography and NMR system. Nat Protoc 2: 2728– 2733.

Cecere G, Zheng GXY, Mansisidor AR, Klymko KE, Grishok A. 2012. Promoters recognized by forkhead proteins exist for individual 21U-RNAs. Mol Cell 47: 734–745.

Cordeiro Rodrigues RJ, de Jesus Domingues AM, Hellmann S, Dietz S, de Albuquerque BFM, Renz C, Ulrich HD, Sarkies P, Butter F, Ketting RF. 2019. PETISCO is a novel protein complex required for 21U RNA biogenesis and embryonic viability. Genes Dev 33: 857– 870.

Cowtan K. 2006. The Buccaneer software for automated model building. 1. Tracing protein chains. Acta Crystallogr D Biol Crystallogr 62: 1002–1011.

Czech B, Hannon GJ. 2016. One Loop to Rule Them All: The Ping-Pong Cycle and piRNA-Guided Silencing. Trends Biochem Sci 41: 324–337.

Das PP, Bagijn MP, Goldstein LD, Woolford JR, Lehrbach NJ, Sapetschnig A, Buhecha HR, Gilchrist MJ, Howe KL, Stark R, et al. 2008. Piwi and piRNAs act upstream of an endogenous siRNA pathway to suppress Tc3 transposon mobility in the Caenorhabditis elegans germline. Mol Cell 31: 79–90.

Delaglio F, Grzesiek S, Vuister GW, Zhu G, Pfeifer J, Bax A. 1995. NMRPipe: a multidimensional spectral processing system based on UNIX pipes. J Biomol NMR 6: 277–293.

Drakouli S, Lyberopoulou A, Papathanassiou M, Mylonis I, Georgatsou E. 2017. Enhancer of rudimentary homologue interacts with scaffold attachment factor B at the nuclear matrix to regulate SR protein phosphorylation. FEBS J 284: 2482–2500.

Edgar RC. 2004. MUSCLE: multiple sequence alignment with high accuracy and high throughput. Nucleic Acids Res 32: 1792–1797.

Emsley P, Lohkamp B, Scott WG, Cowtan K. 2010. Features and development of Coot. Acta Crystallogr D Biol Crystallogr 66: 486–501.

Faller M, Matsunaga M, Yin S, Loo JA, Guo F. 2007. Heme is involved in microRNA processing. Nat Struct Mol Biol 14: 23–29.

Fang W, Bartel DP. 2020. MicroRNA Clustering Assists Processing of Suboptimal MicroRNA Hairpins through the Action of the ERH Protein. Mol Cell 78: 289–302.e6.

Goddard TD, Huang CC, Meng EC, Pettersen EF, Couch GS, Morris JH, Ferrin TE. 2018. UCSF ChimeraX: Meeting modern challenges in visualization and analysis. Protein Sci 27: 14– 25.

Goh WSS, Seah JWE, Harrison EJ, Chen C, Hammell CM, Hannon GJ. 2014. A genome-wide RNAi screen identifies factors required for distinct stages of C. elegans piRNA biogenesis. Genes and Development 28: 797–807.

Gu W, Lee H-C, Chaves D, Youngman EM, Pazour GJ, Conte D Jr, Mello CC. 2012. CapSeq and CIP-TAP identify Pol II start sites and reveal capped small RNAs as C. elegans piRNA precursors. Cell 151: 1488–1500.

Hazra D, Andrić V, Palancade B, Rougemaille M, Graille M. 2020. Formation of S. pombe Erh1 homodimer mediates gametogenic gene silencing and meiosis progression. Sci Rep 10: 1034.

Hocine S, Singer RH, Grünwald D. 2010. RNA processing and export. Cold Spring Harb Perspect Biol 2: a000752.

Hutter K, Lohmüller M, Jukic A, Eichin F, Avci S, Labi V, Szabo TG, Hoser SM, Hüttenhofer A, Villunger A, et al. 2020. SAFB2 Enables the Processing of Suboptimal Stem-Loop Structures in Clustered Primary miRNA Transcripts. Mol Cell 78: 876–889.e6.

Joosten RP, Joosten K, Murshudov GN, Perrakis A. 2012. PDB_REDO: constructive validation, more than just looking for errors. Acta Crystallogr D Biol Crystallogr 68: 484–496.

Kasper DM, Wang G, Gardner KE, Johnstone TG, Reinke V. 2014. The C. elegans SNAPc component SNPC-4 coats piRNA domains and is globally required for piRNA abundance. Dev Cell 31: 145–158.

Keiper BD, Lamphear BJ, Deshpande AM, Jankowska-Anyszka M, Aamodt EJ, Blumenthal T, Rhoads RE. 2000. Functional Characterization of Five eIF4E Isoforms inCaenorhabditis elegans *. J Biol Chem 275: 10590–10596.

Kovalevskiy O, Nicholls RA, Long F, Carlon A, Murshudov GN. 2018. Overview of refinement procedures within REFMAC5: utilizing data from different sources. Acta Crystallogr D Struct Biol 74: 215–227.

Kwon SC, Jang H, Shen S, Baek SC, Kim K, Yang J, Kim J, Kim J-S, Wang S, Shi Y, et al. 2020. ERH facilitates microRNA maturation through the interaction with the N-terminus of DGCR8. Nucleic Acids Res 48: 11097–11112.

Lee W, Tonelli M, Markley JL. 2015. NMRFAM-SPARKY: enhanced software for biomolecular NMR spectroscopy. Bioinformatics 31: 1325–1327.

Linge JP, Habeck M, Rieping W, Nilges M. 2003. ARIA: automated NOE assignment and NMR structure calculation. Bioinformatics 19: 315–316.

Luteijn MJ, Ketting RF. 2013. PIWI-interacting RNAs: from generation to transgenerational epigenetics. Nat Rev Genet 14: 523–534.

Maris C, Dominguez C, Allain FH-T. 2005. The RNA recognition motif, a plastic RNA-binding platform to regulate post-transcriptional gene expression. FEBS J 272: 2118–2131.

Ozata DM, Gainetdinov I, Zoch A, O’Carroll D, Zamore PD. 2019. PIWI-interacting RNAs: small RNAs with big functions. Nat Rev Genet 20: 89–108.

Pabis M, Popowicz GM, Stehle R, Fernández-Ramos D, Asami S, Warner L, García-Mauriño SM, Schlundt A, Martínez-Chantar ML, Díaz-Moreno I, et al. 2019. HuR biological function involves RRM3-mediated dimerization and RNA binding by all three RRMs. Nucleic Acids Res 47: 1011–1029.

Pei J, Kim B-H, Grishin NV. 2008. PROMALS3D: a tool for multiple protein sequence and structure alignments. Nucleic Acids Res 36: 2295–2300.

Pervushin K, Riek R, Wider G, Wüthrich K. 1997. Attenuated T2 relaxation by mutual cancellation of dipole-dipole coupling and chemical shift anisotropy indicates an avenue to NMR structures of very large biological macromolecules in solution. Proc Natl Acad Sci U S A 94: 12366–12371.

Peter D, Weber R, Köne C, Chung M-Y, Ebertsch L, Truffault V, Weichenrieder O, Igreja C, Izaurralde E. 2015. Mextli proteins use both canonical bipartite and novel tripartite binding modes to form eIF4E complexes that display differential sensitivity to 4E-BP regulation. Genes Dev 29: 1835–1849.

Potterton L, Agirre J, Ballard C, Cowtan K, Dodson E, Evans PR, Jenkins HT, Keegan R, Krissinel E, Stevenson K, et al. 2018. CCP4i2: the new graphical user interface to theCCP4 program suite. Acta Crystallogr D Struct Biol 74: 68–84.

Pühringer T, Hohmann U, Fin L, Pacheco-Fiallos B, Schellhaas U, Brennecke J, Plaschka C. 2020. Structure of the human core transcription-export complex reveals a hub for multivalent interactions. Elife 9. http://dx.doi.org/10.7554/eLife.61503.

Ripin N, Boudet J, Duszczyk MM, Hinniger A, Faller M, Krepl M, Gadi A, Schneider RJ, Šponer J, Meisner-Kober NC, et al. 2019. Molecular basis for AU-rich element recognition and dimerization by the HuR C-terminal RRM. Proc Natl Acad Sci U S A 116: 2935–2944.

Roundtree IA, Evans ME, Pan T, He C. 2017. Dynamic RNA Modifications in Gene Expression Regulation. Cell 169: 1187–1200.

Ruby JG, Jan C, Player C, Axtell MJ, Lee W, Nusbaum C, Ge H, Bartel DP. 2006. Large-scale sequencing reveals 21U-RNAs and additional microRNAs and endogenous siRNAs in C. elegans. Cell 127: 1193–1207.

Salzmann M, Pervushin K, Wider G, Senn H, Wüthrich K. 1998. TROSY in triple-resonance experiments: new perspectives for sequential NMR assignment of large proteins. Proc Natl Acad Sci U S A 95: 13585–13590.

Schnabl J, Wang J, Hohmann U, Gehre M, Batki J, Andreev VI, Purkhauser K, Fasching N, Duchek P, Novatchkova M, et al. 2021. Molecular principles of Piwi-mediated cotranscriptional silencing through the dimeric SFiNX complex. Genes Dev 35: 392–409.

Schuller SK, Schuller JM, Prabu JR, Baumgärtner M, Bonneau F, Basquin J, Conti E. 2020. Structural insights into the nucleic acid remodeling mechanisms of the yeast THO-Sub2 complex. Elife 9. http://dx.doi.org/10.7554/eLife.61467.

Shen Y, Bax A. 2012. Identification of helix capping and b-turn motifs from NMR chemical shifts. J Biomol NMR 52: 211–232.

Simon B, Köstler H. 2019. Improving the sensitivity of FT-NMR spectroscopy by apodization weighted sampling. J Biomol NMR 73: 155–165.

Simon B, Madl T, Mackereth CD, Nilges M, Sattler M. 2010. An efficient protocol for NMR-spectroscopy-based structure determination of protein complexes in solution. Angew Chem Int Ed Engl 49: 1967–1970.

Sugiyama T, Thillainadesan G, Chalamcharla VR, Meng Z, Balachandran V, Dhakshnamoorthy J, Zhou M, Grewal SIS. 2016. Enhancer of Rudimentary Cooperates with Conserved RNA-Processing Factors to Promote Meiotic mRNA Decay and Facultative Heterochromatin Assembly. Mol Cell 61: 747–759.

Tang W, Tu S, Lee HC, Weng Z, Mello CC. 2016. The RNase PARN-1 Trims piRNA 3′ Ends to Promote Transcriptome Surveillance in C. elegans. Cell 164: 974–984.

Teplova M, Farazi TA, Tuschl T, Patel DJ. 2016. Structural basis underlying CAC RNA recognition by the RRM domain of dimeric RNA-binding protein RBPMS. Q Rev Biophys 49: e1.

Terwilliger TC, Adams PD, Read RJ, McCoy AJ, Moriarty NW, Grosse-Kunstleve RW, Afonine PV, Zwart PH, Hung LW. 2009. Decision-making in structure solution using Bayesian estimates of map quality: the PHENIX AutoSol wizard. Acta Crystallogr D Biol Crystallogr 65: 582–601.

Terwilliger TC, Grosse-Kunstleve RW, Afonine PV, Moriarty NW, Zwart PH, Hung LW, Read RJ, Adams PD. 2008. Iterative model building, structure refinement and density modification with the PHENIX AutoBuild wizard. Acta Crystallogr D Biol Crystallogr 64: 61–69.

Upadhyay SK, Mackereth CD. 2020. Structural basis of UCUU RNA motif recognition by splicing factor RBM20. Nucleic Acids Res 48: 4538–4550.

Wan C, Tempel W, Liu Z-J, Wang B-C, Rose RB. 2005. Structure of the conserved transcriptional repressor enhancer of rudimentary homolog. Biochemistry 44: 5017– 5023.

Wang G, Reinke V. 2008. A C. elegans Piwi, PRG-1, regulates 21U-RNAs during spermatogenesis. Curr Biol 18: 861–867.

Waterhouse AM, Procter JB, Martin DMA, Clamp M, Barton GJ. 2009. Jalview Version 2--a multiple sequence alignment editor and analysis workbench. Bioinformatics 25: 1189– 1191.

Weick E-M, Miska EA. 2014. piRNAs: from biogenesis to function. Development 141: 3458–3471.

Weick E-M, Sarkies P, Silva N, Chen RA, Moss SMM, Cording AC, Ahringer J, Martinez-Perez E, Miska EA. 2014. PRDE-1 is a nuclear factor essential for the biogenesis of Ruby motif-dependent piRNAs in C. elegans. Genes Dev 28: 783–796.

Weng C, Kosalka J, Berkyurek AC, Stempor P, Feng X, Mao H, Zeng C, Li W-J, Yan Y-H, Dong M-Q, et al. 2019. The USTC co-opts an ancient machinery to drive piRNA transcription in C. elegans. Genes Dev 33: 90–102.

Weng MT, Luo J. 2013. The enigmatic ERH protein: Its role in cell cycle, RNA splicing and cancer. Protein and Cell 4: 807–812.

Williams CJ, Headd JJ, Moriarty NW, Prisant MG, Videau LL, Deis LN, Verma V, Keedy DA, Hintze BJ, Chen VB, et al. 2018. MolProbity: More and better reference data for improved all-atom structure validation. Protein Sci 27: 293–315.

Williamson MP. 2013. Using chemical shift perturbation to characterise ligand binding. Prog Nucl Magn Reson Spectrosc 73: 1–16.

Winter G, Lobley CMC, Prince SM. 2013. Decision making in xia2. Acta Crystallogr D Biol Crystallogr 69: 1260–1273.

Xie G, Vo TV, Thillainadesan G, Holla S, Zhang B, Jiang Y, Lv M, Xu Z, Wang C, Balachandran V, et al. 2019. A conserved dimer interface connects ERH and YTH family proteins to promote gene silencing. Nat Commun 10. http://dx.doi.org/10.1038/s41467-018-08273-9.

Yang J, Yan R, Roy A, Xu D, Poisson J, Zhang Y. 2015. The I-TASSER Suite: protein structure and function prediction. Nat Methods 12: 7–8.

Yu S, Kim VN. 2020. A tale of non-canonical tails: gene regulation by post-transcriptional RNA tailing. Nat Rev Mol Cell Biol 21: 542–556.

Zeng C, Weng C, Wang X, Yan YH, Li WJ, Xu D, Hong M, Liao S, Dong MQ, Feng X, et al. 2019. Functional Proteomics Identifies a PICS Complex Required for piRNA Maturation and Chromosome Segregation. Cell Rep 27: 3561–3572.e3.

