## Supplement for "Structural basis of PETISCO complex assembly during piRNA biogenesis in *C. elegans*"

#### Overview of supplemental Figures

**Supplemental Figure S1:** related to Figure 1.

**Supplemental Figure S2**: related to Figure 2.

**Supplemental Figure S3**: related to Figure 3.

**Supplemental Figure S4**: related to Figure 3.

**Supplemental Figure S5**: related to Figure 3.

**Supplemental Figure S6:** related to Figure 4.

**Supplemental Figure S7:** related to Figure 4.

**Supplemental Figure S8:** related to Figure 4.

**Supplemental Figure S9:** related to Figure 5.

**Supplemental Figure S10:** related to Figure 5.

**Supplemental Figure S11:** related to Figure 6.

**Supplemental Figure S12:** related to the discussion

### Supplemental Figure S1

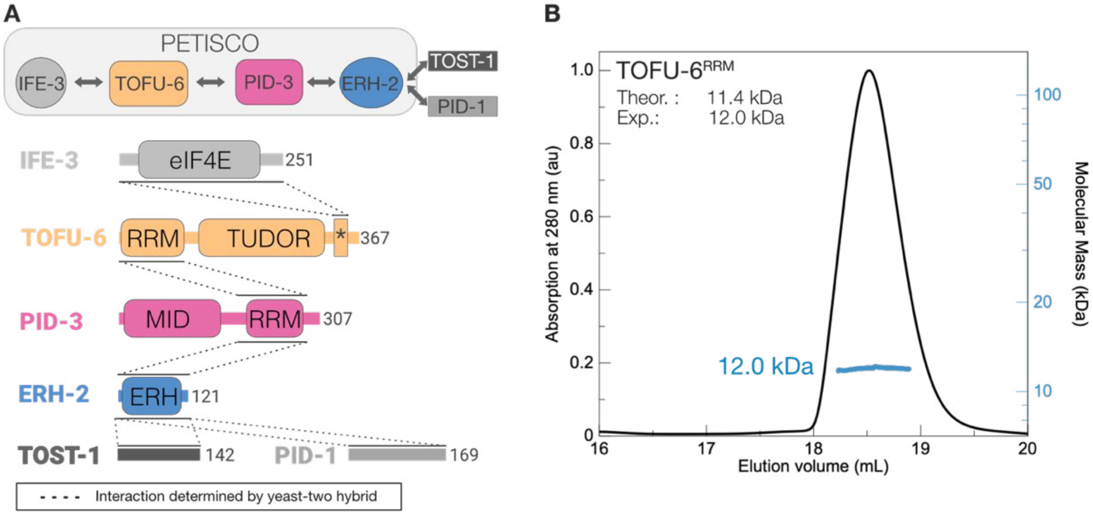


C

| Sample | Polydispersity Mw/Mn | Mw $_{theor}$(kDa) | Mw $_{exp}$(kDa) | oligomeric state |
|---|---|---|---|---|
| PETISCO | 1.000 | 118.5 | 231.1 | heterooctamer (2:2:2:2) |
| IFE-3/TOFU-6 | 1.001 | 70.0 | 67.5 | heterodimer (1:1) |
| PID-3/ERH-2 | 1.000 | 48.5 | 91.6 | heterotetramer (2:2) |
| ERH-2 | 1.000 | 13.3 | 26.7 | homodimer (1:1) |
| PID-3$^{MID}$ | 1.000 | 17.7 | 17.9 | monomer |
| PID-3$^{RRM\text{-}NC}$ | 1.001 | 14.6 | 28.1 | homodimer (1:1) |
| IFE-3 | 1.001 | 28.2 | 28.3 | monomer |
| TOFU-6$^{RRM}$ | 1.000 | 11.4 | 12.0 | monomer |
| PID-3$^{RRM}$/TOFU-6$^{RRM}$ | 1.000 | 20.5 | 39.6 | heterotetramer (2:2) |
| PID-3$^{RRM\text{-}N}$ | 1.002 | 12.1 | 23.8 | homodimer (1:1) |
| PID-3$^{RRM\text{-}N}$ A220E | 1.001 | 12.1 | 12.7 | monomer |
| PID-3$^{RRM\text{-}N}$ F247E Q251R | 1.000 | 12.1 | 21.6 | homodimer (1:1) |
| TOST-1 | 1.000 | 16.6 | 15.5 | monomer |

**Supplemental Figure S1.** PETISCO has a linear topology. (A) Interaction network between individual PETISCO proteins (IFE-3, TOFU-6, PID-3 and ERH-2) and the effector proteins TOST-1 and PID-1. Dashed lines indicate the interactions previously determined by yeast two-hybrid experiments. (B) SEC-MALS profile of TOFU-6$^{RRM}$ reveals that it is a monomer. (C) Extended summary SEC-MALS data with molecular masses, polydispersity (Mn/Mw) and stoichiometries of PETISCO and its subunits.

#### Supplemental Figure S2

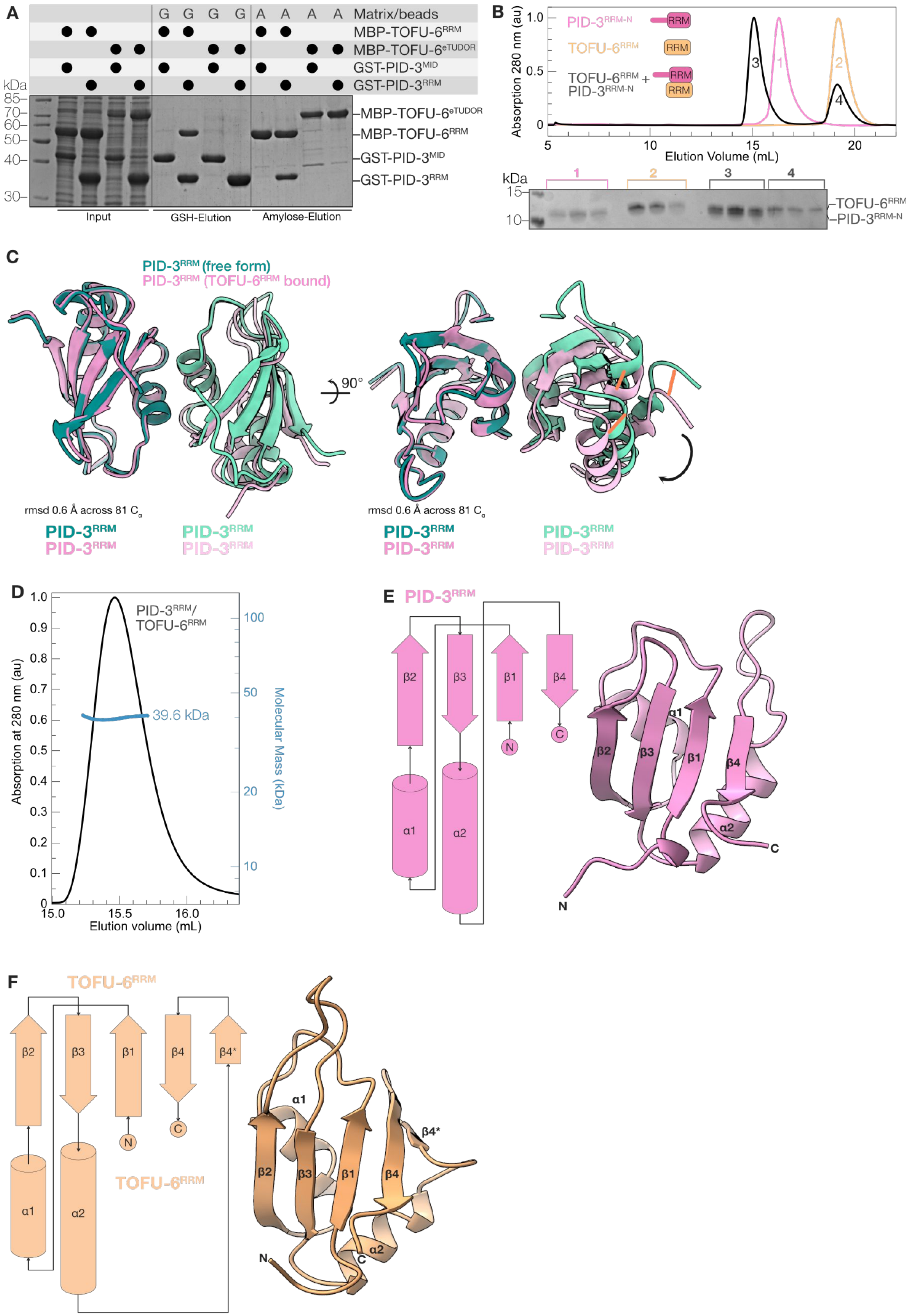

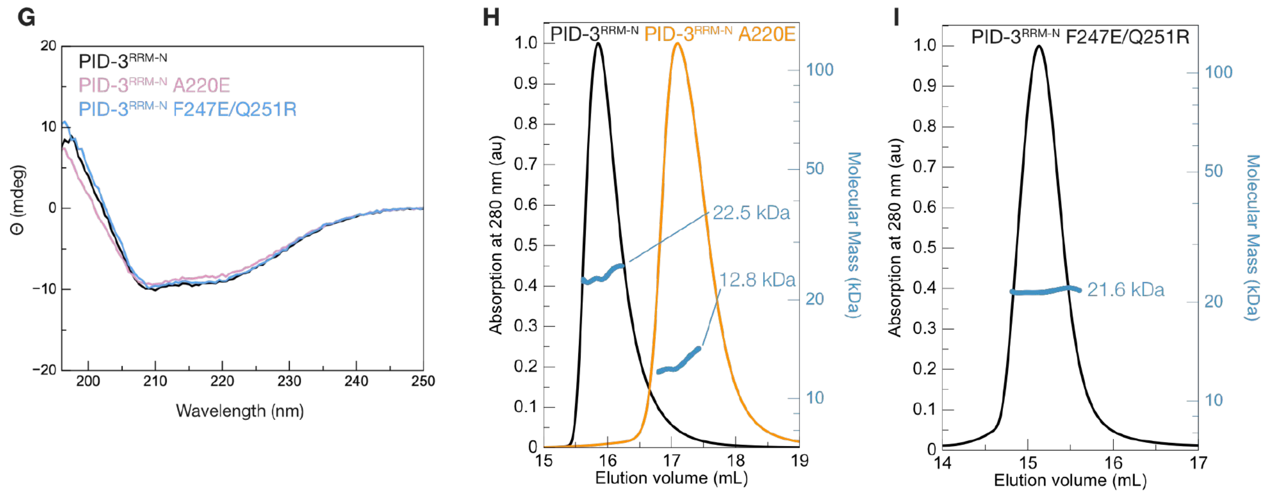


**Supplemental Figure S2.** Biochemical and structural analysis of the PID-3 and TOFU-6 RRM domains. (A) Protein pulldown assays. MBP-tagged TOFU-6 constructs (MBP-TOFU-6RRM and MBP-TOFU-6eTUDOR) were co-expressed with GST-tagged PID-3 constructs (PID-3$^{MID}$ or PID-3$^{RRM}$) in bacterial cells and then subjected to co-precipitation using glutathione and amylose-coupled matrix. The matrix used is indicated above the lane: MBP-binding amylose resin [**A**] and GST-binding GSH resin [**G**]. Input and elution fraction were analyzed on SDS-PAGE gels with Coomassie brilliant blue staining. (B) SEC assays to assess the interaction between PID-3$^{RRM}$ and TOFU-6$^{RRM}$. Purified samples were incubated and injected alone or co-injected (in a 1:1.5 molar ratio) on a size-exclusion column (Superdex 200 Increase 10/300, GE Healthcare). Top: Overlay of the chromatograms; the absorption at 280 nm was normalized to the highest peak. Bottom: Coomassie-stained SDS-PAGE gel showing the corresponding peak fractions. (C) Comparison of the PID-3$^{RRM}$ structures in the free (green colors) and the in the TOFU-6$^{RRM}$ bound form (pink colors). Both structures were superposed on one protomer and reveal a relative movement of the second protomer. (D) SEC-MALS analysis of the PID-3$^{RRM}$/TOFU-6$^{RRM}$ complex. The chromatograms show the UV absorption at 280 nm and the calculated molecular mass in kDa. The UV absorption signal was normalized to the highest peak. The analysis was performed using a Superdex 200 10/300 column (GE Healthcare). (E) Left: Topology diagram of the PID-3$^{RRM}$ domain. The secondary structure elements are numbered, β-strands are shown as arrows and α-helices as cylinders. Right: structure of an isolated PID-3$^{RRM}$ protomer. (F) Left: Topology diagram of the TOFU-6$^{RRM}$ domain. The secondary structure elements are numbered, β-strands are shown as arrows and α-helices as cylinders. Right: structure of an isolated TOFU-6$^{RRM}$ domain. (G) Far-UV

CD spectra of PID-3$^{RRM-N}$ wild type and the A220E and F247E/Q251R mutants. The spectra show two minima around 208 and 220 nm, indicative for the presence of $\alpha$-helices and a maximum around 195 nm, indicating that the proteins are folded. (H-I) The A220E mutation renders the PID-3$^{RRM}$ monomeric, while the F247E/Q251R mutant is homodimeric. SEC-MALS analysis of wild-type (black) and the A220E mutant (orange) PID-3$^{RRM}$ domain (H) and the F247E/Q251R mutant (black) PID-3$^{RRM}$ domain (I). Note: The SEC MALS runs in (H) and (I) were performed using different instruments, therefore the absolute elution volumes cannot be compared. The chromatograms show the UV absorption at 280 nm and the calculated molecular mass in kDa. The UV absorption signal was normalized to the highest peak. The analysis was performed using a Superdex 200 10/300 column (GE Healthcare).

#### Supplemental Figure S3

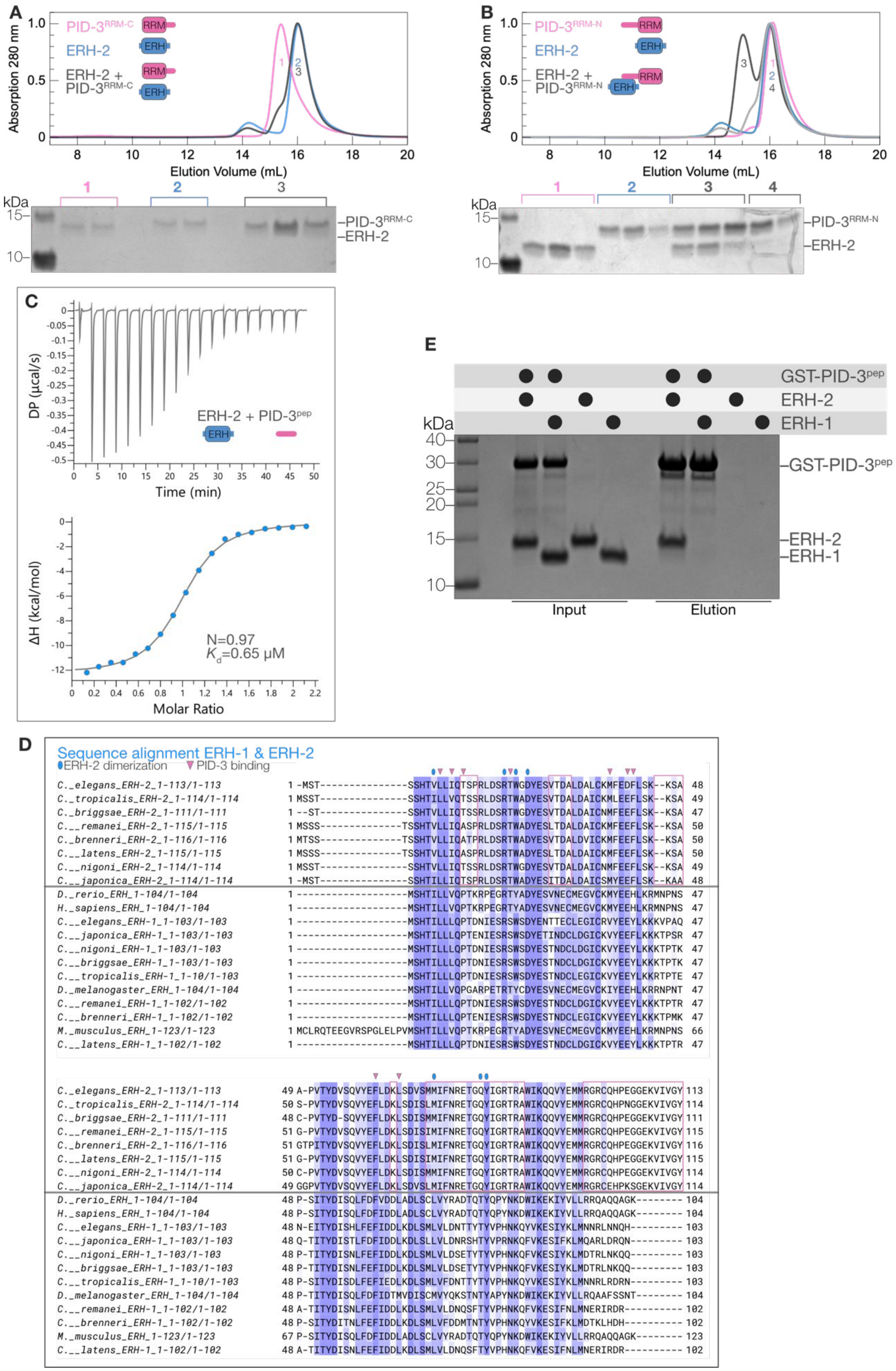

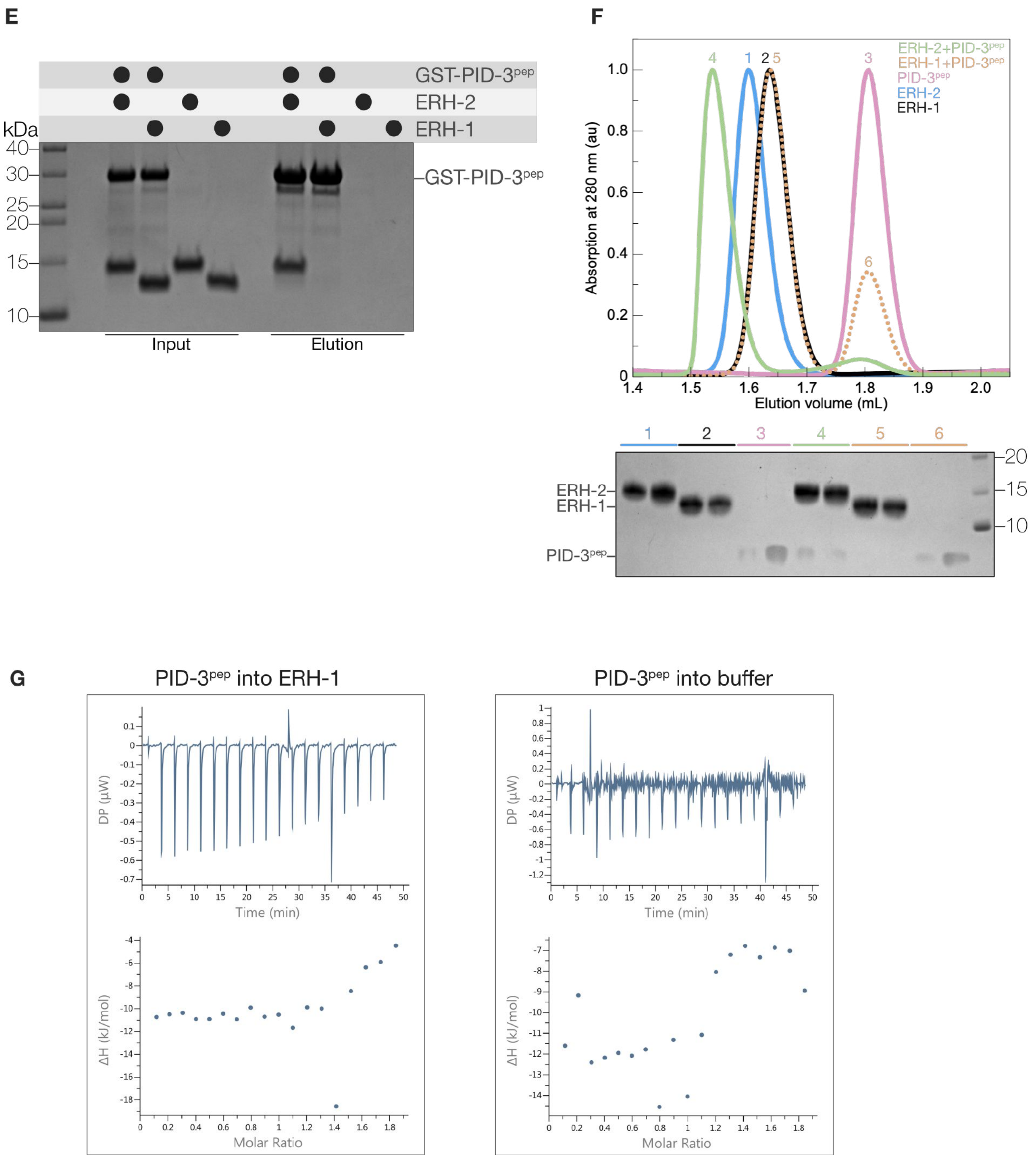


**Supplemental Figure S3.** ERH-2 binds a region upstream of the PID-3$^{RRM}$. (A, B) SEC assays to assess the interaction between different PID-3$^{RRM}$ constructs and ERH-2. Purified PID-3$^{RRM}$ domains carrying either a C-terminal (PID-3$^{RRM-C}$, (A) or an N-terminal (PID-3$^{RRM-N}$, (B)) extension were incubated alone or together with ERH-2 while keeping PID-3$^{RRM}$ in excess, and injected on a size-exclusion column (Superdex 200 increase 10/300, GE Healthcare). Top: Overlay of the chromatograms; the absorption at 280 nm was normalized to the highest peak. Bottom:

Coomassie-stained SDS-PAGE gel showing the corresponding peak fractions. (C) Isothermal titration calorimetry (ITC) experiments analyzing the interaction between ERH-2 and PID-$3^{pep}$. The calculated dissociation constant $K_d$ and the number of binding sites (N) are shown. (D) Multiple sequence alignment of ERH proteins from various species. Most species contain only one ERH gene/protein, whereas some nematodes contain two ERH paralogs, ERH-1 and ERH-2. ERH-1 and ERH-2 protein sequences cluster in distinct groups. *C. elegans* ERH-2 residues involved in homodimerization are highlighted with a blue oval, while residues that mediate binding to PID-$3^{pep}$ are highlighted with pink triangles above the sequence. The regions in which ERH-1 and ERH-2 differ most significantly are highlighted with pink boxes. (E) PID-$3^{pep}$ binds to ERH-2 but not to ERH-1. GST-tagged PID-$3^{pep}$ was incubated with purified ERH-1 and ERH-2 proteins and subjected to a pull-down with glutathione beads. Input and elution fraction were analyzed on SDS-PAGE gels with Coomassie brilliant blue staining. (F) SEC assays to assess binding of PID-$3^{pep}$ to ERH-1 and ERH-2. Purified PID-$3^{pep}$, ERH-1 and ERH-2 were incubated alone or with PID-$3^{pep}$ in 1.2 fold molar in excess, and injected on a size-exclusion column (Superdex 200 increase 3.2/300, GE Healthcare). Top: Overlay of the chromatograms with the absorption at 280 nm normalized to the highest peak. Bottom: Coomassie-stained SDS-PAGE gel showing the corresponding peak fractions. (G) Isothermal titration calorimetry (ITC) experiments analyzing the interaction between ERH-1 and PID-$3^{pep}$. The injection of PID-$3^{pep}$ into the ITC buffer in the cell generated a similar signal as PID-$3^{pep}$ injection into the cell containing ERH-1. Therefore, we concluded that ERH-1 and PID-$3^{pep}$ do not interact.

#### Supplemental Figure S4

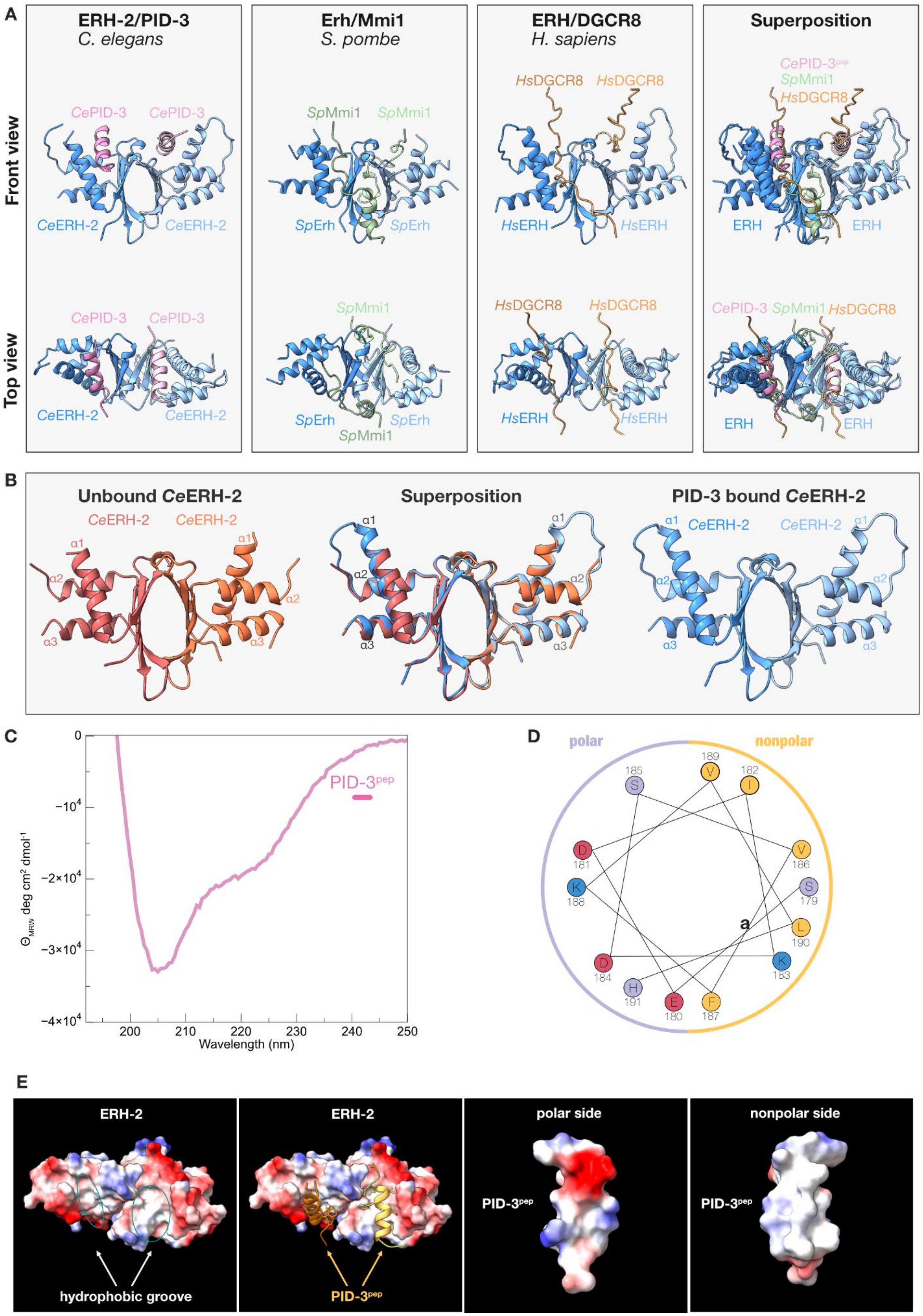

**Supplemental Figure S4.** (A) Structural comparison of ERH-containing protein complexes. The *C. elegans* ERH-2/PID-3, the fission yeast (*S. pombe*) Erh/Mmi1 and human (*H. sapiens*) ERH/DGCR8 complexes are shown in two different orientations. The two ERH protomers constituting the homodimer are shown in different shades of blue, CePID-3 in pink, SpMmi1 in green, and HsDGCR8 in orange. CePID-3, SpMmi1, and HsDGCR8 all bind to a similar ERH surface. On the right hand-side, a superposition of all three complexes is shown. (B) Comparison of the ERH-2 structures in the unbound (red colors) and PID-3$^{pep}$ bound state (pink colors). (C) PID-3$^{pep}$ is unfolded in solution. The far-UV CD spectrum of PID-3$^{pep}$ shows a minimum close to 200 nm, indicative of the absence of secondary structure. (D) PID-3$^{pep}$ forms an amphipathic helix. Helical wheel representation of PID-3$^{pep}$. Nonpolar residues are shown in yellow, positively charged residues in blue, negatively charged residues in red and polar residues in purple. The polar and nonpolar side of the PID-3$^{pep}$ $\alpha$-helix is indicated. (E) PID-3$^{pep}$ binds into a hydrophobic cleft in ERH-2. Surface presentation of ERH-2 colored according to the electrostatic surface potential, where blue and red highlight positively and negatively charged regions, respectively and white indicates uncharged regions. The PID-3$^{pep}$ is shown in two orientations, illustrating the amphipathic character of the helix.

Supplemental Figure S5

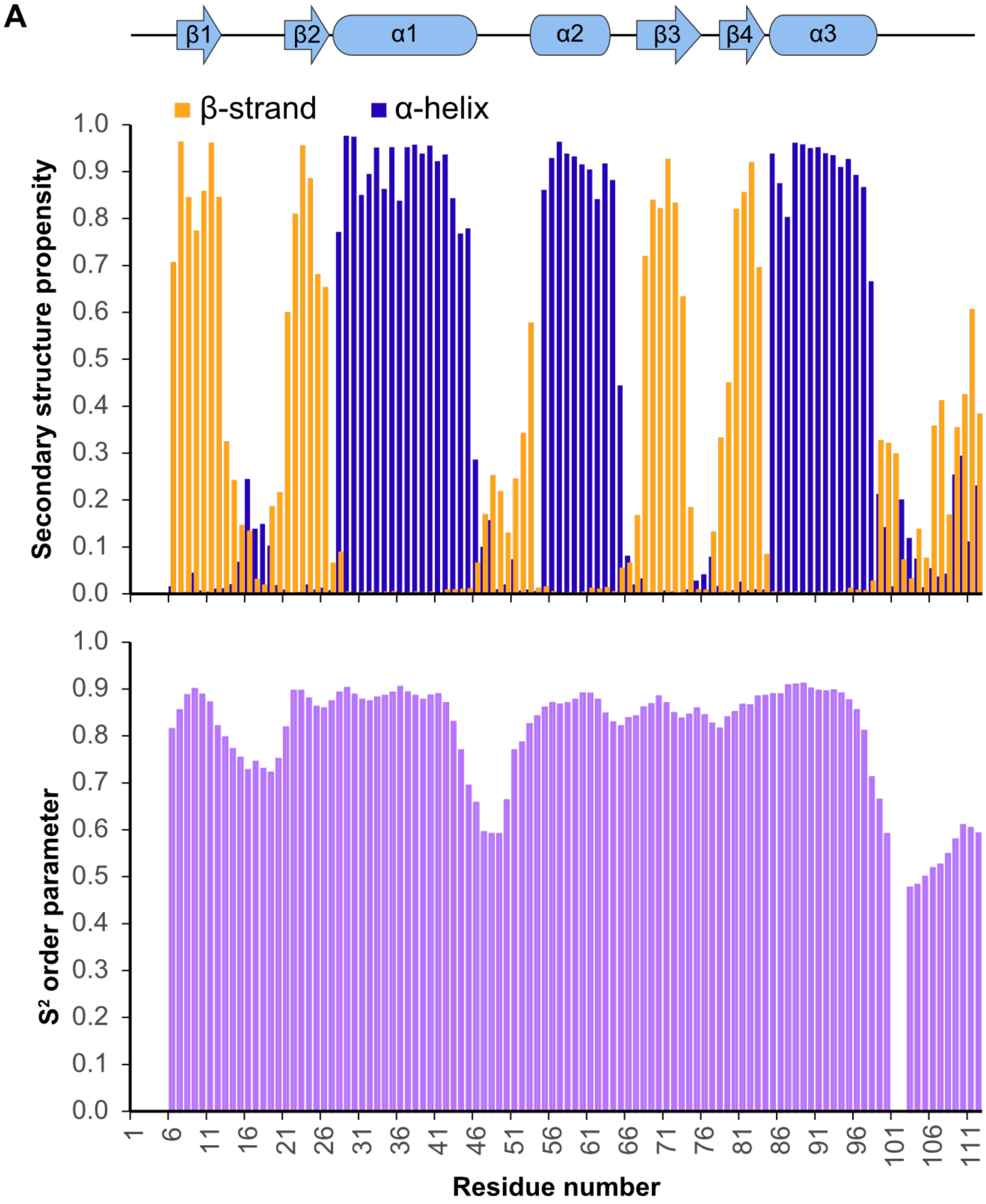

Supplemental Figure S5 continued

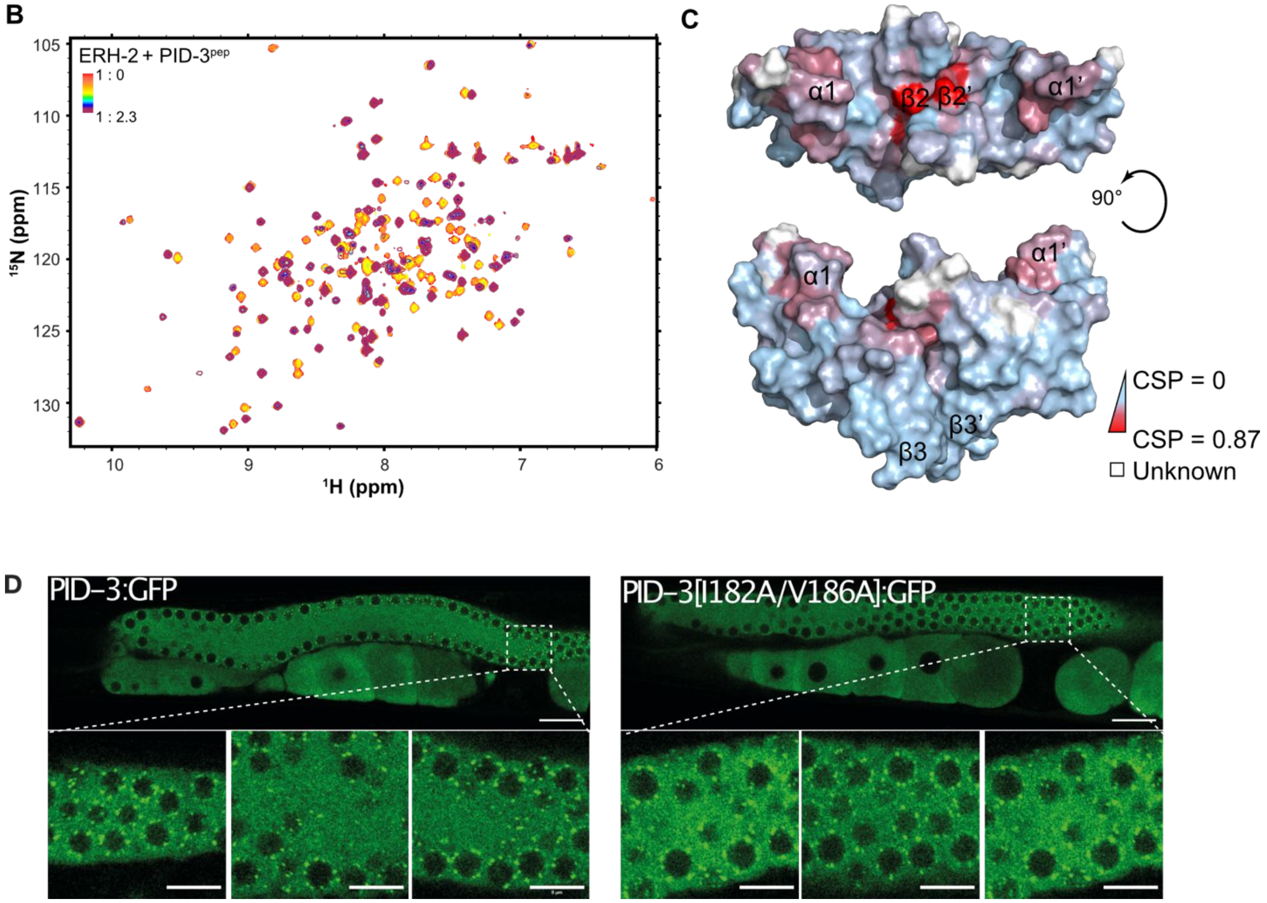


**Supplemental Figure S5: Structural analysis of full-length ERH-2 by NMR** (A) Secondary structure prediction and flexibility of ERH-2. The backbone chemical shifts of free full-length ERH-2 ($^1H^N$, $^{15}N$, $^{13}C^\alpha$, and $^{13}C^\beta$) were used as input for the MICS program to calculate secondary structure propensities (above), together with the $S^2$ order parameter as a function of residue (below, see Methods). The predicted β-strands and $\alpha$-helices found in solution are colored orange and blue, respectively, and are fully consistent with the crystal structures of the ERH domain. In addition, the data shows increased flexibility in the loops connecting helices $\alpha$1 and $\alpha$2, strands β1 and β2, as well as the C-terminal ~10 residues, highlighted also by the lower $S^2$ order parameters. (B) $^1H$-$^{15}N$ HSQC titrations of $^{15}N$-labeled ERH-2 with PID-3$^{pep}$. Incremental amounts of the peptide concentrated to ~ 1.5 mM were added to ~200 μM of $^{15}N$-labeled ERH-2, and the changes monitored at each step. The resulting spectra are overlaid, beginning with free ERH-2 colored in red, and ending with the ERH-2/peptide complex in maroon (1:2.3 molar ratio). (C) The CSP values are mapped onto the structure of the dimeric ERH-2 (PDB: 7O6N), with the regions most affected by binding colored in increasing gradations of red. Unassigned residues, prolines, or amino acids which we cannot track in the bound state are colored white. (D) Single-plane confocal micrographs of PID-3::GFP(WT) and PID-3[I182A; V186A]::GFP at 25°C. The boxes indicate the regions (above the spermatheca) from which three zoomed-in examples are given below. Scale bars: 20 µm in overview, 8 µm in zoom-in.

Supplemental Figure S6

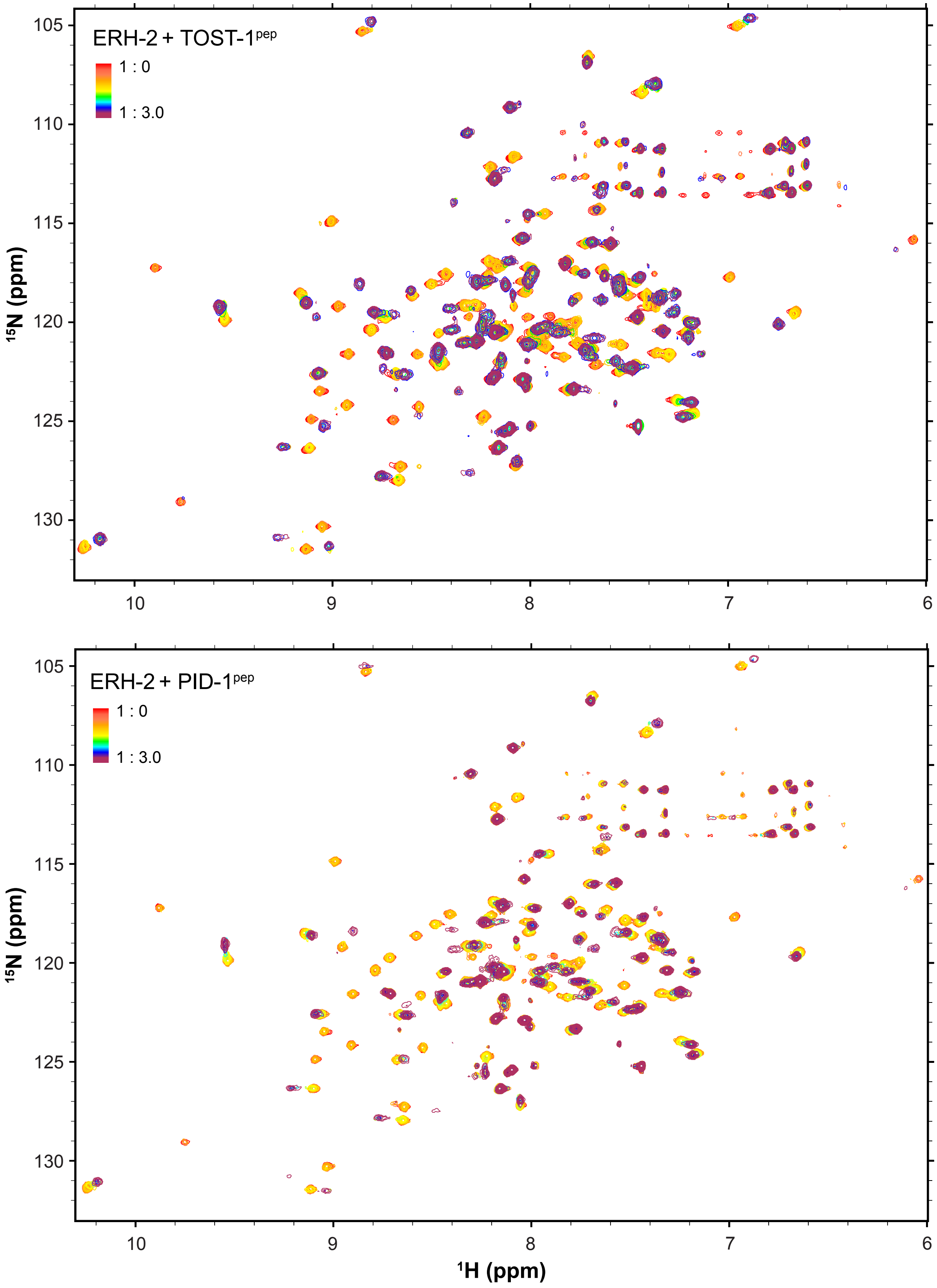

**Supplemental Figure S6:** $^{1}H$-$^{15}N$ HSQC titrations of $^{15}N$-labeled ERH-2 with TOST-1$^{pep}$ (above) and PID-1$^{pep}$ (below). In each case, incremental amounts of the peptide concentrated to ~ 1.5 mM were added to ~200 μM of $^{2}H$/$^{15}N$/$^{13}C$-labeled ERH-2, and the changes monitored at each step. The resulting spectra are overlaid, beginning with free ERH-2 colored in red, and ending with the ERH-2/peptide complex in maroon (1:3 molar ratio for both).

#### Supplemental Figure S7

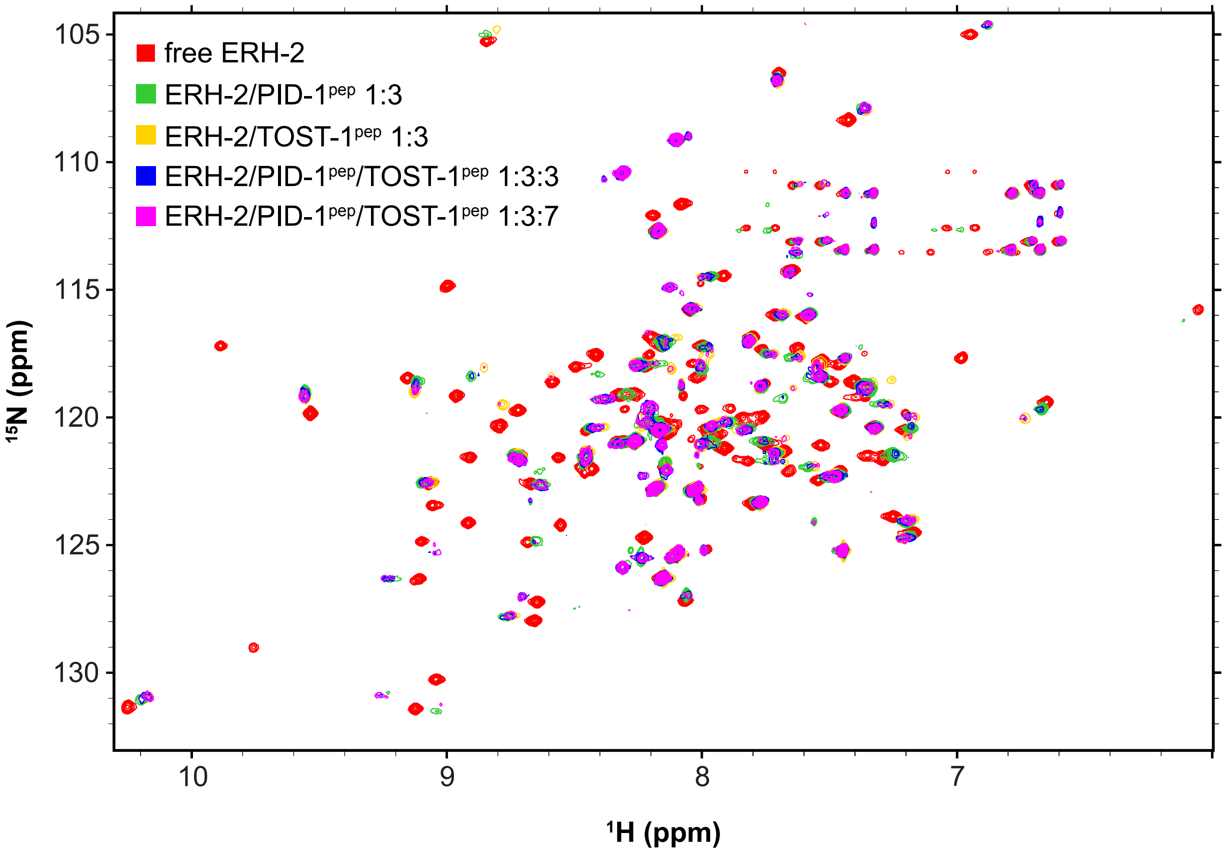


**Supplemental Figure S7.** Interplay of PID-1pep and TOST-1pep binding to ERH-2. Shown is an overlay of $^{1}H$-$^{15}N$ HSQCs of ERH-2 in various states, including unbound (red), in the presence of excess PID-1pep (green), excess TOST-1pep (yellow), as well as ERH-2 in the presence of equimolar amounts of both peptides (blue), or with excess TOST-1pep over PID-1pep (magenta). The molar ratios are indicated in the figure legend. When TOST-1pep is in excess relative to PID-1pep, we see chemical shifts consistent with the formation of an ERH-2/TOST-1pep complex, while in the presence of equal amounts of both peptides, ERH-2 exhibits intermediate chemical shifts which indicates contribution from both binding events.

Supplemental Figure S8

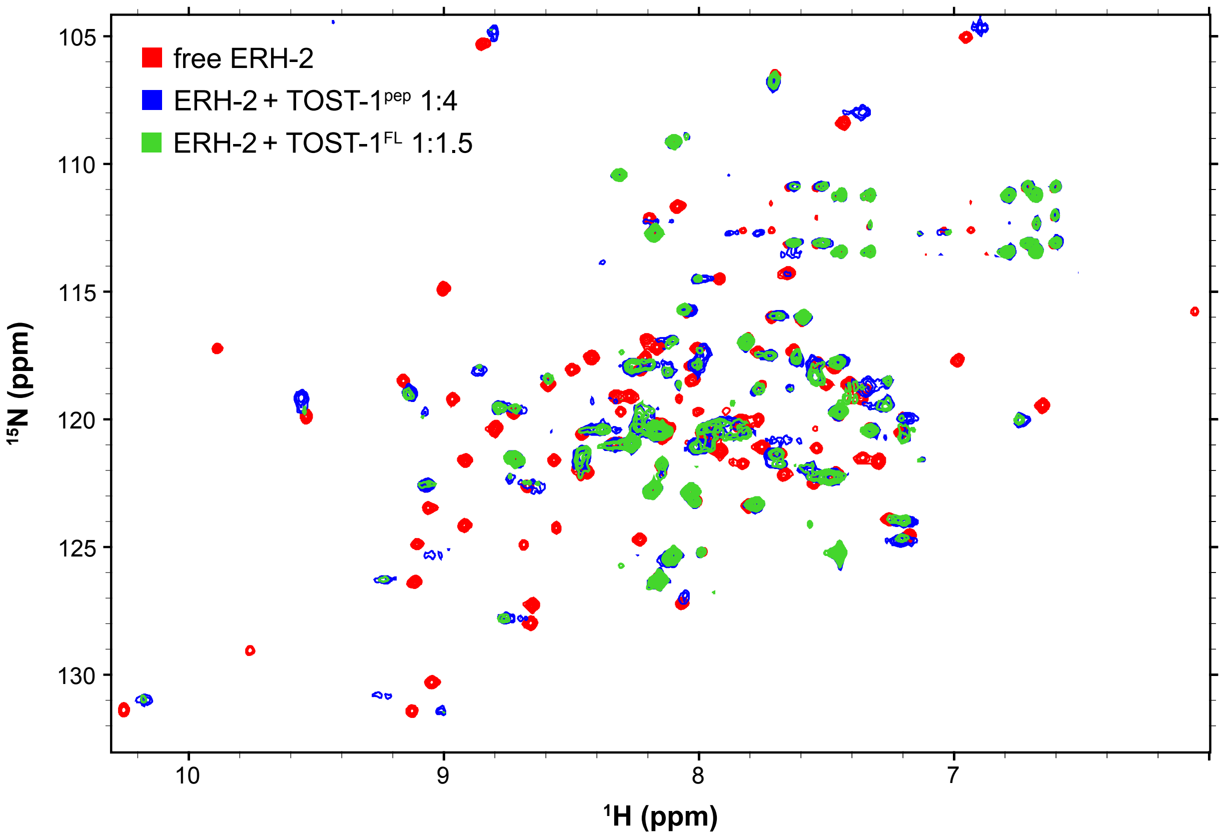


**Supplemental Figure S8.** Binding of ERH-2 by TOST-1$^{pep}$ is very similar to that of TOST-1$^{FL}$. Overlaid $^{1}$H-$^{15}$N HSQC spectra of $^{2}$H/$^{15}$N/$^{13}$C-labeled ERH-2 in the presence of 4-fold molar excess TOST-1$^{pep}$ (blue), or 1.5-fold molar excess of TOST-1$^{FL}$ (green). Although many amide peaks are not visible in the larger ERH-2/TOST-1$^{FL}$ complex, those that we can detect overlap well between the two TOST-1 molecules, indicating that the shorter TOST-1 peptide largely recapitulates full-length TOST-1 binding.

### Supplemental Figure S9

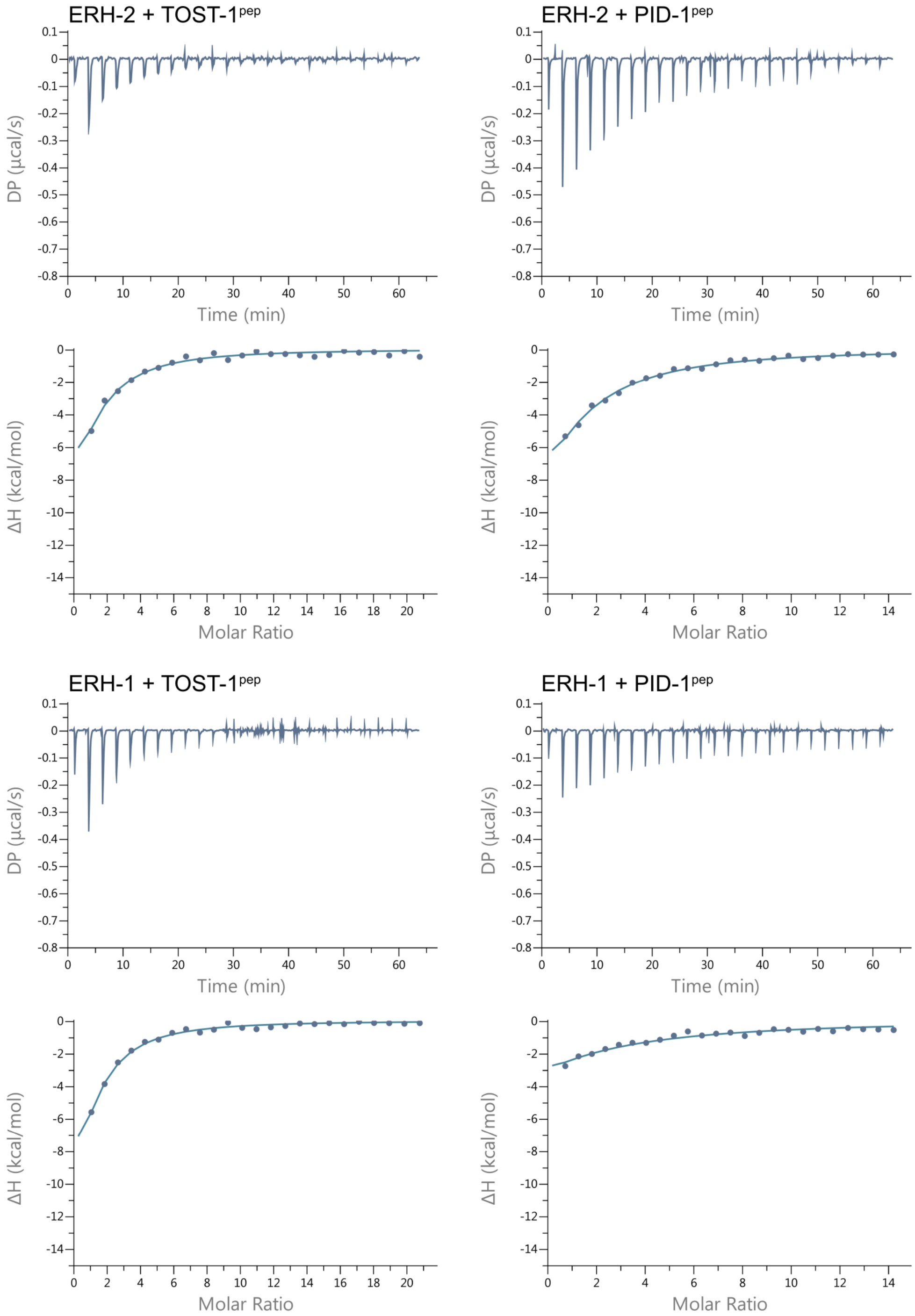

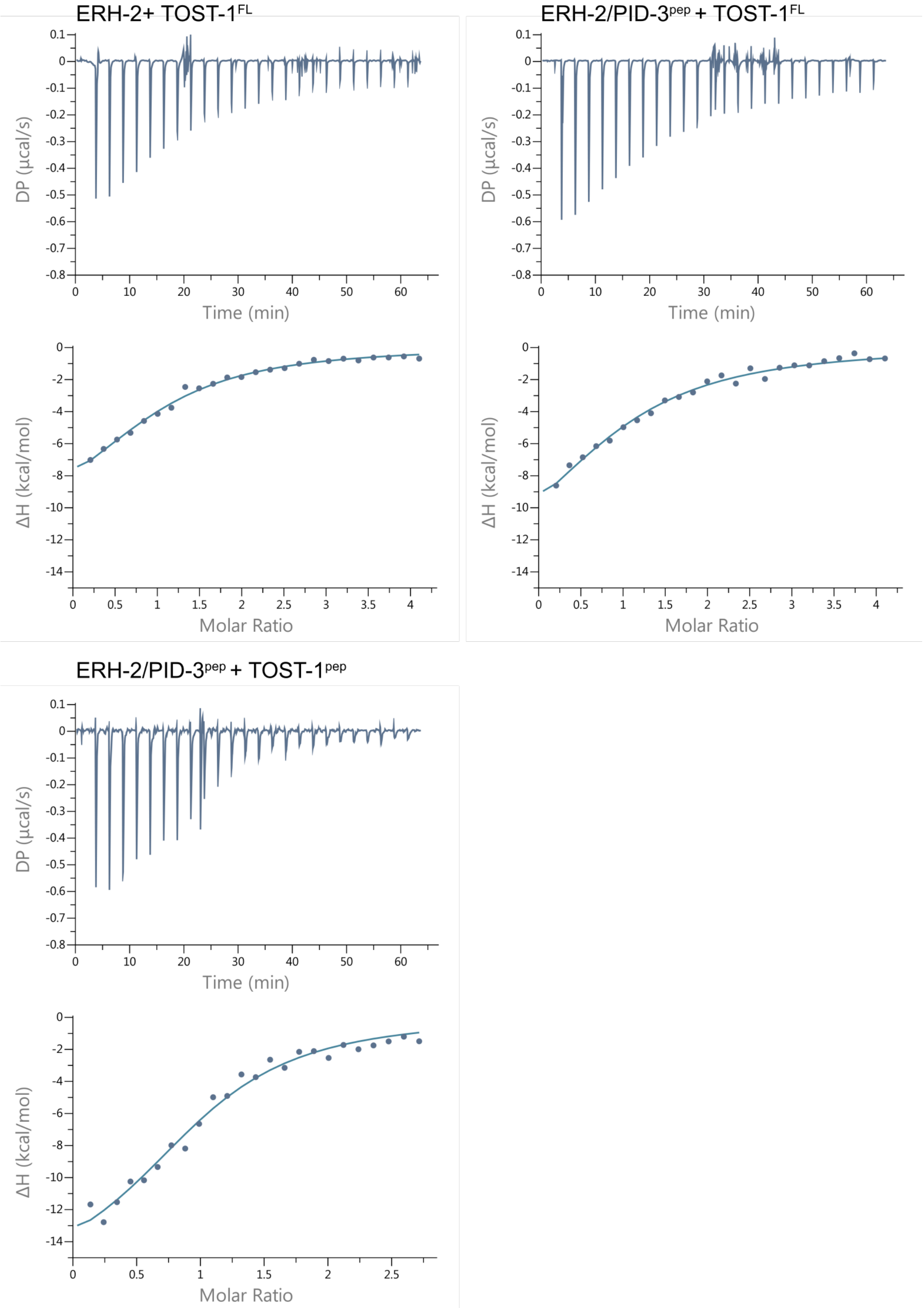


**Supplemental Figure S9.** Analysis of the interaction of ERH- with TOST-1$^{FL}$, TOST-1$^{pep}$, and PID-3$^{pep}$ by ITC. The baseline-corrected data (top) were integrated and fitted to a simple 1:1 binding isotherm (bottom). One representative binding isotherm is shown in each case. The average thermodynamic parameters calculated from two or three independent experiments is shown in Table 1.

#### Supplemental Figure S10

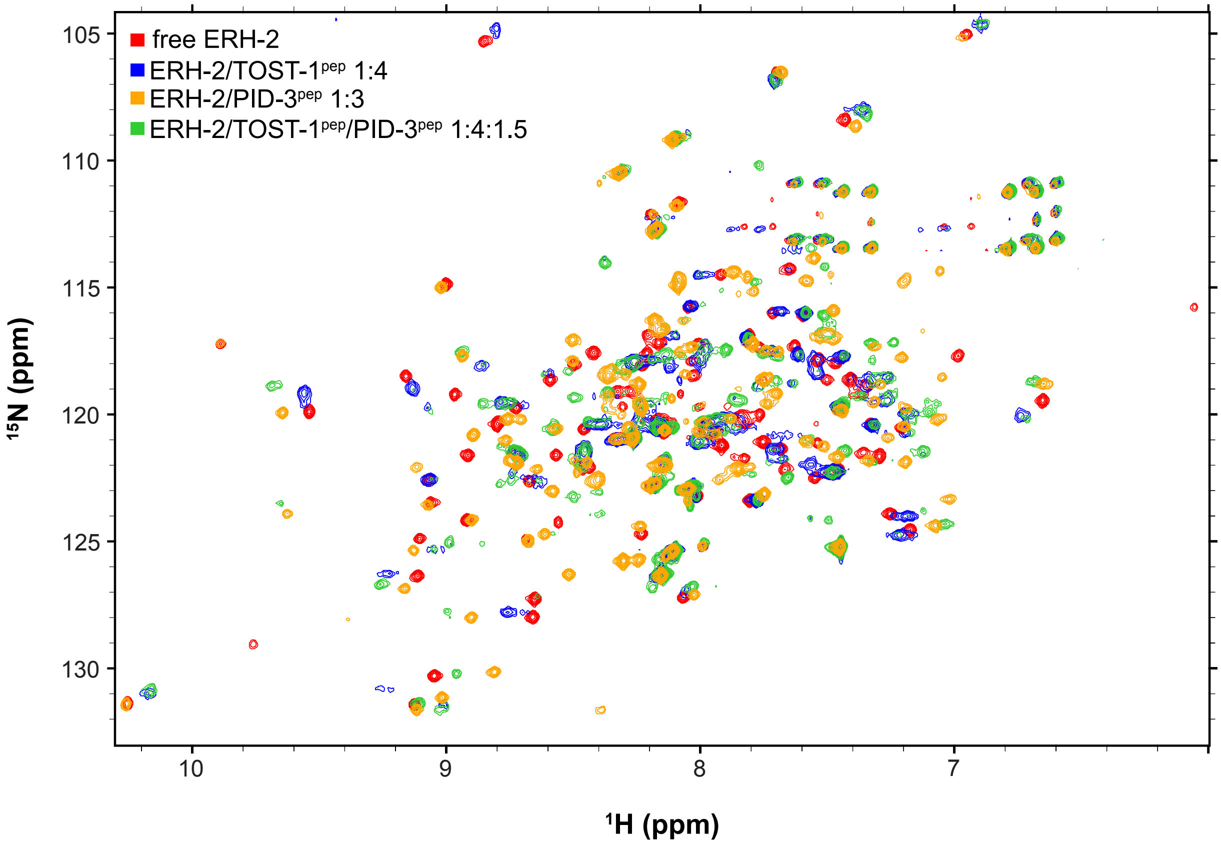


**Supplemental Figure S10.** Interplay of PID-3pep and TOST-1pep binding to ERH-2. Shown is an overlay of $^{1}H$-$^{15}N$ HSQCs of ERH-2 in various states, including unbound (red), in the presence of excess TOST-1pep (blue), excess PID-3pep (orange), and in the presence of both peptides (green). The molar ratios are indicated in the figure legend. In the presence of both peptides, we detect new peaks not observed in the isolated subcomplexes, indicating the formation of a new ternary structure. As well, characteristic TOST-1pep-bound peaks appear stronger, suggesting that PID-3pep may stabilize binding of ERH-2 to TOST-1pep.

#### Supplemental Figure S11

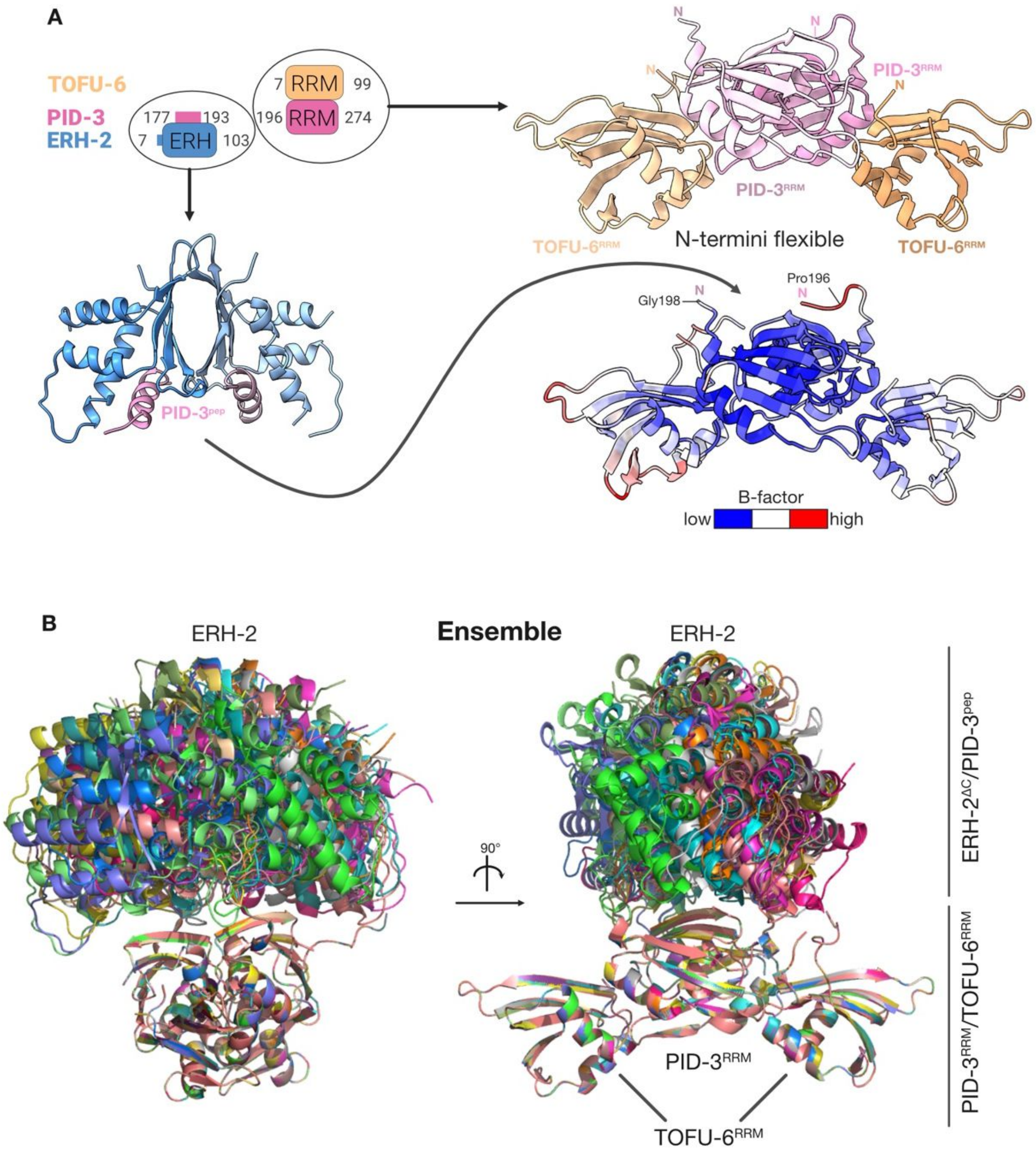


**Supplemental Figure S11.** Modelling of the PETISCO core complex. (A) Strategy for core complex modelling. In the ERH-2$^{\Delta C}$/PID-3$^{pep}$ subcomplex (bottom left) PID-3$^{pep}$ contains residues 177-193 and in the PID-3$^{RRM}$/TOFU-6$^{RRM}$ subcomplex (top right) the PID-3$^{RRM}$ domain contains residues 196-274. The crystal structure of the PID-3$^{RRM}$/TOFU-6$^{RRM}$ is shown on the top right in the same colors as in Fig. 2. On bottom right the structure is colored according to the B factor. Blue indicates low, white intermediate and red high B factors. (B) Ensemble of 20 different models superimposed on the PID-3$^{RRM}$/TOFU-6$^{RRM}$ core of the template structure. Each structure is shown in a different color.

### Supplemental Figure S12

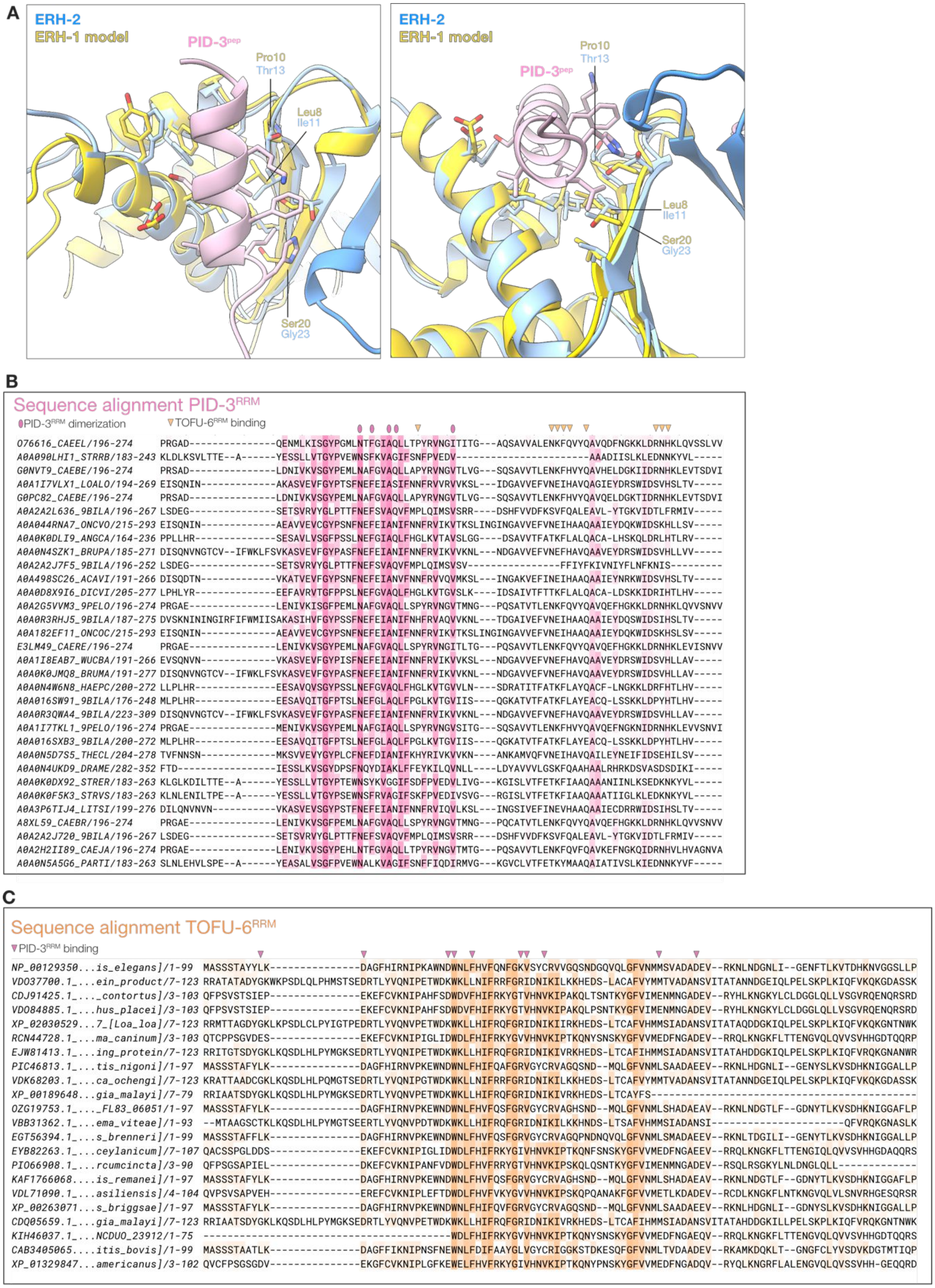

**Supplemental Figure S12.** Evolutionary analysis. (A) A homology model of ERH-1 was generated using I-TASSER with the ERH-2 structure as a template. ERH-1 was then superimposed onto the ERH-2$^{\Delta C}$/PID-3$^{pep}$ complex. Several differences between ERH-1 and ERH-2 at the PID-3$^{pep}$ binding interface are highlighted. (B) Multiple sequence alignment of PID-3$^{RRM}$ from different nematodes. Highly conserved residues are shown in dark pink, less conserved residues are in light pink. Above the sequence, residues involved in the homodimerization of the PID-3$^{RRM}$ are labeled with a pink oval, while those that interact with the TOFU-6$^{RRM}$ are labeled with an orange triangle. The accession number of the nematode PID-3 protein sequences are shown on the left. (C) Multiple sequence alignment of TOFU-6$^{RRM}$ from different nematodes. Highly conserved residues are shown in dark orange and less conserved residues are in light orange. Above the sequence, the residues of the TOFU-6$^{RRM}$ that interact with the PID-3$^{RRM}$ are labeled with a pink triangle. The accession number of the nematode TOFU-6 protein sequences are shown on the left.
