## Supplementary material for "Structural basis of PETISCO complex assembly during piRNA biogenesis in *C. elegans*": Table S1

### Supplementary Table S1: Crystal Structures data collection and refinement statistics

R.m.s.d., Root mean square deviation

| Dataset | ERH-2ΔC | ERH-2 ΔC/PID-3$^{pep}$ | PID-3$^{RRM}$ | PID-3$^{RRM}$/TOFU-6$^{RRM}$ | |
|---|---|---|---|---|---|
| PDB | 7O6L | 7O6N | 7OCX | | 7OCZ |
| Space group | P 1 $2_1$ 1 | P $2_1$ $2_1$ $2_1$ | P$6_5$ | P $2_1$ $2_1$ $2_1$ | P $2_1$ $2_1$ $2_1$ |
| Cell dimensions | | | | | |
| *a, b, c* (Å) | 38.27, 54.05, 46.76 | 44.99, 52.95, 125.61 | 43.29, 43.29, 158.18 | 43.88, 68.47, 130.79 | 43.98, 68.00, 130.59 |
| $\alpha, \beta, \gamma$ (°) | 90.0, 93.4, 90.0 | 90.0, 90.0, 90.0 | 90.0, 90.0, 120.0 | 90.0, 90.0, 90.0 | 90.0, 90.0, 90.0 |
| **Data Collection** | | | | phasing | native |
| Wavelength (Å) | 0.87 | 0.97 | 1.28 | 1.54 | 1.00 |
| Resolution range | 35.35 - 1.50 (1.54 - 1.50) | 48.79 - 2.17 (2.25 - 2.17) | 36.48 - 1.82 (1.89 - 1.82) | 68.53 – 1.93 (1.93 – 1.89) | 41.68 - 1.70 (1.76 - 1.70) |
| No. of reflections | 224,226 | 210,614 | 256,407 | 348,668 | 552,651 |
| No. of unique reflection | 30,574 | 16,347 | 14,314 | 29,377 | 43,394 |
| $R_{merge}$ (%) | 6.2 (160.8) | 12.3 (209.1) | 22.0 (160.2) | 11.3 (169.0) | 8.6 (352.7) |
| $R_{pim}$ (%) | 2.4 (62.9) | 3.6 (59.9) | 5.2 (50.1) | 3.4 (64.5) | 2.5 (119.0) |
| $I / \sigma I$ | 10.0 (1.1) | 13.2 (1.1) | 38.8 (0.6) | 14.2 (1.7) | 16.2 (0.56) |
| Completeness (%) | 99.8 (99.5) | 98.9 (90.0) | 95.0 (68.32) | 90.8 (51.6) | 97.9 (80.9) |
| Multiplicity | 7.3 (7.4) | 12.9 (12.6) | 17.9 (10.5) | 11.9 (7.4) | 12.7 (9.3) |
| $CC_{1/2}$ | 1 (0.56) | 1 (0.84) | 0.99 (0.44) | 0.99 (0.38) | 1 (0.17) |
| **Refinement** | | | | | |
| Resolution range | 31.2–1.50 (1.55–1.50) | 48.79 - 2.17 (2.25 - 2.17) | 36.48 - 1.82 (1.89 - 1.82) | | 41.68 - 1.70 (1.76 - 1.70) |
| Reflections used in refinement | 30,535 (3029) | 16,337 (1455) | 14,244 (1031) | | 43,052 (3501) |
| $R_{work}$ / $R_{free}$ (%) | 20.0 / 21.6 | 20.1 / 24.5 | 18.2 / 21.0 | | 19.4 / 23.1 |
| Wilson B-factor (Å$^2$) | 25.68 | 59.32 | 24.56 | | 33.62 |
| Average B-factors (Å$^2$) | 23.93 | 76.55 | 31.77 | | 46.29 |
| No. of atoms | | | | | |
| Proteins | 1433 | 1735 | 1232 | | 2579 |
| Ligands | 1 | 3 | 2 | | 0 |
| Solvent | 102 | 27 | 106 | | 139 |
| **Stereochemistry** | | | | | |
| R.m.s.d. bond lengths (Å) | 0.010 | 0.003 | 0.005 | | 0.009 |
| R.m.s.d. bond angles (°) | 1.05 | 0.66 | 0.85 | | 1.10 |
| Ramachandran favored (%) | 99.4 | 99.1 | 98.1 | | 98.2 |
| Ramachandran outliers (%) | 0 | 0 | 0 | | 0 |
